# Mapping disease critical spatially variable gene programs by integrating spatial transcriptomics with human genetics

**DOI:** 10.1101/2025.09.24.678397

**Authors:** Hanbyul Lee, Haochen Sun, Xuewei Cao, Berke Karaahmet, Zhijian Li, Hans-Ulrich Klein, Mariko Taga, Gao Wang, Philip L. De Jager, David A. Bennett, Luca Pinello, Xin Jin, Rahul Mazumder, Kushal K. Dey

## Abstract

Spatial gene expression patterns underlie tissue organization, development, and disease, yet current methods for detecting spatially variable genes (SVGs) lack the flexibility to capture multi-scale structure, ensure robustness across platforms, and integrate with genetic data to assess disease relevance. We present Spacelink, a unified framework that models spatial variability of a gene at both whole-tissue and cell-type resolution using an adaptive mixture of data-driven spatial kernels and summarizes it using an Effective Spatial Variability (ESV) metric. Spacelink achieved up to 3.2x higher detection power over eight existing global SVG and cell-type SVG methods while showing consistently superior FDR control across 34 different simulation settings and also showed superior cross-platform concordance in matched tissue Visium and CosMx datasets. Applied to 3 healthy CosMx human tissues (brain cortex, lymph node, liver), Spacelink revealed that SVGs are highly informative for 113 complex traits and diseases (average GWAS sample size = 340,406). Spacelink showed up to 2.2x higher disease informativeness over competing methods in tissue-relevant complex diseases and traits, conditional on putative non-spatial expression-level confounders. Applied to a mouse organogenesis Stereo-seq atlas (8 developmental stages), Spacelink identified 145 genes with stage-associated ESV within brain independent of mean expression, that are enriched in pathways like Wnt signaling and Rap1 signaling characterizing early and late development, respectively. Integration with in vivo Perturb-seq targeting 35 de novo ASD risk genes revealed that perturbations in excitatory neurons and astrocytes preferentially altered spatially structured downstream gene programs (1.7–2.2x higher average ESV across stages than other cell types), many of which were enriched for polygenic autism GWAS loci. In neurodegeneration, analysis of 32 Visium dorsolateral prefrontal cortex samples spanning Alzheimer’s disease (AD) pathology stages identified 334 genes with decreasing ESV along amyloid burden (enriched for glycolysis) and 216 genes with decreasing ESV along tau tangle accumulation (enriched for apoptotic pathways). Several AD risk genes (*PKM, CLU, GPI*) showed conserved reductions in spatial variability with AD pathology in both human and 5xFAD mouse, with *PKM* linking to a colocalized splicing QTL and amyloid burden QTL variant. These results highlight the utility of Spacelink in decoding spatially variable gene programs that connect tissue architecture to disease genetics.

## Introduction

Spatially resolved transcriptomics (SRT) enable measurement of gene expression at cellular or near-cellular resolution while retaining spatial coordinates, thereby providing an unprecedented opportunity to map spatially variable genes (SVGs) that capture localized regulatory programs and tissue organization^1–3^. Linking this spatial variability to human disease genetics holds promise for uncovering tissue microenvironments and cellular niches through which genetic risk factors act, providing new insights into disease mechanisms and informing therapeutic interventions^4–7^. Despite their potential, current approaches for detecting SVGs face several limitations that restrict their utility for disease genetics. Standard global SVG methods – nnSVG^8^, SpatialDE^9^, SpatialDE2^10^, SPARK^11^ - typically assume a single spatial scale or a limited set of fixed kernels, and thus fail to capture the gene-specific multi-scale organization of expression that arises from layered cell type distributions, fine-grained niches, and broad domains within tissues. Recent cell-type SVG methods (STANCE^12^, Celina^13^) can capture finer changes in spatial variability within a cell type, however, we highlight that they often fail to appropriately correct for spatial colocalization with other cell types, leading to inflated false positives or reduced power. It is also unclear which metric of spatial variability would be most appropriate for comparing genes in a manner that is stable across platforms^14^, experimental conditions^15,16^ and physiological contexts^17–19^ —an essential requirement for integrating SVG programs with human genetics to robustly assess disease relevance.

Here, we present *Spacelink*, a unified statistical framework for detecting and prioritizing SVGs at both global tissue and cell-type resolution. Spacelink employs an adaptive multi-kernel model to capture spatial variance across diverse length scales, and its cell-type-specific version introduces a data-driven gating strategy to correct for spatial colocalization, designed to improve the specificity for cell types that are weakly represented in mixed spots relative to more abundant colocalizing cell types. To summarize spatial variability, we define *Effective Spatial Variability (ESV)*, a metric which integrates variance magnitude of each component kernel and its corresponding spatial scale into a single interpretable score directly suited for genetic analyses. We extensively benchmarked the global and cell-type versions of Spacelink against 8 state-of-the-art global and cell-type SVG methods across 34 diverse simulation settings informed by both real-world spatial datasets and theoretically distinct spatial patterns; Spacelink consistently outperforms existing methods in power, robustness, and reproducibility. Applications to spatial data from multiple healthy tissue atlases, mouse organogenesis, and Alzheimer’s disease neurodegeneration demonstrate Spacelink’s ability to integrate spatial variability with human genetics, revealing disease-relevant programs that are not captured by conventional expression-based analyses.

## Results

### Overview of Methods

In Spacelink, we propose a framework to identify and prioritize spatially variable genes using spatially resolved transcriptomics data, both at the resolution of the whole tissue and individual cell types, and evaluate their informativeness in human disease, organogenesis, and tissue degeneration. Spacelink employs multiple data-driven spatial kernels to model the data covariance of each gene (**Figure 1a**). Spacelink models normalized and transformed expression values of a gene g at *N* spatial locations **s** = (**s**_1_, **s**_2_, …, **s**_*N*_) as follows.

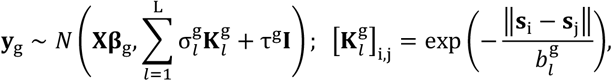

where **X** denotes the covariates including the intercept and other spot-level metadata if available, 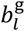 represents kernel bandwidth (spatial length scale), 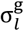 denotes the variance component attributed to each exponential kernel 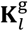, and 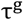 captures residual noise. By combining kernels with different length scales, Spacelink accommodates spatial variability ranging from fine-scale microenvironments to broad tissue domains. The set of spatial kernels 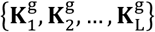 can be viewed as a data-driven selection from a potentially infinitely large collection of spatial kernels of varying bandwidths, which we term as a “*Kernel Store*” (**Methods, Supplementary Figure S1a**). To test whether a gene *g* is spatially variable, Spacelink evaluates the null hypothesis 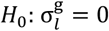 for all *l*, against the alternative 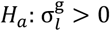 for some *l*. To this end, we utilize the score test statistic for each kernel which follows a mixture of chi-square distributions^20^. By applying the Satterthwaite method, we approximate the p-value for each statistic and then combine these p-values using the Cauchy combination rule^21^ to yield a single, aggregated p-value (**Methods, Supplementary Note**). We also estimate variance components 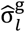 using a Non-Negative Least Squares (NNLS) approach^22^, which links the empirical covariance of each gene to the pre-selected spatial kernels. Beyond hypothesis testing, we introduce a novel gene-level metric, Effective Spatial Variability (ESV), which integrates estimated variance components with their corresponding kernel bandwidths to provide an interpretable score for ranking genes by spatial variability and facilitating genetic integration.

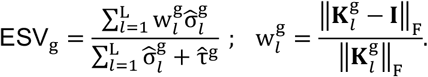

**Figure 1.**
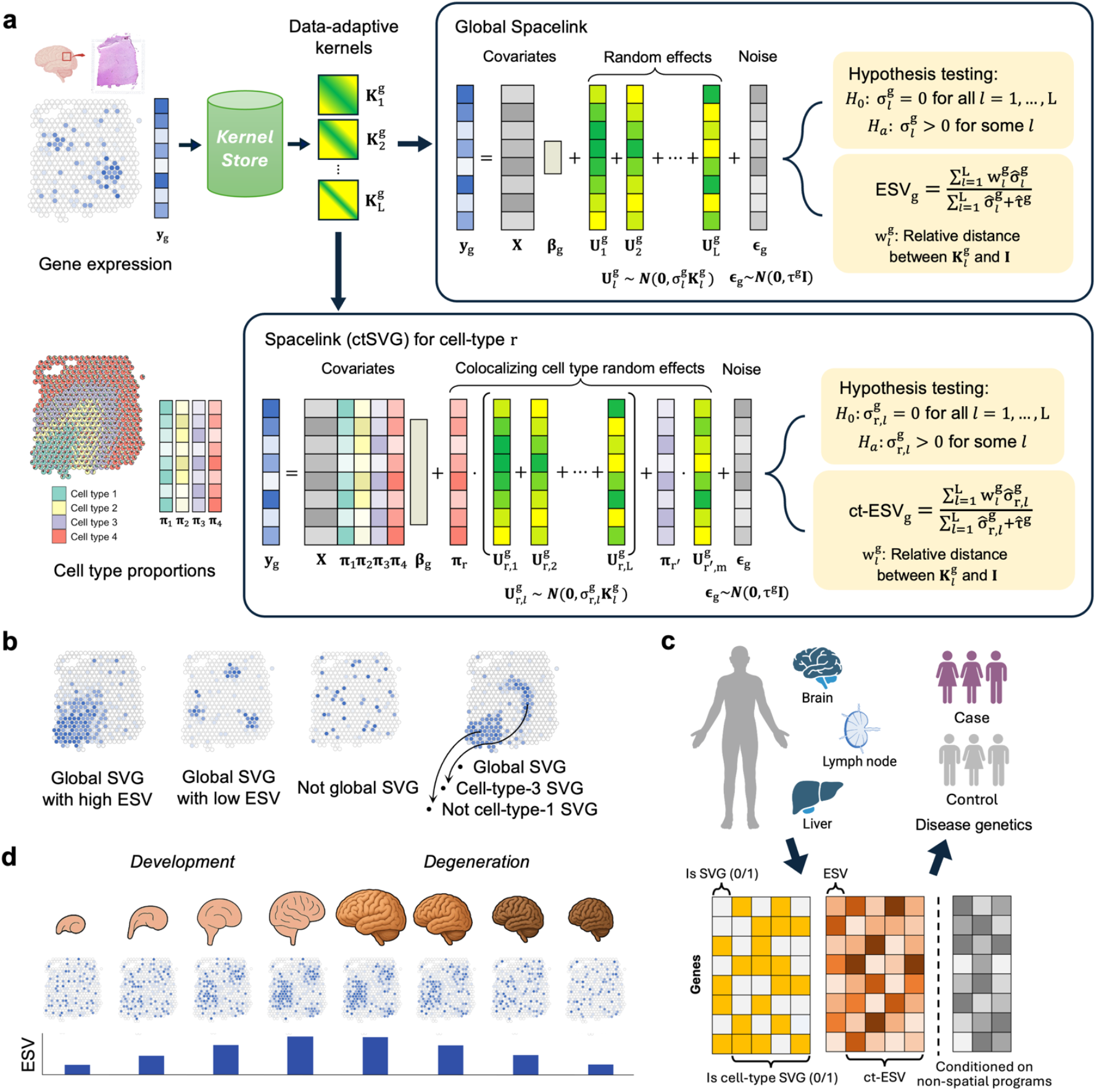
Illustration of the Spacelink framework for identifying spatially variable genes (SVGs) at both global and cell-type levels, and its disease-focused applications. (a). Spacelink selects data-adaptive kernels from a “Kernel Store”—a collection of exponential kernels spanning multiple bandwidths—and models gene expression using a linear mixed model, where total spatial variance is represented as a mixture of kernel-specific variance components. At the global level, SVGs are identified by testing whether all variance components are zero, and spatial variability is quantified using the Effective Spatial Variability (ESV) metric. At the cell-type level, Spacelink extends the model for lower-resolution platforms (e.g., 10x Visium) by (i) incorporating estimated cell type proportions as fixed-effect covariates, (ii) projecting kernel random effects to the focal cell type, and (iii) conditioning out the spatial effects of colocalizing cell types via a data-driven gating procedure. Hypothesis testing and ct-ESV calculation follow the same principles as the global model. (b). Illustrative examples of genes exhibiting diverse global and cell-type level spatial expression patterns. From left to right: (i) a global SVG with high spatial bandwidth of variation resulting in high ESV, (ii) a global SVG showing multiple clusters of signal with low spatial bandwidth of signal, thereby resulting in low ESV, (iii) a non-SVG with random, spotty expression, (iv) a global SVG that is also a ct-SVG in cell type 3, but is not a ct-SVG for cell type 1, based on the cell type distribution in Panel a. (c) Spacelink is applied to SRT data from 3 tissues (brain cortex, lymph node, liver) with a large number of diseases and quantitative traits with well-powered Genome-wide Association Studies (GWAS). We evaluate gene programs based on Spacelink hypothesis test, ESV and ct-ESV in terms of disease informativeness in both matched and non-matched diseases and traits, conditional on putative non spatial confounders. (d). Spacelink is applied to spatiotemporal data along mouse organogenesis and Alzheimer’s disease (AD) human and mouse neurodegeneration data to identify genes whose spatial variability is associated with developmental and degenerative processes.

ESV is bounded between 0 and 1 and is determined by two key factors—the spatial length scale *l* and the spatial scale-specific signal-to-noise ratio (SNR) 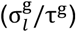 —and it increases with larger values of either parameter (**Supplementary Figure S1c**). In comparison, other analogous scores like PropSV (nnSVG^8^) and FSV (SpatialDE^9^) are largely insensitive to changes in length scale and vary largely with SNR (**Supplementary Figure S1c**). The weighting scheme of ESV penalizes kernels with minimal spatial structure, prioritizing genes with strong, non-trivial spatial variability (**Figure 1b**). Genes that do not reach statistical significance in the Spacelink hypothesis test are assigned an ESV score of 0. By construction, ESV downweighs trivial, very short-range autocorrelations that may arise from technical artifacts such as transcript spillover or cell segmentation errors, thereby prioritizing genes with robust spatial organization at biologically meaningful length scales. Further details on the ESV formulation are provided in the **Methods** section.

In lower-resolution SRT platforms such as 10x Visium, where each spot aggregates transcripts from multiple cells spanning different types, it is difficult to resolve spatial variability specific to a single cell type. For example, a weakly representative cell type, like microglia in brain cortex, may colocalize with other dominant cell types, like oligodendrocytes within mixed spots, leading to confounded signals. To this end, we propose a cell-type-specific extension of Spacelink that attributes spatial variability to a focal cell type r, while correcting for potential confounding due to spatial colocalization with other cell types (**Figure 1a**). For a specific cell type r, we model the expression data **y**_g_ as follows.

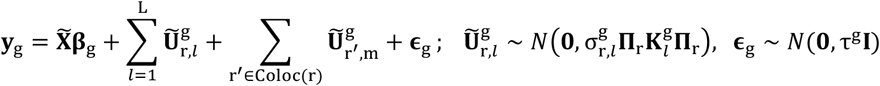

where 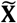 denotes fixed effect covariates comprising the intercept, estimated cell type proportions in spots as evaluated by RCTD method^23^, as well as other spot-level metadata if available. **Π**. represents a *N* × *N* diagonal matrix of the proportions of cell type r across *N* spots. 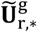 represent random effects capturing the projection of the pre-selected kernels for gene g in the global SVG model to the focal cell type r. A key technical challenge in cell-type SVG modeling is in disentangling the spatial random effects of the focal cell type from that of other colocalizing cell types. This colocalization may artificially introduce correlation among the random effects, and not correcting for it may lead to false positives. On the other hand, naively including random effects for all cell types and length scales can lead to an overcrowding of kernels, where the model becomes overparameterized, diluting the power to detect true spatial signals in the focal cell type. To resolve this, the cell-type-specific version of Spacelink employs a data-driven gating procedure to determine a set of cell types, Coloc(r), that are spatially colocalized with the focal cell type r (see **Methods** for details) and then corrects for the random effects of each of these cell types at the median bandwidth for gene g. We leverage this model to test whether a gene is spatially variable within a cell type 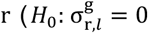 for all *l*, r is fixed ) and use Restricted Maximum Likelihood (REML) approach^24^ to generate estimates of 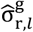 . Analogous to *ESV*, we provide cell-type-specific *ct-ESV* scores, by replacing 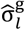 with 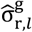 (see **Methods** for additional details). Genes that do not reach statistical significance in the cell-type-specific Spacelink hypothesis test are assigned a ct-ESV score of 0. The ESV and ct-ESV scores are used as primary metrics to link spatial variability of genes to human disease. We also benchmark ESV against other available spatial metrics, such as PropSV^8^, FSV^9^, model LLR (log-likelihood ratio)^8,9^ and p-values from SVG hypothesis tests, by evaluating their ability to predict disease-associated genes relevant to the underlying tissue.

Spacelink offers several key advantages over existing methods for detecting spatially variable genes (SVGs). First, it provides a unified kernel-based framework for identifying SVGs at both the whole-tissue level and in a cell-type-specific manner (**Figure 1a**) — a capability that, outside of STANCE^12^, is largely missing from other tools. Spacelink selects multiple kernels in a data-driven manner for each gene, which (i) reduces reliance on a limited set of pre-defined kernels as used in SPARK^11^ and SpatialDE/SpatialDE2^9,10^, and (ii) enables accurate modeling of genes with heterogeneous spatial scales due to multi-scale tissue architecture — a limitation in methods employing a single gene-level spatial length scale such as nnSVG^8^. Second, as highlighted below, Spacelink ESV demonstrates superior transferability across datasets and platforms as well as higher disease informativeness over existing spatial metrics, making it better suited for cross-sample comparisons and downstream phenotypic association. Third, the data-driven random effect gating in the cell type version of Spacelink is designed to improve the specificity and sensitivity of the model compared to no cell type colocalization correction (as in Celina^13^) and full random effect adjustment across all cell types (as in STANCE^12^) respectively. Fourth, Spacelink extends the ESV framework to the cell-type level via the ct-ESV score, enabling quantification and prioritization of cell type SVGs; Celina and STANCE do not provide a comparable quantitative metric of cell-type-specific spatial variability. Finally, Spacelink is computationally efficient and scalable to tens of thousands of genes and thousands of spatial locations. It uses closed-form updates for NNLS-based variance component estimation and supports parallelization across genes, enabling rapid inference at large scale. Furthermore, we provide *Spacelink-lite* for larger datasets with hundreds of thousands of spatial locations, which produces results nearly identical to the original Spacelink while being significantly faster and more memory-efficient. Spacelink-lite achieves this through a two-scale distance approximation: spots are first aggregated into a grid, and kernel distances are computed at spot resolution for neighboring locations and at grid resolution for more distant locations. This preserves fine-scale local spatial structure while substantially reducing the computational cost of modeling long-range spatial dependencies, retaining the ability to capture both local and global spatial variability (see **Methods** for details).

We applied Spacelink to a broad spectrum of spatial transcriptomics datasets from human and mouse, including (i) 3 high single-cell resolution CosMx samples (see **Data Availability**) and matched lower-resolution 10x Visium samples from brain cortex, lymph node and liver^25,26^ (**Data Availability**), (ii) mouse organogenesis Stereo-seq data spanning 8 developmental stages (E9.5 to E16.5)^27^, and (iii) 32 ROSMAP 10x Visium human samples spanning different stages of Alzheimer’s disease pathology^28^ (**Data Availability**) and 80 10x Visium mouse samples spanning varying degrees of amyloid deposition in 5xFAD mouse model^6^. We assessed the disease informativeness of Spacelink by using ESV and ct-ESV scores from healthy brain cortex, lymph node, and liver to predict gene-level disease prioritizations from PoPS^29^ and MAGMA^30,31^ across 21, 16 and 31 relatively independent (*r*_*g*_ < 0.8) brain, blood and liver-related diseases and traits (**Figure 1c, Supplementary Table S1**). Subsequently, we expanded this analysis to 113 complex diseases and traits (average GWAS sample size = 340,406). All disease-related evaluations of Spacelink are conditioned on putative non-spatial confounders such as the mean and overall variance (spatial + non-spatial) gene expression and cell-type specificity of expression. For the organogenesis and AD neurodegeneration, we assess variation in gene-level ESV scores against developmental time and pathological progression in AD samples (**Figure 1d**). We have publicly released Spacelink hypothesis test results, together with ESV and ct-ESV scores at tissue and cell type levels across different datasets (**Data Availability**).

### Spacelink outperforms other SVG methods in simulation experiments

We evaluated Spacelink against 6 leading global SVG detection methods – Moran’s *I*^32^, nnSVG^8^, SpatialDE^9^, SpatialDE2^10^, SPARK^11^ and SPARK-X^33^ – that demonstrated the highest predictive accuracy across diverse simulation settings in a recent systematic benchmarking analysis^34^. To ensure robust and representative evaluation, we adopted a framework to design 28 spatial simulation datasets, reflective of realistic spatial patterns. These datasets include (i) scDesign3-based simulations parameterized using 10x Visium data from 10 human samples^35^, (ii) covariance-based Gaussian process capturing multi-scale spatial variance^34^, and (iii) custom-designed simulations representing canonical spatial patterns such as gradients, streaks, and hotspots (**Methods, Supplementary Figure S2**). Each simulation dataset included 50% non-SVGs out of the total gene set, allowing for a balanced evaluation of sensitivity and specificity. The scDesign3 framework allows varying the degree of spatial variability assigned to simulated SVGs through tunable parameters, which we leveraged to evaluate different SVG ranking methods (**Methods**). For hypothesis testing, we evaluated Spacelink using statistical power and false discovery rate (FDR). For gene prioritization, we used area under the precision–recall curve (AUPRC) across different p-value cutoffs and Kendall’s tau rank correlation between ESV scores and the ground-truth scDesign3 rankings. We benchmarked Spacelink ESV and p-value based rankings against other methods based on method-specific log-likelihood ratio (nnSVG and SpatialDE), hypothesis test p-values (SPARK, SPARK-X and SpatialDE2), Moran’s I coefficient, FSV (SpatialDE) and PropSV (nnSVG).

Across diverse simulation scenarios, Spacelink consistently achieved high statistical power (mean: 0.792), outperforming other SVG detection methods by 1.07-3.66x on average across the datasets, while also effectively controlling the FDR below the expected 5% across all simulations (**Figure 2a, Supplementary Figure S3a**). Moran’s I showed comparable power to Spacelink in most scenarios but exhibited poor FDR control, yielding inflated false positives across many simulation scenarios. When evaluated using the Kendall’s tau correlation across 10 scDesign3 simulation scenarios, Spacelink ESV score demonstrated the strongest performance, significantly outperforming all spatial metrics across methods (**Figure 2b**). Notably, ESV outperformed comparable metrics such as FSV (SpatialDE) and PropSV (nnSVG), achieving 12.7–37.3% higher correlation across datasets. Moreover, while Spacelink’s global SVG test p-value rankings were slightly less accurate than ESV, they still consistently outperformed p-value-based rankings from other methods by 2.1–12.3% in AUPRC (on average) over the datasets (**Figure 2b, Supplementary Figure S3b**). To assess the robustness of Spacelink to spot dropout, data sparsity, and reduced read depth, we performed (i) spot downsampling at varying probabilities as in Chen et al.^36^, (ii) data sparsification, in which gene expression values were set to zero at selected spatial locations with varying proportions, and (iii) binomial thinning of library size, applied with different probabilities of read-depth reduction following the strategy in Dey et al.^37^ (see **Methods** for details). Spacelink showed robust performance in both SVG identification and gene ranking (based on p-value and ESV) at moderate levels of spot downsampling, data sparsification and library size thinning (**Figure 2c, Supplementary Figure S3c**). In contrast, competing methods showed greater sensitivity to these transformations and degraded more rapidly under extreme conditions. Because nnSVG^8^ assumes a single spatial length scale per gene, we generated exponential kernel-based simulation data in which each SVG was simulated with exactly one true length scale – a design that aligns directly with nnSVG’s modeling assumptions. We then assessed performance by comparing the estimated scales from Spacelink and nnSVG to the ground truth, computing the absolute difference between the true length scale and (i) the single scale estimated scale by nnSVG or (ii) the Spacelink-estimated scale with the largest variance contribution 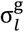 (**Methods**). Despite the simulation being tailored to nnSVG’s framework, Spacelink consistently recovered length scales closer to the ground truth over nnSVG (pairwise t-test p-value = 5e-22). nnSVG also exhibited instability under certain spatial configurations (**Figure 2d, Supplementary Figure S3d**); this likely reflects limitations of nnSVG’s nearest-neighbor Gaussian process (NNGP) approximation^38^. By contrast, Spacelink’s multi-kernel framework is able to capture one or more spatial scales that approximate the true length scale more accurately, even in cases where only a single dominant scale is present in the data.

**Figure 2.**
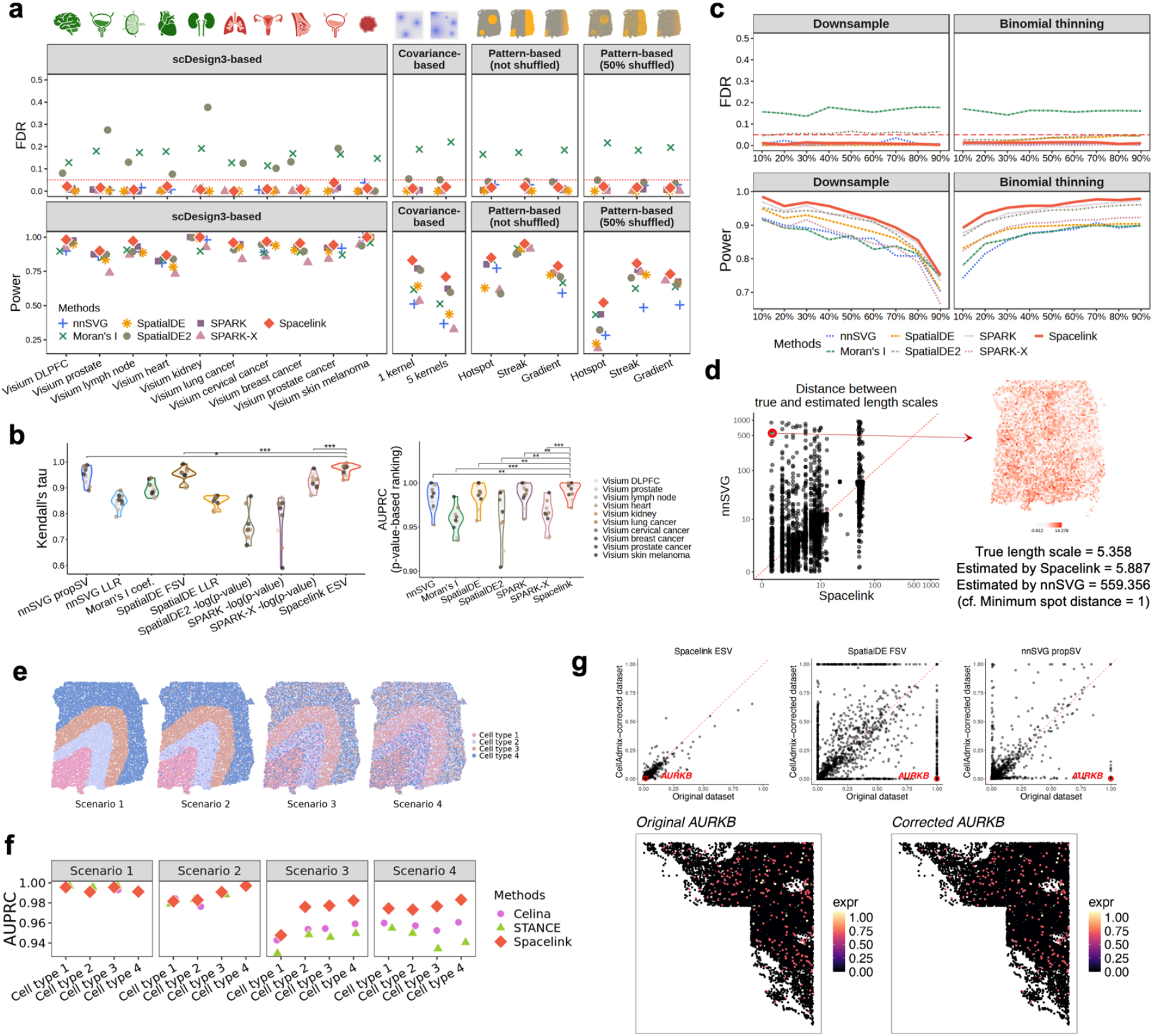
Spacelink outperforms other global and cell type SVG detection and prioritization methods across a broad range of simulation scenarios. (a). Statistical power and false discovery rate (FDR) of Spacelink versus six other global SVG identification methods across 18 simulation settings informed by real-world spatial data and canonical spatial patterns (results for 10 additional settings are shown in **Supplementary Figure S3a**). (b). Left: Kendall’s tau correlation of Spacelink ESV with 8 alternative spatial scoring metrics derived from the 6 methods in Panel (a), across 10 scDesign3-based simulations with known ground-truth ranks. Right: Area under the precision–recall curve (AUPRC) of Spacelink p-values compared with p-values from the six global SVG hypothesis tests across 10 scDesign3-based smulations. Asterisks indicate significance from 1-sided pairwise t-tests comparing Spacelink against each method (*** P < 0.001, ** P < 0.01). (c). Statistical power and FDR of Spacelink against global SVG methods under varying levels of spot downsampling and library size thinning (**Methods**). (d). Left: Comparison of length scale estimates from Spacelink and nnSVG against the ground truth in a set of 3,750 simulated SVGs aligned with nnSVG’s framework. Right: An example simulation where Spacelink closely recovers the true length scale whereas nnSVG does not. (e). Distributions of four cell types in 4 different simulation scenarios constructed based on the human dorsolateral prefrontal cortex (DLPFC) dataset to evaluate ct-SVG methods. (f). AUPRC of Spacelink (ct-SVG) based on hypothesis test p-values compared to Celina and STANCE for each of four cell types across four simulation scenarios. (g) Top: Comparison of Spacelink ESV, SpatialDE FSV, and nnSVG PropSV between the original and corrected CosMx lymph node datasets in a partial tissue region. Bottom: *AURKB* expression in the same datasets. Numerical results are reported in **Supplementary Table S2**.

While our primary benchmarking studies include SVG methods selected based on previous benchmarking analyses^34^, there is a broad class of spatial methods in the statistical literature, that have not been adequately tested at scale using spatial transcriptomics data^39–43^. Therefore, as secondary analyses, we evaluated Spacelink against two such approaches—spNNGP^44^ and spBayes^45^—which, like nnSVG, assume a single spatial scale per gene but use a more elaborate Bayesian inference framework for parameter estimation. spNNGP showed more stable length scale estimates than nnSVG, but lower accuracy than Spacelink and spBayes, likely due to limitations of the nearest-neighbor Gaussian process (NNGP) approximation (**Supplementary Figure S3e**). In contrast, spBayes, based on a Gaussian predictive process, achieved higher accuracy in length-scale estimation than Spacelink, highlighting the competitiveness of Bayesian approaches. However, both Bayesian methods exhibited lower SVG detection power and weaker spatial variability ranking than Spacelink (**Supplementary Figure S3f, S3g**), possibly due to instability in estimating the variance component σ^g^ (determined using posterior mean in these cases), particularly when its true value is small. Using scDesign3 simulations based on Visium human DLPFC data, we examined how the estimated signal-to-noise ratio (SNR) from each method changes with increasing spatial variability parameter *α*. We observed that for spNNGP and spBayes, the SNR was substantially misestimated when *α* was small, whereas no such issue was observed for Spacelink or nnSVG (**Supplementary Figure S3h**). Additionally, the extensive posterior sampling required by these methods leads to a substantial increase in runtime, which presents a practical bottleneck when analyzing tens of thousands of genes across large spatial datasets (**Supplementary Figure S3i**). Overall, although spNNGP and spBayes offer certain advantages—such as more accurate length-scale estimation and the ability to quantify uncertainty in estimates—we demonstrate that Spacelink provides a complementary approach with improved scalability, higher SVG detection power, and more robust signal-to-noise ratio estimation. However, we note that the field of spatial Gaussian Process (GP) models is vast^39–43^, and an exhaustive benchmarking comparison of Spacelink with all such approaches is beyond the scope of this paper.

In order to compare the performance of Spacelink with Celina^13^ and STANCE^12^ for detecting cell type level spatially variable genes (ct-SVGs), we adopted a simulation framework previously proposed in Shang et al.^13^. Under this framework, we simulated single-cell and spot-resolution spatial transcriptomics data from (i) human brain dorsolateral prefrontal cortex (DLPFC)^25^ and (ii) mouse primary visual cortex^46^. We assume that cells belong to four different cell types, each mapped to one of four spatial domains corresponding to cortical layers. We designed four distinct scenarios (Scenarios I–IV), varying in the degree of cell type mixing and spatial domain overlap (**Figure 2e, Supplementary Figure S4a**; see **Methods** for details). For each scenario, we first simulated single-cell resolution expression data for 2,304 genes, including 1,536 ct-SVGs and 384 marker genes that were global SVGs but not ct-SVGs, with varying expression levels across cell types or spatial patterns (see **Methods** for design considerations including the number of genes). Spot-resolution data were then generated by overlaying square grids onto the single-cell resolution data and aggregating expression counts of cells within each grid to obtain spot-level expression. We applied ct-SVG methods to this spot-resolution data using either oracle or RCTD-estimated cell type proportions^23^, with the simulated single-cell data serving as the reference for RCTD. We benchmarked the predictive accuracy of Spacelink against other methods for each cell type and each scenario using AUPRC. In the human DLPFC simulation using RCTD-estimated cell type proportions, Spacelink achieved significantly higher AUPRC values than Celina and STANCE, with maximum improvements of 2.5- and 4.5%, respectively, across different settings (**Figure 2f**). Similar trends were observed for the mouse visual cortex simulation (**Supplementary Figure S4b**) and when using oracle cell type proportions over estimated RCTD proportions (**Supplementary Figure S4c**). Spacelink’s ct-ESV scores effectively prioritized true ct-SVGs over non-ct-SVGs across all scenarios (**Supplementary Figure S4d**). Note that due to differences in levels of cell type mixing across layers in the simulation design, the cell type level spatial bandwidths and consequently, the ct-ESV values, are not directly comparable across scenarios. Celina and STANCE were not included in this comparison, as they do not provide a cell-type-specific spatial variability metric like ct-ESV. Finally, since Spacelink differs from Celina and STANCE in (i) employing kernel matrices with data-driven spatial bandwidths and (ii) using a data-driven gating strategy to correct for spatial colocalization in weakly represented cell types, we evaluated the relative performance gain contributed by each of these innovations. The primary performance gain was attributable to the data-driven kernel matrices (**Supplementary Figure S4e**), while the gating mechanism provided an additional modest improvement in performance compared to including all random effects from other cell types (as in STANCE) and excluding them entirely (as in Celina) (**Supplementary Figure S4f**).

We further evaluated the sensitivity of Spacelink ESV to technical artifacts such as cell segmentation errors that generate spurious short-range autocorrelation. To demonstrate this, we applied the cellAdmix correction method^47^ to the CosMx lymph node dataset that corrects for transcript spillover from neighboring cells, and compared Spacelink ESV with nnSVG PropSV and SpatialDE FSV between the corrected and uncorrected data. While Spacelink ESV produced consistent estimates between the corrected and uncorrected data, PropSV and FSV showed substantial discrepancies (**Figure 2g**). To illustrate this, we highlight a specific case example of *AURKB* gene whose PropSV and FSV decreased from 1 (uncorrected) to 0 (corrected); in this instance, the apparent spatial signal was driven entirely by short-range spurious autocorrelation due to transcript spillover, which was eliminated upon correction. More broadly, when both σ^g^ and τ^g^ are estimated to be small after correction, PropSV (σ^g^/(σ^g^ + τ^g^)) becomes numerically unstable due to the near-zero denominator, yielding unreliable estimates regardless of true spatial structure. Across all cases, Spacelink ESV remained consistently low, demonstrating robustness to cell segmentation errors.

We assessed the p-value calibration of Spacelink using 20 distinct null simulations (**Methods**). Q–Q plots indicate that Spacelink yields well-calibrated p-value distributions in most cases (**Supplementary Figure S5a, S5b**). In terms of the K–S distance, Spacelink achieves smaller values than most competing methods, with the exception of SPARK and SPARK-X (average K–S distance across simulation settings: Spacelink = 0.15, SPARK = 0.09, SPARK-X = 0.09, other methods = 0.20–0.68) (**Supplementary Figure S5c**). Under an additional high-sparsity null simulation (**Methods**), Spacelink showed some p-value inflation, performing worse than SPARK-X (**Supplementary Figure S5d**). This, together with the lower statistical power of Spacelink compared to SPARK-X under high sparsity (**Supplementary Figure S3c**), reflects a known limitation of applying a Gaussian likelihood to normalized versions of highly sparse count data, as discussed in Zhu et al. (2021)^48^. This suggests that Spacelink’s performance is limited in extreme sparsity settings and that SPARK-X may be preferred for SVG hypothesis testing under such conditions. Despite this limitation, we demonstrate that Spacelink has practical utility across a broad range of sparsity regimes. Spacelink’s AUPRC and power are within 10% of SPARK-X for sparsity levels below 95% (**Supplementary Figure S5e**). The Visium, CosMx, and Stereo-seq datasets analyzed in this paper exhibit 26–61%, 86–93%, and 86–94% sparsity, respectively (**Supplementary Table S8**), which all fall within or near this range. Also, in practice, Slide-seq v2 and similar high-resolution platforms are routinely analyzed after spatial binning or aggregation^49,50^, which reduces per-gene sparsity to levels where methods such as Spacelink are applicable. We further show that under high-sparsity settings (>80%), Spacelink applied to binned data achieves higher AUPRC, power, and Kendall’s τ compared to the standard un-binned model, with overall performance more comparable to SPARK-X (**Supplementary Figure S5f**). Furthermore, Spacelink achieves 3.7x higher statistical power on average across standard alternative simulation settings (**Figure 2a**). Critically, SPARK-X does not provide a continuous spatial variability-based gene-ranking metric analogous to ESV. Therefore, Spacelink offers important complementary advantages over SPARK-X.

We compared the computational efficiency and performance of Spacelink and Spacelink-lite. **Supplementary Figure S6a** present average runtime and peak memory usage for the original Spacelink, Spacelink-lite, and other SVG methods across a range of simulated dataset sizes extended up to hundreds of thousands of spots (**Methods**). Methods requiring more than 100 GB of memory or exceeding 40,000 seconds of runtime per gene were excluded due to resource constraints. The original Spacelink achieved comparable speed to other fast global SVG methods for small to moderate datasets (a few hundred to several thousand spots) but was not efficient for larger datasets. Spacelink-lite, in contrast, maintained high speed and memory efficiency even at very large scale. We validated that Spacelink-lite produces results closely comparable to the original Spacelink by comparing FDR and power across 10 scDesign3-based simulations and 6 covariance-based simulations. Spacelink-lite showed nearly identical performance (**Supplementary Figure S6b**), and ESV estimates between the two versions were highly correlated (average r = 0.99, **Supplementary Figure S6c**). To further assess large-scale performance, we generated covariance-based simulations within the spatial domains of CosMx cortex and liver datasets and compared Spacelink-lite against other computationally feasible methods for large datasets, including nnSVG, Moran’s I, and SPARK-X. In these settings, Spacelink-lite consistently achieved higher power while controlling FDR (**Supplementary Figure S6d**). Among ct-SVG methods, Spacelink was faster than STANCE but slower than Celina (**Supplementary Figure S6e**), which is attributable to Spacelink’s use of a more complex model incorporating an expanded set of random effects in select cases to correct for the effect of other colocalizing cell types. All timing and memory profiling experiments were conducted on an x86_64 HPC cluster node equipped with 2 CPUs (18 cores per CPU, 72 logical threads); runtime and memory usage were obtained from the CPU time and maximum resident memory (Max RSS) reported in the LSF job stdout logs.

We performed five secondary analyses. First, we evaluated how ESV varies with the number of spots, level of data sparsity, and degree of technical noise across tissues with different structural complexity (kidney, prostate cancer, and DLPFC; **Supplementary Figure S7a**) using simulated datasets (see **Methods**). We compared ESV behavior against PropSV (nnSVG), FSV (SpatialDE), and Moran’s I coefficient. Randomly reducing the number of spots increases the apparent complexity of spatial structure; however, ESV and Moran’s I remained relatively stable across resolutions, whereas PropSV and FSV fluctuated substantially, indicating greater robustness of ESV at lower spot densities (**Supplementary Figure S7b**). Introducing sparsity is expected to weaken spatial autocorrelation for all metrics, but compared to other scores, ESV remained more stable. At 25% imposed sparsity, 29% of genes exhibited less than 10% change in ESV scores on average across tissues, compared to less than 3% for competing metrics (**Supplementary Figure S7c)**. Increasing technical noise variance parameter weakens spatial autocorrelation by design, and should in principle lead to the reduction of all spatial variability metrics. We observe this expected decreasing trend across all metrics and tissues; however, ESV exhibited a more consistent monotonic decrease with increasing noise variance, with a more negative Kendall’s *τ* correlation with noise variance (−0.99 vs. −0.92 to −0.97; **Supplementary Figure S7d**). Overall, these results indicate that the Spacelink ESV metric is more stable under different data quality degradation scenarios compared to other SVG rank metrics across tissue contexts. Second, to assess the impact of kernel choice, we conducted supplementary comparisons using alternative stationary and non-stationary kernels alongside or instead of the exponential kernel in scDesign3-based simulations (**Supplementary Figure S8a**). These analyses revealed no substantial differences in performance. Overall, the default kernel performed better than or similarly to other kernel families, indicating that while the exponential kernel is a principled default, the results are robust to this modeling choice. Third, to assess the robustness of Spacelink to imbalanced data, we generated simulations with varying proportions of non-SVGs, ranging from 5% to 95% of the total gene set. Spacelink demonstrated robust performance and outperformed the other methods across all settings (**Supplementary Figure S8b**). Fourth, using the human DLPFC Visium cortex data with annotated cortical layer information^25^, we evaluated the accuracy of domain detection using the top Spacelink ESV genes, compared to the same number of genes prioritized using other spatial metrics. Spacelink achieved comparable or superior accuracy in identifying spatial domains relative to other methods (**Supplementary Figure S8c**).

The simulation results demonstrate that Spacelink provides accurate, robust, and interpretable detection and prioritization of spatially variable genes across a wide range of spatial patterns and cell type configurations, consistently outperforming existing methods in statistical calibration and spatial signal recovery.

### Spacelink outperforms other SVG methods in cross-platform consistency across tissues

A desirable quality in a good spatial metric is robustness in gene prioritization across different spatial platforms and assays from matched tissue systems. To assess this, we applied *Spacelink* to single-cell resolution CosMx spatial transcriptomics data and matched spot-resolution 10x Visium data from 3 healthy human tissues^25,26^ - brain cortex (CosMx: 188,686 cells, 6,278 genes; Visium: 3,639 spots, 33,525 genes), lymph node (CosMx: 1,852,946 cells, 6,175 genes; Visium: 4,035 spots, 9,116 genes), and liver (CosMx: 332,877 cells, 1,000 genes; Visium: 3,200 spots, 36,602 genes) (**Figure 3a, see Data Availability**). These tissues were selected because they are linked to a broad range of diseases and quantitative traits with well-powered genome-wide association studies (GWAS), making them particularly informative for disease-related benchmarking of Spacelink^51–53^. To ensure fair cross-platform comparison, we rasterized the CosMx datasets using SEraster^54^ and matched the average number of cells per spot to the corresponding Visium data for each tissue (4.69 cells in brain cortex, 44.13 in lymph node, and 11.06 in liver). We note that rasterization had minimal impact on Spacelink analyses, as evidenced by high Spearman correlations of ESV estimates (0.75, 0.81, 0.85) and strong concordance in SVG test results (odds ratios: 41.3, 24.4, 74.1) between Spacelink-lite applied to the original CosMx data and the original Spacelink applied to rasterized data across three datasets (**Supplementary Figure S9a**). We observed strong correlations in ESV scores between CosMx and Visium across all three tissues (r=0.77 in brain cortex, 0.60 in liver, and 0.59 in lymph node) (**Figure 3b**). Consistently across platforms, Spacelink selected fewer spatial bandwidths with non-zero weights for genes in brain cortex compared to lymph node and liver (**Figure 3c, Supplementary Figure S9b**). Moreover, genes in the brain cortex exhibited, on average, 3.3- and 5.6-fold larger normalized spatial length scales—scaled by the minimum distance among spots in each tissue—relative to the lymph node and liver, respectively. These patterns likely reflect the simpler layered organization of gene expression in brain cortex, characterized by long-range spatial variability, in contrast to the more heterogeneous architecture of lymphoid and hepatic tissues (**Supplementary Figure S9c**). We highlight two example genes, *CNP* and *EIF5B*, with consistent high and low spatial variability in brain cortex across platforms; in contrast, gene *SCGB2A2* shows specific spatial variability only in 10x Visium data (**Figure 3d, Supplementary Figure S9d**).

**Figure 3.**
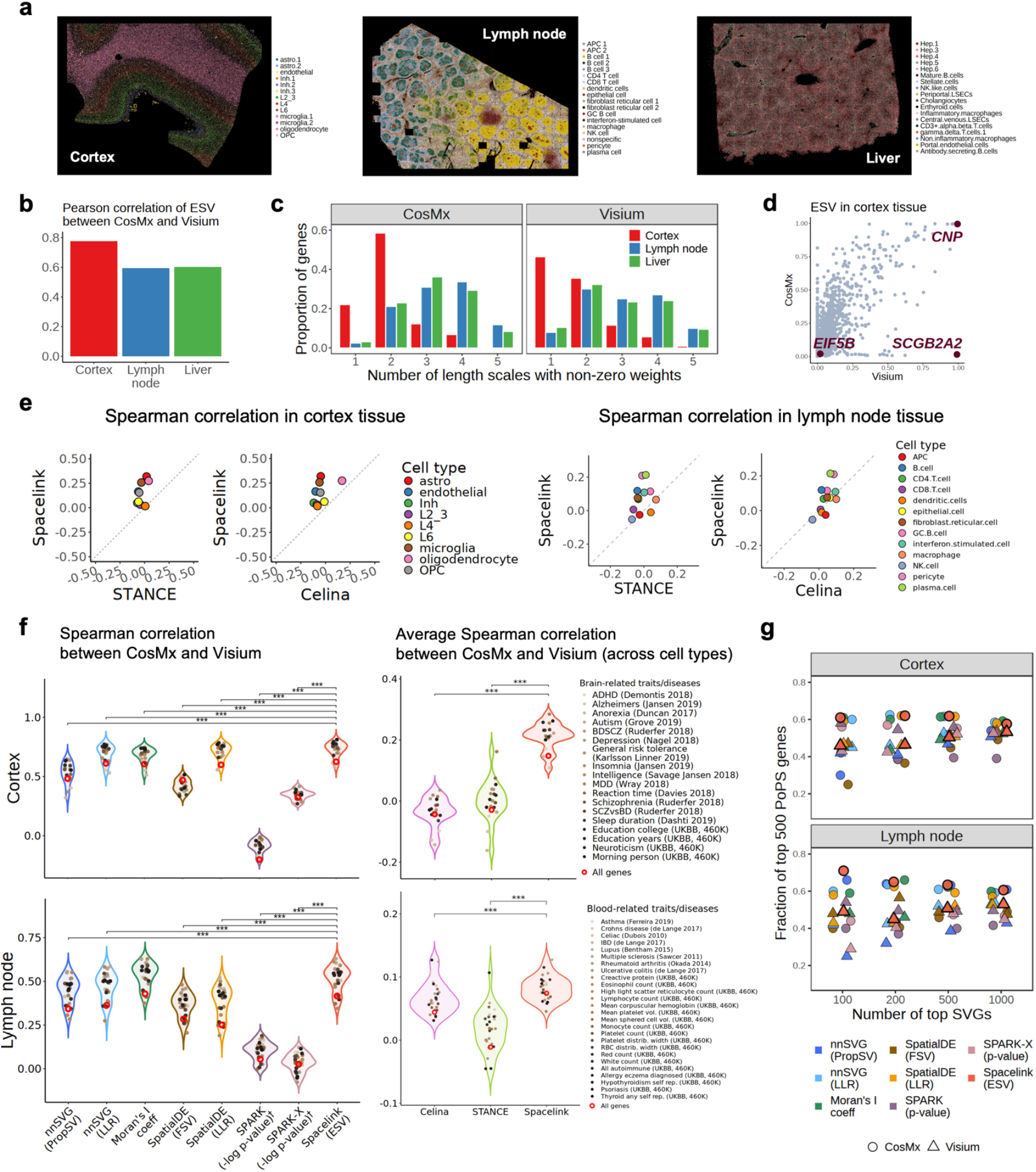
Spacelink demonstrates superior cross-platform consistency in quantifying spatial variability compared to other methods. (a). Spatial annotations of major cell types in CosMx single-cell resolution SRT data from three human tissues—brain cortex, lymph node, and liver—used as the reference for cross-platform comparison. (b). Pearson correlations of Spacelink global ESV scores, calculated on matched gene panels, between CosMx and Visium data across the three tissues. (c). Proportion of genes with different numbers of spatial length scales with non-zero estimated weights in Spacelink whole tissue resolution models applied to CosMx and Visium data in the three tissues. (d). Scatter plot of global ESV values for all genes in the brain cortex CosMx and Visium panels. (e). Cross-platform consistency of cell type– specific SVG (ct-SVG) methods (Spacelink, Celina, STANCE) between CosMx and Visium panels for two matched tissue systems (left: brain cortex, right: lymph node), assessed by Spearman correlations of p-values for each cell type. Scatter plots show correlations across cell types in brain cortex (left) and lymph node (right), comparing Spacelink against Celina and STANCE. (f). **Left:** Spearman correlations of Spacelink ESV and seven alternative metrics between CosMx and Visium data, restricted to tissue-relevant disease/trait genes in cortex (top) and lymph node (bottom). ^†^Since SPARK and SPARK-X do not provide metrics quantifying the extent of spatial variability, we instead used −log p-values. A comprehensive comparison of p-value–based metrics across all methods is presented in **Supplementary Figure S11b. Right:** Average Spearman correlations of p-values from Spacelink, STANCE, and Celina across cell types between CosMx and Visium, restricted to tissue-relevant disease/trait genes in the cortex (top) and lymph node (bottom). In both sets of panels, red circles denote values computed using all genes. Other circles correspond to different tissue-relevant complex diseases and traits. Asterisks indicate significance from 1-sided pairwise t-tests (*** P < 0.001). (g). Fractions of the union of the top 500 PoPS-prioritized genes for tissue-matched traits implicated as top N ranked global SVGs based on Spacelink ESV and 7 other spatial gene prioritization metrics applied to CosMx and Visium data from cortex (top) and lymph node (bottom). We consider 4 values of N (100, 200, 500 and 1,000). Numerical results are provided in **Supplementary Table S3**.

Next, we evaluated the cell-type-specific version of Spacelink against Celina and STANCE by comparing correlations of p-values obtained from Visium and CosMx data. In Visium, we applied each ct-SVG method using cell type proportions estimated via RCTD^23^ with CosMx as the reference (**Supplementary Figure S10a**). For the single-cell-resolution CosMx data, methods were applied directly to cells from the target cell type. Spacelink achieved up to 1.38x, 1.22x, and 1.22x higher cross-platform Spearman correlations than Celina and STANCE in the brain cortex, lymph node, and liver, respectively (**Figure 3e, Supplementary Figure S10b**). These findings underscore the robustness of Spacelink in identifying ct-SVGs across platforms.

We further evaluated the cross-platform consistency by focusing on genes relevant to relatively independent (*r*_*g*_ < 0.8) complex diseases and traits linked to each tissue (**Supplementary Table S1**). We also used this consistency measure as a metric to evaluate Spacelink gene prioritization based on ESV and p-value against other spatial metrics. As primary analysis, we benchmarked Spacelink ESV against other metrics quantifying the extent of spatial variability – nnSVG (PropSV), nnSVG (LLR), Moran’s I coefficient, SpatialDE (FSV), SpatialDE (LLR). As secondary analysis, we compared cross-platform consistency of rankings based on Spacelink p-values against SVG hypothesis test p values from 6 other methods. When restricted to the top 500 tissue-relevant PoPS disease genes^29^, Spacelink ESV and other spatial metrics achieved consistently stronger cross-platform correlations—improving by 10%, 30% and 20% in brain, lymph node and liver respectively (**Figure 3f, Supplementary Figure S11a**). Notably, Spacelink ESV scores exhibited significantly (pairwise t-test, P < 0.01) stronger correlations between CosMx and Visium platforms compared to 5, 4 and 4 out of the 5 competing methods when restricted to tissue-relevant disease genes in brain cortex, lymph node and liver respectively. Spacelink’s p-value-based metric showed lower correlation than the ESV score, it still outperformed all six other p-value-based metrics from competing methods (P < 0.05) in all 3 tissues (**Supplementary Figure S11b**). Spacelink (ct-SVG) achieved significantly higher average cross-platform correlations (across cell types) for disease-prioritized genes than STANCE and Celina in the brain cortex and lymph node, and higher than Celina in the liver (P < 0.001) (**Figure 3f, Supplementary Figure S11a**). Together, these results highlight that Spacelink consistently delivers stronger cross-platform reproducibility than other methods, particularly for disease-associated genes across diverse tissue contexts.

We systematically compared the CosMx and Visium spatial platforms in terms of disease informativeness of genes matched across panels. Using Spacelink-detected SVGs at varying rank cutoffs (top 100, 200, 500, and 1000 by ESV), we quantified overlap with the union of the top 500 PoPS-prioritized genes for tissue-matched traits. For this comparison, we subsetted the two panels to the same set of genes for a fair comparison. Across all thresholds, CosMx consistently outperformed Visium, yielding on average 24%, 32%, and 26% higher excess overlap in brain cortex, lymph node, and liver (**Figure 3g, Supplementary Figure S11c**). Similar patterns were observed across other SVG detection methods, underscoring the broader advantage of CosMx over Visium for matched genes for downstream disease association studies; however, CosMx panels assay a more limited set of genes compared to Visium’s whole-transcriptome coverage.

Finally, we note that although SPARK-X exhibited better p-value calibration than Spacelink under extreme sparsity conditions in simulations, across all three tissues, Spacelink p-value-based scores showed significantly stronger cross-platform concordance than those of SPARK and SPARK-X when restricted to tissue-relevant disease genes (pairwise t-test, P < 0.01; **Supplementary Figure S11b**). In **Supplementary Figure S11c**, performance was comparable between Spacelink and SPARK-X in liver, whereas Spacelink demonstrated superior performance in brain cortex and lymph node. Together, these results indicate that both Spacelink ESV and p-values yield more reproducible gene rankings across platforms than SPARK and SPARK-X, particularly for disease-associated genes.

In summary, Spacelink delivers consistently stronger cross-platform reproducibility of spatially variable genes, especially for disease-relevant genes, outperforming existing methods across a diverse set of tissues.

### Spacelink prioritized genes are uniquely informative for human diseases

We assessed the relevance of spatially variable genes (SVGs) and cell-type-specific SVGs (ct-SVGs), identified by Spacelink in CosMx brain cortex, lymph node, and liver datasets, for 113 complex traits and diseases (average GWAS sample size = 340,406) (**Supplementary Table S1**). First, we constructed Spacelink-based SVG programs defined by ESV (whole-tissue) and ct-ESV (cell-type-specific) scores. These programs were tested for association with two disease-relevant gene sets: (i) top 500 PoPS-scored genes for brain-, blood-, and liver-related traits, and (ii) MAGMA-prioritized genes (z-score > 3) for the same traits (**Supplementary Table S1**). Because SVG detection methods often prioritize genes with high expression levels or elevated expression variability, we fit logistic regression models associating each ESV program (X) with disease gene set of each trait (Y) while conditioning on putative confounders (Z) such as mean and variance of log-transformed and normalized expression. To enable balanced evaluation across programs, we computed adjusted Odds Ratios (aORs) and probabilistic recall against the disease gene sets (**Methods**). We benchmarked Spacelink ESV and ct-ESV programs against (i) sc-linker probabilistic cell type programs^51^ (genes enriched in specific cell types) and (ii) gsMap cell-level gene specificity scores (GSS)^7^. All programs are non-negative by design, and to ensure fair comparison, each was rescaled to the [0,1] interval (**Methods**). **Figure 4a** shows the aOR and probabilistic recall of programs with positive and significant regression coefficients (P < 0.05) for each trait. In brain cortex and lymph node, Spacelink global ESV programs achieved 2.6x and 2.2x higher aOR on average and 1.5x and 2.3x higher recall on average compared to sc-linker and GSS programs, while ct-ESV programs showed 5.5x and 4.7x higher aORs but lower probabilistic recall compared to other methods. In liver, only a small number of programs reached significance, likely due to the limited number of genes in the dataset (1,000), which constrained detecting strong trends. Results for ct-ESVs remained consistent when additionally adjusting for the mean and variance of corresponding cell-type level normalized expression (**Supplementary Figure S12a**). Replacing PoPS-scored genes with MAGMA-prioritized genes in the regression models resulted in weaker associations across all programs (**Supplementary Figure S12b**). The restricted gene panels in CosMx and other single-cell–resolution platforms such as Xenium limit the application of genome-wide heritability analyses methods. We note that 60% of cell types implicated fewer than 100 genes even at a relaxed cut-off of ct-ESV (>0.2), all of which when mapped to enhancer-gene links relevant to the tissue, as in sc-linker, generated SNP annotations with very small average annotation size (<0.001); this annotation size is deemed unsuitable for genome-wide heritability regression-based approaches such as S-LDSC^51,55^.

**Figure 4.**
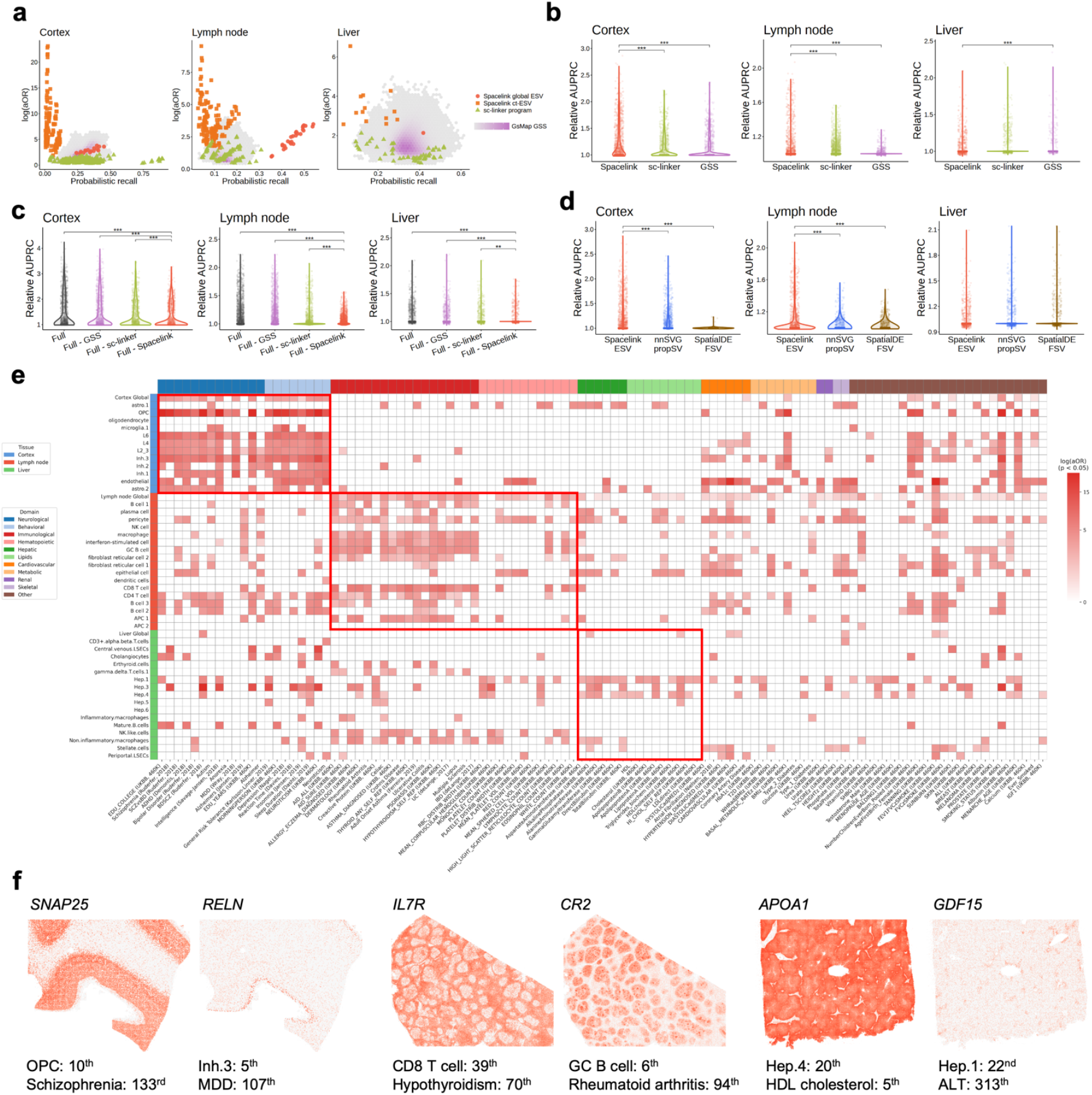
Spacelink-prioritized genes are uniquely informative for complex diseases and traits. (a). Logistic regression analyses associating each gene program (probabilistic gene score between 0 and 1) with the top 500 PoPS-prioritized disease gene set for each tissue-matched trait, conditioned on putative non-spatial confounders (overall mean and variance of log-normalized expression), across brain cortex, lymph node, and liver. Gene programs from Spacelink, sc-linker, and gsMap GSS were evaluated using log-transformed adjusted odds ratio (aOR) and probabilistic recall against the disease gene set. Details of the aOR and probabilistic recall calculations are provided in **Methods**. (b). Method-specific joint regression analyses of all nominally significant (p < 0.05) global and cell-type level Spacelink ESV, sc-linker, and GSS programs in each tissue from the marginal analyses in (a). Similar to (a), the regression model is conditioned on overall mean and variance of log-normalized expression. For GSS, we only include programs that have correlation < 0.5, to ensure less collinearity across programs in the model. Relative AUPRC (against baseline AUPRC; see **Methods**) was used to evaluate each joint regression model with respect to each tissue-matched trait. Asterisks indicate significance from 1-sided pairwise t-tests (*** P < 0.001, ** P < 0.01). (c). Full joint regression analyses (“Full”) including all gene programs across the 3 methods, Spacelink, sc-linker, and GSS programs, that are nominally significant (p < 0.05) Spacelink, sc-linker, and GSS programs. For GSS, we only include programs that have correlation < 0.5, to ensure less collinearity across programs in the model. We also consider reduced versions of the full model each excluding sets of programs from each method separately. For example, “Full – GSS” excludes all nominally significant GSS programs from the gsMap method. Relative AUPRC (against baseline AUPRC; see **Methods**) was used to evaluate each joint regression model with respect to each tissue-matched trait. Asterisks indicate significance from 1-sided pairwise t-tests (*** P < 0.001, ** P < 0.01). (d). Joint regression analyses of all whole-tissue and cell-type level nominally significant (p < 0.05) spatial gene programs corresponding to Spacelink ESV, nnSVG PropSV and SpatialDE FSV; significance is assessed using 1-sided pairwise t-tests (*** P < 0.001, ** P < 0.01). (e). Heatmap of disease or trait-specific enrichment (using log-normalized adjusted odds ratio: log(aOR), conditioned on cell type level overall mean and variance of cell-type resolution sctransform expression data) of 48 Spacelink global ESV and cell-type level ct-ESV programs from three tissues across 113 complex diseases and traits. The top 500 PoPS-prioritized genes were used in the regression model for each trait, as suggested in Weeks et al.^29^. Only results for nominally significant (p < 0.05) log aORs from the disease association analyses are shown. (f). Spatial expression plots of six top-ranked ct-SVGs that are also top PoPS-prioritized genes for relevant traits in brain cortex (left), lymph node (middle), and liver (right). Numbers next to each cell type and trait indicate the gene’s ranks within the full panel based on ct-ESV and PoPS scores, respectively. Numerical results are reported in **Supplementary Table S4**.

Next, we performed a joint regression analysis of all nominally significant (P < 0.05) Spacelink ESV programs, sc-linker programs, and GSS programs in each tissue from the marginal analysis above, conditioning on the same set of putative confounders as in **Figure 4a** (**Methods**). Because many GSS programs are highly correlated due to expression similarity across neighboring cells, we restricted the joint analysis to those with pairwise correlations < 0.5. For each disease, we implemented a chromosome-based cross-validation scheme by randomly splitting all genes into two equal groups of training and evaluation set 50 times. For each split, we performed a logistic regression model with a ridge penalty on the training set, and evaluated the performance on the held-out evaluation set. The evaluation metric, relative AUPRC, was defined as the ratio of the model’s test-set AUPRC to the baseline AUPRC (**Methods**). Spacelink ESV programs achieved significantly higher relative AUPRC than GSS programs across all tissues (up to 2.15x higher) and outperformed sc-linker programs in every tissue except liver (up to 2.09x higher; pairwise t-test, P < 0.001) (**Figure 4b**). We next compared a full joint regression model—including all significant Spacelink, sc-linker, and (moderately uncorrelated) GSS programs—with three reduced models, each excluding one program set (Full – Spacelink, Full – sc-linker, Full – GSS). Across tissues, removing Spacelink programs led to significantly lower relative AUPRC than the other models (pairwise t-test, P = 1e-03 or lower) (**Figure 4c**). Notably, models combining Spacelink with other programs (Full, Full – sc-linker, Full – GSS as in **Figure 4c**) outperformed Spacelink-only model (as in **Figure 4b**), especially in brain cortex. This highlights that Spacelink captures complementary disease-relevant information to sc-linker and gsMap and that combining methods can be advantageous. Next, we benchmarked Spacelink against programs derived from other SVG methods. In the marginal regression analysis of individual programs as in **Figure 4a**, Spacelink global ESV programs achieved higher probabilistic recall (1.1x and 1.3x) for comparable aOR against PropSV (nnSVG) and FSV (SpatialDE) respectively (**Supplementary Figure S12c**). When applied to individual cell types, Spacelink ct-ESV exhibited 2.8x and 8.6x higher aOR compared to analogous cell-type PropSV and FSV scores (**Supplementary Figure S12c**). In the joint regression model comprising all cell type and global programs for each SVG method, Spacelink ESV programs cumulatively achieved significantly higher relative AUPRC in both cortex and lymph node (pairwise t-test, P < 0.001) (**Figure 4d**). When using binary indicators of the top N SVGs (N = 1,000 in cortex and lymph node; N = 500 in liver) defined by p-value, all six alternative methods—and even Spacelink’s own p-value–based programs—showed markedly lower performance than Spacelink ESV programs in cortex and lymph node (**Supplementary Figure S12d, S12e**). In liver, results were similar across methods, likely due to the smaller gene panel; analyses with a broader gene set may uncover clearer performance differences. To further demonstrate the practical utility of Spacelink in comparison to SPARK-X, which exhibits better p-value calibration under extreme sparsity conditions in simulations, we compared ESV-based disease informativeness from CosMx brain DLPFC and lymph node data using two approaches: Spacelink-lite applied to the full high-sparsity data, and Spacelink applied to rasterized data. In both cases, disease informativeness was higher than the SPARK-X p-value-based metric (up to 2.1x and 2.9x higher relative AUPRC, respectively; **Supplementary Figure S12f**), demonstrating that Spacelink ESV retains its advantage over SPARK-X in real high-sparsity datasets under standard analytical workflows.

Finally, we examined individual disease-specific enrichment of 48 Spacelink ESV programs (whole tissue and cell type resolution) from the 3 CosMx tissues against 113 complex diseases and traits, using the adjusted odds ratio (aOR) metric (conditional on mean and variance of normalized expression confounders at the whole-tissue level for whole-tissue programs and at the cell-type level for cell type programs) with respect to top 500 PoPS-prioritized gene sets (**Figure 4e; Methods**). As expected, we observed strong alignment between tissue-resident ct-SVGs and diseases/traits relevant to the corresponding tissue. Ct-ESV programs in brain and lymph node showed 5.93x and 2.02x higher disease informativeness for related traits, compared to other traits (**Figure 4e**). Ct-ESV programs in liver did not show higher disease informativeness for related traits, likely due to the small gene panel. We also observed several notable cross-tissue disease informativeness of ct-SVGs for diseases and traits – such as brain endothelial cells associated with Coronary Artery Disease (CAD)^56^ and blood pressure^57^, liver Hep.3 hepatocyte subtype associated with Schizophrenia^58^, lymph node macrophages associated with lung capacity (FEV1FVC) adjusted for smoking^57^. Endothelial cells form the lining of heart and the broader circulatory system, and are known to become dysfunction in CAD^59,60^. Previous studies have shown increased prevalence of liver disease among individuals with Schizophrenia^61,62^, and spatial programs in Hep,3 hepatocyte subtype may affect shared pathways like ferroptosis between schizophrenia and metabolic dysfunction^63^. Macrophage subtypes have been implicated in lung fibrosis due to their spatial interaction with tissue-resident fibroblasts, promoting fibrotic processes^19,64^. We considered a secondary analysis by replacing PoPS with MAGMA-prioritized (Z score > 3) genes in the disease enrichment analysis in **Figure 4e**; the resulting enrichment signals were weaker but still showed consistent tissue-specific patterns (**Supplementary Figure S12g**).

We highlight several top-ranked ct-SVGs with high PoPS scores for relevant traits (**Figure 4f**). *SNAP25*, ranked 10^th^ in terms of ct-ESV in brain cortex and among top 150 PoPS-scored genes for Schizophrenia^58^, a synaptic gene associated with multiple neurodevelopmental disorders, is primarily expressed in neurons but has also been shown to be regulated in OPCs^65^. *RELN*, a top PoPS-prioritized gene (rank 107) for Major Depressive Disorder and the 5th strongest ct-ESV gene in inhibitory neurons (Inh.3 subtype), is implicated to affect specific groups of inhibitory neurons and the development of inhibitory synapses^66^. *CR2*, a top 100 PoPS gene (rank=94) for Rheumatoid Arthritis (RA)^67^ and a top 10 (rank=6) ct-ESV gene in germinal center B cells, is a known B-cell surface complement receptor whose dysregulation contributes to B-cell hyperactivation and autoantibody production^68^. *GDF15*, a top-ranked ct-ESV gene in Hepatocyte 1 (Hep.1) cell subtype in liver and top 500 PoPS gene for Alanine Aminotransferase (ALT) (rank=313), is a stress- and injury-responsive cytokine, which has been found to be associated with plasma ALT concentrations in obese patients with metabolic dysfunction-associated liver disease^69,70^.

In summary, our results demonstrate that Spacelink-derived SVG and ct-SVG programs provide uniquely informative signals for complex traits and diseases, complementing existing approaches and revealing biologically plausible gene–cell type associations for disease. We note that our conclusions depend on the quality of both the SRT and GWAS data. All the GWAS data we used here are publicly available from the PoPS study^29^, and the traits have generally been selected based on the largest available GWAS meta-analyses and are largely well-powered with an average sample size of 340K. We also caution that our spatial program informativeness for disease should not be interpreted as causal link to disease, as these programs may reflect consequences or correlates of disease. However, this is typical of all leading program-to-disease GWAS mapping papers in the field^29,51,71,72^ that focus on gene programs from single-cell RNA-seq and CRISPR perturbation data and associate them with disease GWAS. It is very difficult to establish causality of a program for disease based on genetic data alone, however the goal here is to nominate and prioritize some of these programs through disease associations to perform functional follow-up experiments targeting underlying genes and pathways in the lab.

### Spacelink detects temporal dynamics in spatial variability during mouse organogenesis

To investigate how spatial variability in gene expression evolves during embryonic development, we applied Spacelink to single-cell resolution Stereo-seq spatiotemporal transcriptomics data^27^ spanning eight stages of mouse organogenesis (E9.5 to E16.5; 5,913–121,767 spots, 25,201–28,798 genes) that capture major developmental transitions (**Figure 5a**). To ensure computational feasibility, we used the “Bin50” resolution dataset (as in previous studies) and further rasterized it with SEraster^54^ (**Methods**), yielding spatial spots with an average of 13–14 bins across stages. Because embryos increase in size over time, we corrected for potential systematic biases in raw ESV scores by estimating stage-specific effects and subtracting them from each gene’s ESV values prior to regression analysis (**Methods**). In addition to whole-embryo analyses, we also conducted separate analyses of brain and non-brain compartments, with separate rasterization performed independently for each. This design was motivated by two factors: (i) the brain is robustly represented across all eight stages, consistently accounting for 14–26% of embryo size, whereas other tissues typically cover a much smaller fraction (average 5%) and may be absent at certain stages; and (ii) brain-derived Spacelink SVG programs provided the strongest disease-informative signals compared to other tissues (**Figure 4**), motivating deeper investigation of its developmental spatial architecture. Due to the absence of well-matched dynamic single cell reference atlases matching the exact stages of mouse organogenesis, we did not perform ct-SVG analysis with Spacelink.

**Figure 5.**
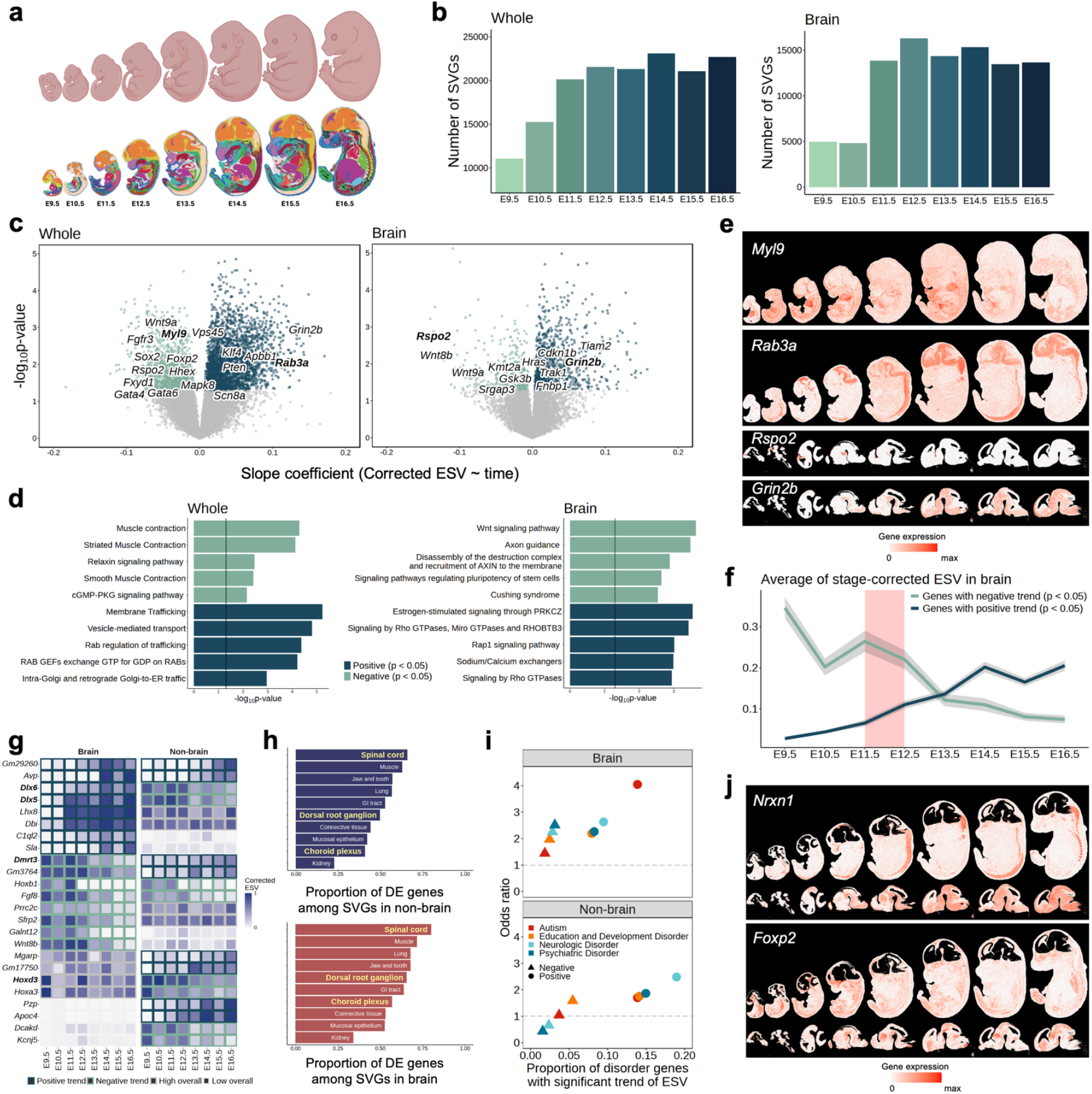
Spacelink detects temporal dynamics in spatial variability during mouse organogenesis. (a). Schematic illustrating mouse embryonic Stereo-seq SRT data across 8 stages (E9.5–E16.5) tracking organogenesis; bottom panel shows spots colored by tissue type. (b). Number of SVGs identified by Spacelink at each stage within the whole embryo (left) and the brain region (right), with bars shaded from light to dark green across developmental time. (c). Volcano plots of gene-level regressions associating stage-corrected ESV (adjusted for embryo size; **Methods**) with developmental time in the whole embryo (left) and brain (right). Each point represents the regression slope (x-axis) and –log10 p-value (y-axis). Genes with nominally significant (P < 0.05) negative and positive associations are shown in light and dark green, respectively. (d). Pathway enrichment analyses (based on ConsensusPathdb^74^) of genes with significantly negative (light green) and positive (dark green) associations between spatial variability and developmental progression in the whole embryo (left) and brain region (right). (e). Spatial expression plots of four genes showing (i) negatively associated ESV with developmental time in whole embryo (*Myl9*), (ii) positively associated ESV with developmental time in whole embryo (*Rab3a*), (iii) negatively associated ESV in brain with developmental time (*Rspo2*), and (iv) positively associated ESV in brain with developmental time (*Grin2b*). (f). Average stage-corrected ESV within brain regions at each stage among genes with nominally significantly (P < 0.05) negative (light green) and positive (dark green) associations between spatial variability and developmental progression. (g). Examples of genes showing divergent trends in ESV between brain and non-brain across developmental time. Heatmap shows corrected ESV within brain and non-brain at each stage. Rows are shaded by temporal patterns: light green demonstrates negative ESV trend, dark green demonstrates positive ESV trend, grey demonstrates consistently high ESV throughout development. (h). Proportion of differentially expressed (DE) genes in each tissue among genes which are SVGs within non-brain (top) and brain (bottom) at least 6 of the 8 stages. We rank the tissues based on the decreasing values of the proportion. (i). Disease informativeness of genes with negative (circles) or positive (triangles) temporal ESV associations in brain (top) and non-brain (bottom), focusing on four brain-related Mendelian disorders. Informativeness is measured by the fraction of disorder genes with significant ESV trends (X axis) and the odds ratio between gene sets (Y axis). (j). Spatial expression plots of two monogenic autism risk genes, *Nrxn1* and *Foxp2*, showing different spatial patterns within brain and non-brain. *Nrxn1* exhibits positive ESV trend with development in both brain and non-brain, whereas *Foxp2* shows positive ESV trend specifically in brain. Numerical results are reported in **Supplementary Table S5**.

In the whole-embryo analysis, we expectedly observed a progressive increase in the number of SVGs across developmental stages. For example, Spacelink identified 2.05x more SVGs in E16.5 stage compared to E9.5 stage (**Figure 5b**). Similar patterns were observed in non-brain regions, however, notably for brain, the number of SVGs peaked at the E12.5 stage (**Figure 5b, Supplementary Figure S13a)**. To quantify these dynamics at the gene level, we fit a regression model relating stage-corrected ESV values against developmental time (**Methods**) and contrasted it with a control model associating overall mean and variance of sc-transform normalized gene expressions^73^ across time. The whole-embryo analysis identified 244 and 51 genes exhibiting (i) nominally significantly positive and negative associations between spatial variability and developmental progression (p_ESV < 0.05), and (ii) not showing significant change in expression over time (p_expression > 0.2) (**Figure 5c, Supplementary Figure S13b**). Pathway enrichment analysis using ConsensusPathdb^74^ showed that negatively ESV associated genes were enriched in muscle contraction pathways, while positively ESV associated genes were linked to later developmental processes such as Rab regulation of trafficking (**Figure 5d**). In brain regions, 104 and 41 genes showed positive and negative associations between spatial variability and development, also without significant expression shifts (p_ESV < 0.05, p_expression > 0.2) (**Figure 5c, Supplementary Figure S13b**). 53% and 43% of ESV-associated genes in whole embryo and brain were not identified when associating overall (spatial + non-spatial) variance of normalized expression against developmental time, underscoring the distinct informativeness of spatial variability beyond overall expression variability (**Supplementary Figure S13c)**. Negatively ESV-associated genes in brain were enriched in Wnt signaling, whereas positively associated genes involved Rap1 signaling (**Figure 5d**). The Wnt signaling pathway is well known to play a critical role in early vertebrate development by controlling the anterior-posterior axis formation and neural plate patterning, subsequently leading to the formation of neural crest^75–77^. Wnt signaling is not among the top 5 pathways (based on p-value < 0.05) when associating mean or variance of log-normalized expression in brain instead of ESV against developmental time (**Supplementary Figure S13d**), highlighting the utility of spatial variability metric. **Figure 5e** highlights representative genes from the aforementioned pathways (*Myl9*: muscle contraction, *Rab3a*: Rab regulation of trafficking, *Rspo2*: Wnt signaling, *Grin2b*: Rap1 signaling), illustrating temporal increases or decreases in spatial variability in the whole embryo or brain regions. In the brain, we observed that genes with decreasing spatial variability along development exhibited a marked decline in ESV from E13.5, while genes with increasing spatial variability showed an upward trend beginning at E11.5 (**Figure 5f**). This opposing dynamic explains the observed peak in SVG counts at the E12.5 stage, as early and late developmental spatial programs appear to converge at this stage.

Next, we examined differences in spatial variability between brain and non-brain regions. **Figure 5g** illustrates representative genes with divergent spatial variability dynamics over developmental time in the two different regions. In total, 9,481 and 16,566 genes were identified as SVGs in brain and non-brain, respectively, in at least 6 of the 8 developmental stages. 982 genes showed divergent trends in ESV between brain and non-brain across the developmental time (**Methods, Supplementary Table S5**; these include key developmental transcription factor genes like *Dlx6/Dlx5, Dmrt3*, and *Hoxd3. Dlx6* and *Dlx5* showed increasing ESV across development in brain (r=0.89 and 0.71, respectively) but decreasing ESV in non-brain regions (r=-0.78 for both). *Dlx6* and *Dlx5* are homeobox genes that are critical to forebrain development and to the specification and differentiation of inhibitory neurons^78,79^. *Dmrt3* shows a decreasing trend in ESV across development in brain (r=-0.90) but an increasing trend in non-brain (r=0.91); *Dmrt3* plays an essential role in spinal cord development by controlling the differentiation of interneuron subtypes involved in locomotor circuits^80,81^. The gene *Hoxd3*, on the other hand, shows a decreasing trend in ESV across development in non-brain (r=-0.94) but does not show a noticeable trend in brain (r=-0.10). *Hoxd3* is a member of the Hox gene family that governs anterior– posterior patterning and maintains cell fate and identity^82,83^.

We assessed the overlap of Spacelink SVG gene programs (P < 0.05) in brain and non-brain with differentially expressed (DE) genes in each tissue; in both cases, we see high overlaps with DE genes in tissues like spinal cord, dorsal root ganglion, and choroid plexus (**Figure 5h**). We next evaluated the odds ratio of genes with developmental time–associated ESV in brain and non-brain regions against gene sets corresponding to 4 brain-related Mendelian disorders^84,85^. Monogenic autism showed a notably stronger excess overlap with genes exhibiting increasing spatial variability in brain compared to those with decreasing variability (OR = 4.05 vs 1.44) (**Figure 5i**). By contrast, overlap with ESV-associated genes in non-brain was comparatively modest (OR = 1.70 and 1.04 for positively and negatively associated genes, respectively). These findings suggest that spatial processes emerging during later stages of brain development (E12.5 and beyond; **Figure 5f**) are specifically important to monogenic autism. On the other hand, genes associated with developmental Mendelian disorders, as well as monogenic psychiatric and neurological disorders, showed relatively weak excess overlap with genes displaying temporal trends in spatial variability in both brain and non-brain regions (**Figure 5i**). We highlight *Nrxn1* and *Foxp2*, both monogenic autism genes, that exhibit increased spatial variability in late developmental stages in the brain while displaying distinct spatial patterns in non-brain regions (**Figure 5j**).

Together, these results highlight that Spacelink can uncover dynamic spatial variability programs during organogenesis that are uniquely informative for monogenic autism and other neurodevelopmental disorders.

### Spacelink can characterize spatiotemporal dynamics of mouse in-vivo perturbation programs

Building on our finding that spatial variability during brain development is particularly informative for monogenic autism, we next examined how perturbations of de novo autism risk genes propagate through spatial programs in the developing brain. To this end, we analyzed an in-vivo Perturb-seq data^86^ targeting 35 de novo loss-of-function (LoF) risk genes associated with autism spectrum disorder (ASD) and neurodevelopmental delay (ND)^87,88^, where a gene was knocked out (CRISPR-ko) at the E12.5 stage, followed by single-cell RNA-seq readout at the P7 stage (**Figure 6a**). Among downstream-altered genes (which we term as “perturbation program” following Geiger-Schuller*, Eraslan* et al.^89^) identified across perturbations in five brain cell types—excitatory neurons (Exc), inhibitory neurons (Inh), astroglia, microglia, and oligodendrocytes, we observed strong concordance between astroglia and excitatory neurons (OR = 14.6), compared to an average OR of 2.3 among other cell type pairs (**Figure 6b**). This likely reflects either (i) activation of radial glia within the astroglial compartment toward neurogenic fates upon perturbation^90,91^, or (ii) perturbation-induced activation of gene programs underlying astrocyte–glutamatergic neuron interactions^92,93^. Overall, we observed a moderate to high concordance in downstream-altered genes across cell types among the 35 autism KO genes (average OR = 5.0) (**Supplementary Figure S14)**. We then applied Spacelink to map spatial variability for each perturbed gene and its downstream-altered genes using the spatiotemporal Stereo-seq mouse organogenesis data from **Figure 5**. Of the 32 perturbed genes present in the mouse organogenesis dataset, 26 were classified as spatially variable in all 8 developmental stages (mean ESV per gene ranging from 0.07 to 0.80; **Supplementary Figure S15a**). Downstream-altered genes in astroglia and excitatory neurons exhibited, on average, 1.76x higher ESV values than the perturbed genes, whereas downstream genes in other cell types (inhibitory neurons, microglia, oligodendrocytes) showed similar ESV levels (to their perturbed genes (**Figure 6c, Supplementary Figure S15b**). Control analyses in two external datasets—(i) genome-wide Perturb-seq in K562 cells^94^ and (ii) transcription factor overexpression in embryonic stem cells^95^ both showed a 40% and 59% lower average ESV in their downstream perturbation programs compared to the in-vivo perturbation programs in astroglia and excitatory neurons (**Figure 6c**). This highlights that perturbation of autism risk genes preferentially amplifies spatially organized downstream gene programs, specifically within astroglia and excitatory neurons. Next, we investigated genes that were differentially affected downstream of a given perturbation exclusively in top 3 cell type pairs exhibiting the strongest odds ratio in **Figure 6b:** astrocyte-Exc, microglia-Exc, and astrocyte-microglia. Genes altered in both astroglia and excitatory neurons, but not microglia, showed consistently high stage-corrected ESV across developmental stages (**Supplementary Figure S15c**). In contrast, cell type pairs involving microglia exhibited a gradual increase in ESV from E9.5 to E16.5 stages (**Supplementary Figure S15c**), likely reflecting the progressive intrusion of microglia that typically begins around E9.5 and continues till approximately E14.5 stages^96^. Across the 8 developmental stages, downstream-altered genes from in-vivo Perturb-seq assay showed 10-20% stronger correlation between brain-specific and non-brain-specific ESV values compared to all genes; this improvement remained consistent when restricting the downstream-altered genes to astrocytes and excitatory neurons (**Figure 6d**). Substituting ESV with mean and overall variance of log-normalized expression abolished or inverted these trends (**Supplementary Figure S15d**), emphasizing the unique insight provided by spatial variability. Next, we assessed the association of the ESV of each downstream-altered gene against (i) the number of perturbations affecting the given gene, and (ii) the number of cell types in which we see the gene with altered expression; in both cases, ESV increased significantly with these factors (**Figure 6e**). Among different cell types, the strongest increase in ESV with the number of upstream perturbations was observed in inhibitory neurons (**Supplementary Figure S15e**).

**Figure 6.**
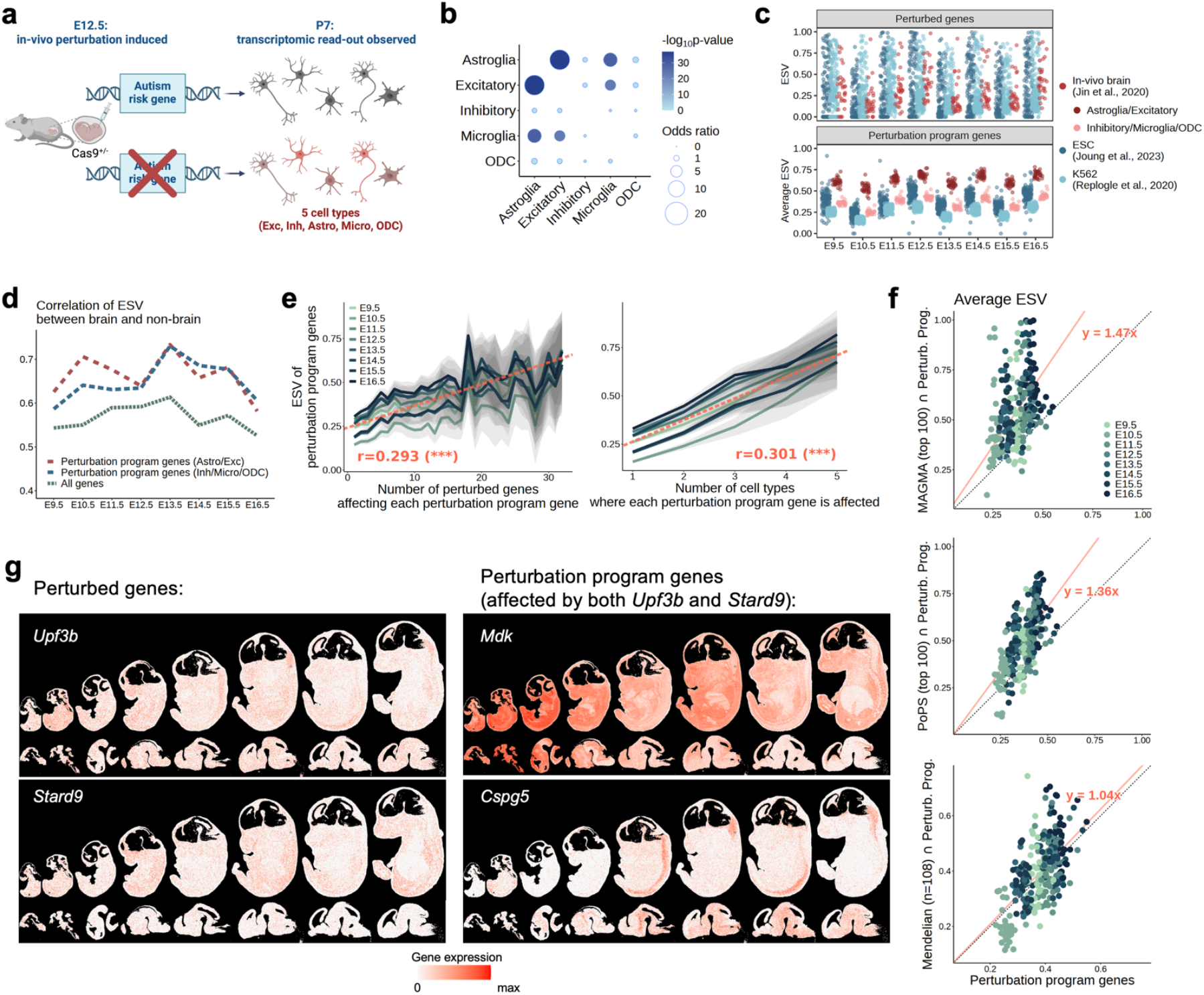
Spacelink characterizes spatiotemporal dynamics of downstream programs in mouse in vivo perturbations of autism risk genes. (a). Schematic of in-vivo Perturb-seq assay from Jin et al.^86^ targeting 35 de novo loss-of-function risk genes related to autism spectrum disorder (ASD) and neurodevelopmental delay (ND). Genes were knocked out (CRISPR-ko) at E12.5, followed by single-cell RNA-seq at the postnatal day 7 (P7) stage. Cells were broadly classified into five types—excitatory neurons (Exc), inhibitory neurons (Inh), astrocytes (Astro), microglia (Micro), and oligodendrocytes (ODC)—and downstream-altered genes were identified per perturbation within each cell type. (b). Odds ratios of excess overlap between downstream altered “perturbation program” genes across perturbations for each pair of cell types. Magnitude (odds ratio, dot size) and significance (−log_10_(p-value), dot color) of the excess overlap. (c). (Top panel): Effective Spatial Variability (ESV) of perturbed genes derived from three Perturb-seq experiments across mouse organogenesis stages. These perturbation experiments include our focal in-vivo Perturb-seq assay in brain^86^, as well as two external Perturb-seq assays - genome-wide Perturb-seq in K562 cells^94^, and transcription factor overexpression in embryonic stem cells^95^. Each dot corresponds to a perturbed gene in one of these assays. The points are colored by the underlying perturbation experiment. (Bottom panel): Average ESV of the perturbation program genes for each perturbation. For the in-vivo Perturb-seq, we grouped perturbation programs into two groups – genes altered downstream of a perturbation in astroglia and excitatory neurons (colored maroon), and genes altered downstream of a perturbation in microglia, oligodendrocytes and inhibitory neurons (colored pink). For the external datasets, given a single underlying cell type context, we consider the full set of altered genes in the perturbation program for each perturbation. (d). For each of the 8 mouse organogenesis stages, correlation between brains-specific and non-brain specific ESV across genes from three broad classes: union of perturbation programs of genes in excitatory neurons and astroglia, union of perturbation programs of genes in inhibitory neurons, microglia, and oligodendrocyte, and (iii) all genes. (e). ESV of downstream-altered genes (any cell type) across 8 organogenesis stages stratified by (left) number of perturbed genes affecting them and (right) number of affected cell types. Thick lines denote average ESV; shaded areas show confidence bands. Red dashed line indicates the overall trend estimated by fitting a linear regression model. Asterisks denote significance of Pearson correlation (*** P < 0.001). (f). Scatter plots comparing average ESV of downstream-altered genes (perturbation program) per perturbation at each stage for all genes versus genes intersecting with three autism-associated gene sets: (i) top 100 MAGMA-prioritized polygenic genes (top), top 100 PoPS-prioritized polygenic genes (middle), and (iii) 108 monogenic autism risk genes from OMIM^84^ by Freund et al.^85^ (bottom). Red lines indicate fitted regression line through the scatter plot based on a linear regression model with zero intercept. We report the slope of the regression line. (g). Spatial expression plots of two de novo autism risk genes, Upf3b and Stard9 across mouse organogenesis stages (left panel), showing minimal spatial variability patterns, and two downstream-altered genes for both these genes (right panel), showing distinct spatial variability patterns in brain and non-brain. Numerical results are reported in **Supplementary Table S6**.

To assess the polygenic and monogenic disease relevance of spatial variability in perturbation programs from de novo autism risk gene perturbations, we calculated the average ESV of program genes at each developmental stage of mouse organogenesis stratified by three classes of autism risk genes - (i) the top 100 MAGMA-prioritized GWAS genes^30,31^, (ii) the top 100 PoPS-prioritized genes (integrating GWAS and gene-level functional features)^29^, and (iii) 108 monogenic autism Mendelian disorder genes^85^. Perturbation program genes overlapping the MAGMA and PoPS sets exhibited markedly higher average ESV (1.47x and 1.36x, respectively) compared to all downstream-altered genes, whereas overlap with Mendelian genes showed no such enrichment (**Figure 6f**). Similarly, relative to the top 100 MAGMA and PoPS gene sets overall, perturbation program genes overlapping these disease gene sets showed stronger ESV (2.43x and 1.75x, respectively; **Supplementary Figure S16a**). These enrichments diminished for both MAGMA and PoPS disease gene sets when ESV was replaced by the mean of normalized gene expression (**Supplementary Figure S16b)**. Replacing the ESV with overall variance (spatial + non-spatial) resulted in diminished enrichments for MAGMA but showed comparable enrichments for PoPS (**Supplementary Figure S16c**). Together, these findings suggest that perturbing high-effect de novo variants tend to preferably impact spatially variable genes linked to common polygenic autism risk loci. Additionally, perturbation program genes overlapping MAGMA were altered by a greater number of perturbations than those implicated by PoPS or Mendelian autism, with microglial programs showing the strongest associations (**Supplementary Figure S16d**). In **Figure 6g**, we highlight two example de-novo autism risk genes, *Upf3b* and *Stard9*, which themselves exhibit low spatial variability in both brain and non-brain regions but influence multiple MAGMA-prioritized autism GWAS genes (Z-score > 3) showing distinct spatiotemporal trends. One such example is *Mdk*, a downstream target of *Upf3b* and *Stard9* in inhibitory neurons and oligodendrocytes, which shows high spatial variability in early development in brain and non-brain. This is consistent with prior studies demonstrating that Midkines (*Mdk)* are strongly expressed in early emryonic development where it assists in neural development, but shows decline in expression in later development stages and in adult tissues^97,98^. Another example, *Cspg5*, a downstream target of *Upf3b* and *Stard9* in microglia and oligodendrocytes, exhibits increased spatial variability in later stages.

We conclude that integrating in-vivo Perturb-seq with Spacelink analysis of spatiotemporal transcriptomic maps enables tracing how genetic disruptions potentially propagate spatially across developmental stages, uncovering convergent spatial programs underlying neurodevelopmental disorders.

### Spacelink characterizes spatially variable genes programs underlying neurodegeneration

To extend our analysis from development to aging and neurodegeneration, we applied Spacelink to 32 ROSMAP 10x Visium samples from brain dorsolateral prefrontal cortex (DLPFC) (4 control samples from 2 healthy donors and 28 samples from 15 donors at different stages of Alzheimer’s disease (AD)) (see **Data Availability**), each with detailed global pathological annotations (N =1,166–2,649 spots, 36,601 genes) (**Figure 7a**). Of the pathological variables, we focused on 3 hallmark indicators of AD progression - the total amyloid plaque burden^99^ (square-root transformed), neurofibrillary tangles accumulation (tau)^100^ (square-root transformed), and the number of neuritic plaques (square-root transformed). We applied Spacelink to each sample at both whole tissue and cortical layer-level. We did not perform Spacelink cell-type-specific analyses, as disease progression is accompanied by substantial heterogeneity within cell types—such as the emergence of disease-associated microglia^101,102^—rendering cell type annotations from static brain cortex snRNA-seq reference-based deconvolution methods like RCTD^23^ less reliable. Unlike mouse organogenesis data, the spatial cross sections across samples had similar sizes, leading to consistent length scale initialization (**Supplementary Figure S17a)**, thereby precluding the need to correct ESV scores for size differences.

**Figure 7.**
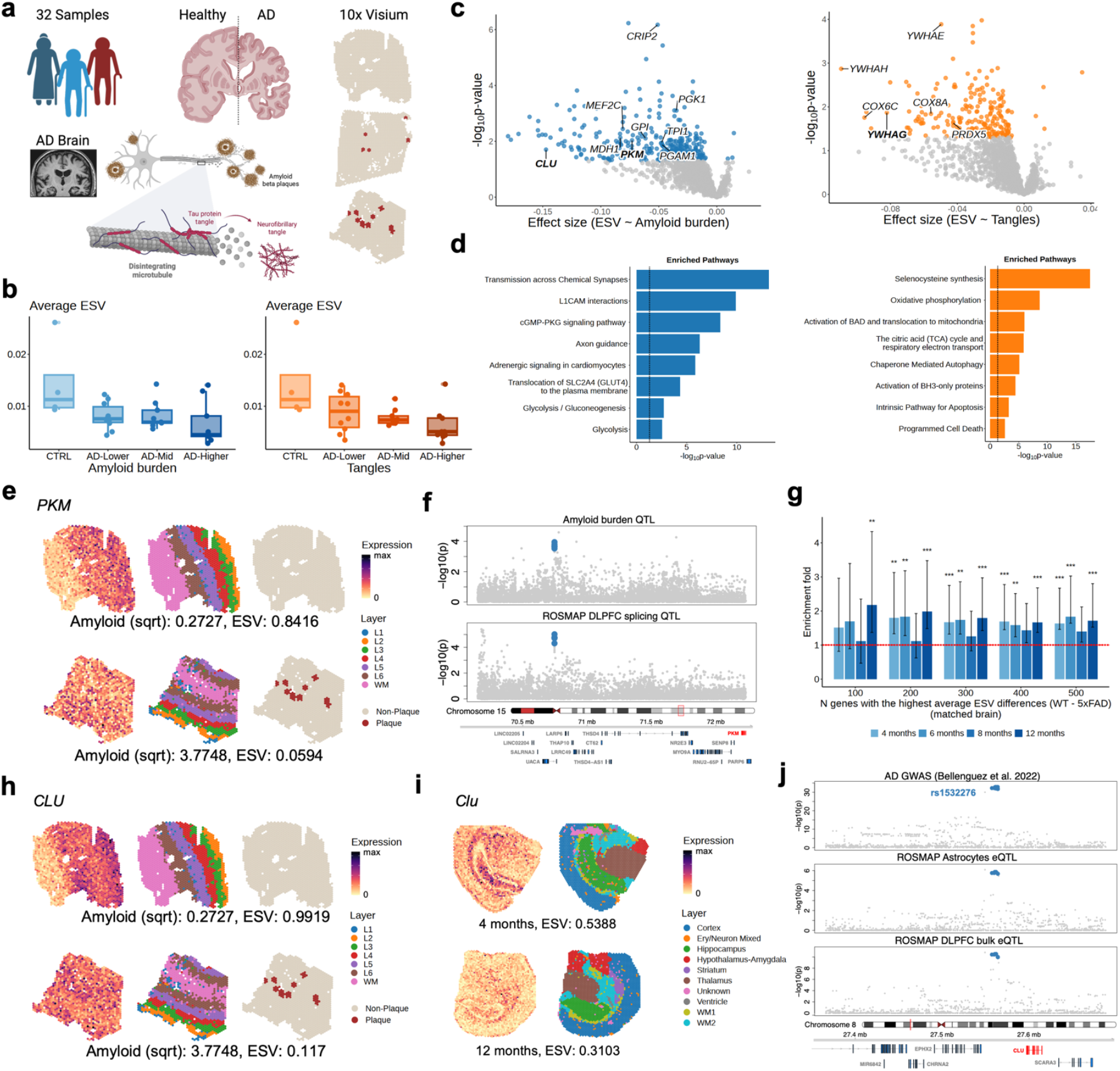
Spacelink captures dynamic alterations in spatial variability across Alzheimer’s disease progression. (a). Illustration of 10x Visium human brain dorsolateral prefrontal cortex (DLPFC) data from 4 control samples from 2 healthy donors and 28 samples from 15 donors at different stages of Alzheimer’s disease (AD), with comprehensive pathological metadata available for each sample, including total amyloid burden, neurofibrillary tangle accumulation, and neuritic plaque counts. Amyloid burden, tangle accumulation, and neuritic plaque counts were square-root transformed to facilitate regression analyses. (b). Boxplots of average Effective Spatial Variability (ESV) scores across genes at control and 3 different stages of AD pathology, determined by levels of amyloid burden (left) and tangle accumulation (right). For each measure, CTRL includes 4 control samples from 2 healthy donors; AD-Lower includes 10 samples with the lowest values of the pathology indicator; AD-Mid includes 9 samples with intermediate values of the pathology indicator; and AD-Higher includes 9 samples with the highest values of the pathology indicator. (c). Volcano plots representing regression analyses associating ESV with amyloid burden (left) or tangle accumulation (right) at the gene level, adjusting for cortical layer composition and mean expression levels, and donor-level random effects. Each point represents the slope coefficient from the regression model and its –log p-value for a gene. We highlight genes that are nominally significant (P < 0.05) for association against amyloid (colored blue) and tau burden (colored orange) respectively. (d). Pathway enrichment analyses of 334 and 216 genes with nominally significant (P < 0.05) negative associations between spatial variability (ESV) and amyloid burden (left) or tangle accumulation (right). Pathway enrichment analysis was performed using ConsensusPathdb^74^ with the set of genes in the Visium panel as the background set. (e).Spatial expression plots of the *PKM* gene (left column), along with layer annotations (middle column), and plaque annotations (right column) in one control sample (top panel) and one high–amyloid-burden sample (bottom panel). (f). Pathology-xQTL colocalization analysis using ColocBoost identifies a 95% colocalized confidence set (chr15: 70531387-70531393) consisting of 5 variants, showing colocalized effect between a splicing QTL phenotype in brain DLPFC and a square-root transformed amyloid burden QTL in an independent ROSMAP cohort with N=595 individuals. (g).Excess overlap analysis (see **Methods**) of genes with negative associations between ESV and amyloid burden (from panel c), among genes with the largest average ESV differences between WT and 5xFAD mouse in 10x Visium SRT data from the 5xFAD mouse model at four time points with progressive amyloid accumulation. Asterisks denote significant enrichment by Fisher’s exact test (*** BH-adjusted P < 0.001, ** BH-adjusted P < 0.01, * BH-adjusted P < 0.05). (h) Spatial expression plots of the *CLU* gene (left column), along with layer annotations (middle column), and plaque annotations (right column) in one control sample (top panel) and one high–amyloid-burden sample (bottom panel) from 10x Visium human DLPFC data. (i) Spatial expression plots of the *Clu* gene (left column), along with layer annotations (right column) from 10x Visium mouse SRT data of 4-month-old (top panel) and 12-month-old (bottom panel) 5xFAD mice. (j). Alzheimer’s disease (AD)-xQTL colocalization analysis with CLU, showing a 95% colocalized confidence set (chr8: 27604964-27610304) consisting of 8 variants between a CLU eQTL specifically in astrocytes, together with bulk eQTL in DLPFC, and AD GWAS. Numerical results are reported in **Supplementary Table S7**.

Spacelink revealed a consistent decline in both the number of spatially variable genes (SVGs) and their average ESV scores with increasing amyloid burden, neurofibrillary tangle accumulation, and neuritic plaque counts, indicating a progressive loss of spatial transcriptional organization during neurodegeneration (**Figure 7b, Supplementary Figure S17b**). Across genes, average ESV scores were negatively associated with all AD pathology indicators, even after adjusting for donor-level random effects (**Methods**; **Supplementary Figure S17b**). To systematically identify genes whose spatial variability is linked to AD pathology, we performed regression analyses relating gene-level ESVs to amyloid and tau burdens and neuritic plaque counts, while adjusting for cortical layer composition and mean expression levels (**Methods**). We identified 334, 216, and 115 genes whose ESVs were significantly negatively associated with amyloid and tau burdens and neuritic plaque counts, respectively (p < 0.05; **Figure 7c, Supplementary Figure S17c**). Pathway enrichment analysis of amyloid-associated SVGs revealed glycolysis and gluconeogenesis as the most enriched pathways (**Figure 7d**), consistent with metabolic dysfunction in early AD. In contrast, genes associated with tangle accumulation were enriched for apoptotic pathways, highlighting neuronal loss and programmed cell death in later disease stages (**Figure 7d**). Genes associated with neuritic plaque counts were enriched for metal ion homeostasis pathways that are known to manage amyloid plaque toxicity^103^ (**Supplementary Figure S17d**). We illustrate spatial patterns for two representative genes: *PKM*, associated with amyloid burden, *YWHAG*, associated with tau accumulation, and *SNCB*, associated with neuritic plaque counts (**Figure 7e, Supplementary Figure S17e, Supplementary Figure S17f**). *PKM*, a key glycolysis enzyme^104^, showed colocalization between a splicing QTL in brain DLPFC and an amyloid burden QTL in an independent ROSMAP cohort of 595 individuals^105^ (**Figure 7f**). In contrast to our cross-gene disease association regression model in **Figure 4**, we did not include overall variance as a covariate in our primary cross-sample pathology regression. Whereas **Figure 4** focused on assessing the unique disease informativeness of entire spatial programs, here our goal was to identify individual genes whose spatial variability is associated with AD pathology. Because overall variance reflects both spatial and non-spatial factors, including it as a covariate risk overcorrecting true SVG signals and reducing power. Nonetheless, we performed a secondary analysis that incorporated overall variance as an additional covariate. The resulting model identified 260, 166, and 103 genes as ESV-associated with amyloid and tau burdens and neuritic plaque counts, respectively, which were enriched in glycolysis, apoptotic, and metal ion homeostasis pathways (**Supplementary Figure S17g, Supplementary Figure S17h**). Finally, in comparing with plaque-proximal differential expression reported by Karaahmet et al.^28^, 25.6% (n=10) of their 39 genes showed significant associations between ESV and at least one of the 3 pathological variables, demonstrating a strong excess overlap between the two classes (19.1x; **Methods**). However, the finding that a large fraction (74.4%) of plaque-proximal genes did not exhibit pathology-associated spatial variability underscores that these two spatial gene classes capture complementary aspects of AD-related pathology—one reflecting broad spatial variability and the other localized responses to plaques.

Next, we examined mean expression changes associated with AD pathology while adjusting for cortical layer composition (**Methods**). We identified 1,051 and 745 genes whose mean expression levels were significantly negatively associated with tau burden and neuritic plaque counts, respectively (p < 0.05). Only 2 genes showed significant association with amyloid burden (**Supplementary Figure S18a**). The significant genes were enriched in multiple broad neurodegeneration-related pathways, which were distinct from those enriched by ESV-associated genes (**Supplementary Figure S18b**). Next, we applied Spacelink to each of the 6 cortical layers (L1-L6) and white matter (WM) separately, and tested association between layer-specific average ESV and AD pathology markers. We observed nominally significant (p < 0.05) reduction in average ESV with tau accumulation in L6 cortical neurons (**Supplementary Figure S18c**). This is consistent with the fact that tau pathology predominantly affects deep cortical layers such as L6, which are more vulnerable in AD^106,107^. However, we observed less pronounced reduction in layer-specific spatial variability compared with global spatial variability for all pathological measures, suggesting that layer-specific spatial architecture is relatively more preserved during AD progression (**Figure 7b, Supplementary Figure S17b, Supplementary Figure S18c**).

To evaluate the cross-species robustness of spatial variability patterns identified in aging human brains with AD, we analyzed 10x Visium spatial transcriptomics data from 5xFAD transgenic mouse model^6^, an established model of early-onset familial AD characterized by progressive amyloid plaque deposition^108^. The dataset encompasses 80 samples spanning four time points (4, 6, 8, and 12 months), enabling assessment of spatial variability dynamics along the trajectory of amyloid accumulation. We applied Spacelink to 5xFAD samples from all time points, as well as wild-type (WT) samples, and evaluated the spatial variability patterns of the 334 human amyloid-associated genes observed in **Figure 7c**. We restricted the Spacelink analysis in mice to the matched cortical layers as AD human brain, and examined differences in average ESV between 5xFAD and WT samples at each time point. We observed higher enrichment of human amyloid-associated ESV genes among the genes with the highest average ESV differences (WT – 5xFAD) at 12 months, the final disease stage, compared to earlier stages with lower amyloid deposition (**Figure 7g**). This is consistent with the human data, where these 334 genes also exhibited reduced spatial variability with increasing amyloid burden. One example is *CLU*, a known AD risk gene^109,110^, which showed 8.48-fold and 1.74-fold lower ESV in the late-stage AD human brain and 12-month-old 5xFAD mouse, respectively, compared to control human samples and early stage mouse 5xFAD samples (4 months) (**Figure 7h, Figure 7i**). Notably, *CLU* is enriched in expression in astrocytes, and we observed a colocalization between *CLU* eQTL and AD GWAS specifically in astrocyte cell type^105^ (**Figure 7j**); astrocytes are known to play an important role in accumulation of glucose in the vicinity of the amyloid plaques. Other AD risk genes showing conserved spatial variability differences based on amyloid burden include *PGK1* and *GPI* (**Supplementary Figure S18d**). *PGK1* is known to support neuronal energy metabolism, and *GPI* is known to act as both a glycolytic enzyme and a neurotrophic factor^111,112^. Some AD risk genes showed distinct ESV trends against amyloid deposition in human and mouse, likely highlighting divergent spatial programs between human AD and mouse 5xFAD models. An example gene is *MEF2C* which showed a decline in spatial variability in human data but an increasing trend in the 5xFAD mouse model (**Supplementary Figure S18e**).

We conclude that Spacelink effectively captures disruption in spatial layered organization of gene expression linked to key pathological features of Alzheimer’s disease—including amyloid and tau burden—and reveals evolutionarily conserved spatial programs between human and mouse models of AD.

## Discussion

We present Spacelink, an adaptive multi-kernel framework for detecting and prioritizing spatially variable genes (SVGs) at both whole-tissue and cell-type resolution. Spacelink introduces three key methodological advances over existing approaches: (i) data-driven estimation of multiple spatial length scales to more accurately model the spatial variance component of each gene; (ii) a rule-based gating strategy for cell-type–specific analysis that corrects for colocalization of weakly represented cell types with other more dominant cell types in mixed spots in lower resolution Visium data, improving both sensitivity and specificity; and (iii) the Effective Spatial Variability (ESV) metric, which integrates spatial length scales and their associated variance components into a single interpretable score. Applied across diverse datasets, Spacelink revealed that spatial variability carries distinctive biological and disease-relevant signals that are not captured by traditional expression-based measures. Spacelink is also able to effectively quantify dynamic changes in spatial variability in organogenesis and tissue degeneration, uncovering regulatory pathways that shift in spatial tissue organization. Finally, Spacelink-derived variability measures provide insights into the preferential spatial activity of gene programs downstream of in vivo perturbations based on cell type context and disease relevance.

We note that the ESV metric provides a unified summary of spatial heterogeneity by integrating contributions across multiple spatial scales without making assumptions regarding specific spatial patterns for a gene. In practice, spatial variability in gene expression can arise from diverse processes, including enrichment within spatially defined niches or domains, anatomical gradients, layered structures, and clustered hotspots. By collapsing these heterogeneous signals into a single robust metric, ESV enables consistent prioritization of genes across tissues, datasets, and platforms. Importantly, ESV should not be viewed as an endpoint but more as a prioritization tool: genes with high ESV can subsequently be interrogated with secondary analyses designed to uncover the distinct spatial processes underlying their expression profiles. While previous studies have established SVGs as valuable molecular markers for tissue organization and disease progression^113,114^, no consensus has emerged on the optimal metric to quantify spatial variability or its general informativeness across disease contexts. Our results demonstrate that ESV consistently outperforms existing approaches in both robustness and disease relevance, underscoring its utility in connecting spatial tissue architecture to human disease genetics.

Our work has several downstream applications. First, Spacelink ESV can be used to pre-filter or weight genes while performing unsupervised clustering or deconvolution of spatial data^115,116^ and pre-select ligand and receptor pairs for spatial spatial cell–cell interaction inference^117,118^, thereby focusing these analyses on genes with strong spatially variable signals. Second, Spacelink tissue-level and cell-type level ESV gene scores can be used as features to inform gene-based disease association statistics^29,119^ and gene-based probabilistic fine-mapping of transcriptome-wide association studies^120^.Third, Spacelink’s decomposition of spatial variance into multiple length scales enables of genes by their spatial bandwidth profiles across the tissue, which can provide insights into pathway-level spatial organization. Fourth, the relative stability of the Spacelink ESV metric across platforms and conditions demonstrates its utility in comparative analyses across individuals and cohorts, including association with pathology profiles and genetic variation, thereby facilitating spatial quantitative trait loci (QTL) mapping.

Our work has several limitations, representing important directions of future research. First, Spacelink prioritizes genes based on overall spatial variability at whole-tissue and cell type resolution, but it is not designed to capture local spatial autocorrelation, as implemented in Hotspot^121^, or model spatial cell-cell interactions^117,118,122^. As such, Spacelink’s gene prioritization should be viewed as complementary rather than exhaustive in characterizing spatially informative gene programs within a tissue. Second, even with the relative cross-platform stability of ESV over other spatial metrics, residual differences in capture efficiency, spot density, and gene detection rates across platforms and assays can influence downstream comparisons, such as meta-analyses. Third, our analyses primarily focus on platforms profiling the full transcriptome or large gene panels (>1,000 genes), which facilitate genome-wide disease association analysis by providing a sufficiently large set of control genes. To this end, we did not consider highly customized smaller gene panels that are often strongly ascertained based on disease relevance^19,123,124^. Fourth, although Spacelink decomposes variance into multiple spatial length scales, linking these scales to exact biological structures and mechanisms is non-trivial and may require additional histological support. To this end, pathway enrichment analysis of groups of genes with similar spatial scales can inform about the underlying mechanisms. Fifth, the cell-type–specific version of Spacelink relies on accurate spot-level cell type proportion estimates from deconvolution methods such as RCTD^23^, and errors from poorly matched reference data or cell type annotations can propagate into cell type ESV inference. For this reason, we do not recommend the cell-type version of Spacelink for modeling spatial transcriptomic data from dynamic developmental or degenerative contexts when using static reference datasets, as cell type composition and states may change substantially over time. Sixth, our findings distinguish specific spatially variable programs of genes that play a greater role in genetic risk of disease but do not localize disease risk to a small number of genes, motivating more precise gene-level characterizations. Seventh, similar to other spatial transcriptomic approaches, cell-type-specific version of Spacelink applied to single-cell resolution SRT data will be affected by the cell segmentation quality, in particular, for less prevalent cell types. However, the extent of effect is less pronounced compared to local approaches, and the integration of spatial length scale into ESV score provides an additional safeguard against this artifact. Eighth, while our analyses highlight intriguing synergies between the spatial variability patterns monogenic and polygenic forms of autism, a systematic investigation across multiple polygenic diseases and their matched monogenic counterparts remains to be explored and represents an important direction for future research. Ninth, although we use donor-specific random effects in the AD neurodegeneration data, the way the cortical tissue section has been cut can potentially influence the spatial length scales and ESV, thereby acting as a confounder; While we typically observed robust laminar organization across samples and expect this effect to be modest relative to AD pathology, it remains an important technical consideration when interpreting spatial variability. Tenth, under high sparsity settings, Spacelink may produce an increased number of false positives in contrast to SPARK-X and SPARK. However, Spacelink offers important complementary advantages: it achieves superior statistical power and spatial variability ranking accuracy across multiple alternative simulations, greater cross-platform robustness, and stronger disease informativeness. For analyses of highly sparse spatial transcriptomic data, we recommend using SPARK-X for SVG hypothesis testing, while applying Spacelink or Spacelink-lite ESV to binned data as a complementary ranking metric for downstream disease integration analysis. Eleventh, Spacelink and most SVG methods lack the formal uncertainty quantification that fully Bayesian approaches provide through posterior distributions, making the latter preferable when rigorous credible intervals on spatial parameters are needed. While we do show improved benchmarking performance of Spacelink against two such approaches (spNNGP and spBayes) based on our metrics of interest, the field of spatial Gaussian Process (GP) models beyond SVG methods is huge^39–43^, and a comprehensive evaluation of Spacelink against those methods is beyond the scope of this work. Robustness to model misspecification is another theoretical strength of fully Bayesian approaches; the Gaussian likelihood approximation used in Spacelink may not be well-suited for such scenarios. However, the impact of this approximation appears modest across the simulation settings we have evaluated, except in high sparsity settings.

Despite all these limitations, Spacelink provides a scalable and interpretable framework for quantifying spatial variability across multiple biological scales, and within and across cell types, enabling robust integration with genetic variation underlying both monogenic and polygenic diseases, and offering a strong foundation for future advances in spatial omics technologies.

## Methods

### Overview of Spacelink

#### Spacelink global model

We denote **y**_g_ as the vector of normalized and transformed expression values for gene g across *N* spatial locations **s** = (**s**_1_, **s**_2_, …, **s**_*N*_). At the whole-tissue level, we model **y**_g_ with a data-driven multi-kernel Gaussian process as follows:

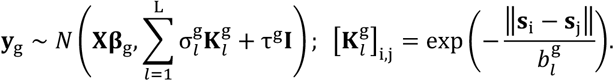

Here, **X** denotes covariates including the intercept and other global metadata at each spatial location; **β**_g_ the corresponding fixed effects; 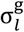 the spatial variance component for each exponential kernel 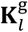 and τ^g^ the noise variance. Spacelink assumes that spatially variable genes can exhibit heterogenous spatial patterns with multiple bandwidths due to spatial localization of cell types, spatial zones or domain structure. To capture this, we use a set of exponential covariance kernels 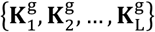, each with a different bandwidth 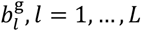 . This set can be viewed as a data-driven subset of a potentially infinite collection of kernels with varying bandwidths, which we term as the “*Kernel Store*”. Theoretically, any completely monotonic isotropic kernel can be approximated by a linear combination of exponential kernels in this store (**Supplementary Note**). Furthermore, as noted in previous work^8^, the exponential kernel is preferred over the Gaussian or Radial Basis Function (RBF) kernel because the squared exponential function in latter decays too rapidly with distance in spatial transcriptomics (SRT) data and is therefore less well-suited for capturing spatial variability across the range of length scales typically encountered in tissue data. We show through secondary simulation experiments (**Supplementary Figure S8a**) that replacing the exponential kernel with a RBF kernel, or adding broader varieties of kernels (Matern, periodic) over exponential kernels does not substantially alter the Spacelink hypothesis test results. Analogous to SpatialDE2 and STANCE, we use a mixture-kernel Gaussian process representation to model gene expression in Spacelink, rather than modeling expression as a mixture of Gaussians with different kernels. This is because the latter is computationally intensive and does not offer substantial advance when assessing global spatial variability, our primary focus, as opposed to local spatial variability.

We adopt a 2-stage selection mechanism to adaptively select kernels for each gene. We first pre-select 2*L* logarithmically spaced bandwidths ranging from the minimum to the maximum spatial distances across the *N* locations (with *L* = 5 as the default) (see schematic in **Supplementary Figure S1a**). We then fit the global model for a particular gene g with these candidate bandwidths using Non-Negative Least Squares (NNLS)^22^ and identify the smallest and largest bandwidths with non-zero weights (denoted 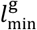 and 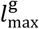 ). Finally, we define *L* log-spaced bandwidths within 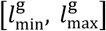, which specify the final set of spatial bandwidths 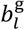 and corresponding set of kernels 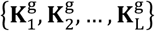 .

Spacelink assesses whether gene g is spatially variable by testing the null hypothesis 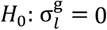 for all *l*, against alternative hypothesis given by 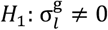 for some *l*. For each kernel *l*, we derive a test statistic of the form

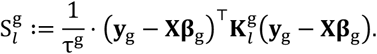

Under the null hypothesis, individual 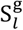 follows a mixture of chi-square distributions, which can be approximated by the Satterthwaite method to obtain a kernel-specific and gene-specific p-value 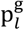. Since Spacelink employs multiple kernels per gene, we aggregate these kernel-level p-values into a single gene-level p-value using the Cauchy combination rule^21^, which is robust to arbitrary dependency structures among the individual p-values. **β**_g_ and τ^g^ in 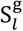 are estimated using maximum likelihood estimation under the null hypothesis (**Supplementary Note**). After obtaining the p-values for all genes, we adjust them using the Benjamini–Hochberg method. The adjusted p-value for each gene is then used to determine whether the gene is spatially variable.

Beyond hypothesis testing, we introduce a novel metric—*Effective Spatial Variability (ESV)*—which integrates variance components with corresponding kernel bandwidths to rank genes by their spatial patterning:

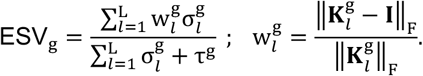

The weight 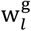 penalizes kernels with minimal spatial structure, thereby prioritizing genes with strong, non-trivial spatial variability. Existing metrics such as PropSV (σ^g^/(σ^g^ + τ^g^), nnSVG^8^) and FSV (σ^g^ w^g^/(σ^g^ · w^g^ + τ^g^), 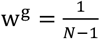, 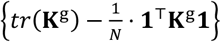 SpatialDE^9^) lack this property: PropSV applies no penalization for spatial bandwidth, while the weighting used in FSV does not effectively filter out genes with minimal spatial structure (e.g., w^g^ = 1 when **K**^g^ = **I** in FSV). ESV therefore provides a more interpretable and robust measure of spatial variability.

We provide several additional remarks on ESV: **1) Detailed properties of the ESV score:** ESV is jointly determined by two key factors—the spatial length scale *l* and the spatial scale-specific signal-to-noise ratio (SNR) 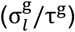 —and it increases with larger values of either parameter. To illustrate this more explicitly, we present how ESV varies with SNR and length scale in a single-kernel model (**Supplementary Figure S1c**); for comparison, the corresponding plots for PropSV and FSV under the same setting are shown alongside. Notably, PropSV and FSV are largely insensitive to changes in length scale and vary only with SNR. In practice, a gene with high ESV is strongly and consistently expressed across spatially extended regions; a gene with low ESV (but detected as an SVG) exhibits spatial structure confined to very short ranges, which may reflect biologically meaningful local enrichment but may also be susceptible to technical artifacts such as transcript spillover.

**2) Bounds on the ESV score:** The maximum value of ESV is 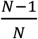, where *N* is the number of spots; in practice, with *N* > 1000, this ESV metric has an upper bound close to 1, so ESV can effectively be considered to be bounded between 0 and 1. ESV attains its maximum when the noise variance component τ^g^ = 0 and all spatial variance components 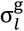 with non-zero contribution have length scales approaching infinity, so that each kernel matrix reduces to a matrix of ones 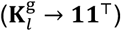 . In this limit, the weight 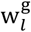 and consequently the entire ESV score approaches 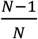 . Across the 10 scDesign3 simulation experiments in **Figure 2**, non-SVG genes typically have ESV < 0.05 (97.5–100% across datasets, **Supplementary Figure S8e**), and we therefore recommend treating ESV scores below this threshold with caution when interpreting spatial variability.

**3) Inclusion of** τ **in the ESV score:** The ESV score, as well as PropSV and FSV scores, quantifies the relative magnitude of 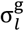 with respect to τ^g^, rather than its absolute magnitude. When 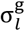’s are fixed, a larger τ^g^ (i.e., a high overall noise level) should correspond to lower spatial variability. The term 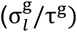 can therefore be viewed as a signal-to-noise ratio metric for a given spatial length scale. We provide an illustrative example in **Supplementary Figure S8f** highlighting this.

4) **Choice of Frobenius norm for the weights:** We use the Frobenius norm to quantify the difference between 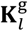 and **I** for its simplicity and computational efficiency. To verify that this choice does not drive our results, we compared performance using the 𝓁_1_ and spectral norms as alternatives across 10 scDesign3-based simulations; all three norms yielded nearly identical results (**Supplementary Figure S8d**), supporting the Frobenius norm as a robust default.

We estimate 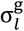 and τ^g^ by using a Non-Negative Least Squares (NNLS) fit of the empirical covariance matrix:

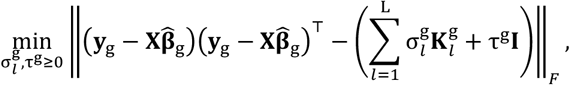

solvable with algorithms such as Lawson-Hanson^22^. Here, 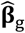 is estimated by least squares method.

While the original Spacelink is competitive with other fast global SVG methods for small to moderate datasets, its computational cost increases substantially for high-resolution spatial datasets involving hundreds of thousands of spots. To address the computational challenges posed by high-resolution data, we propose ***Spacelink-lite***. Instead of computing pairwise distances between all spots when constructing the kernel matrices, Spacelink-lite partitions the tissue into a grid and approximates distances between spots in different grid cells using the distances between the corresponding grid centers. This strategy substantially reduces both computational time and memory usage. For neighboring grid cells and for spots within the same grid cell, however, pairwise distances between spots are computed to preserve local spatial structure. We provide a schematic illustration of this in **Supplementary Figure S1b**. By default, the grid size is set to five times the minimum spot distance, but we recommend adjusting the grid size according to the characteristics of the dataset. For example, if the data are partially densely packed, resulting in a very small minimum spot distance, it may be advisable to use a larger grid size to avoid excessive computational burden.

Lastly, we note that no spatial coordinate transformation is required for our analyses, as the spatial bandwidth parameters in Spacelink automatically absorb differences in coordinate units across platforms. Intuitively, rescaling all spatial coordinates by a positive constant c simply rescales the estimated bandwidth by the same factor, leaving 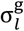, τ^g^, and the kernel weights unchanged, and consequently the ESV metric also stays unchanged (**Supplementary Note**). Users therefore do not need to preprocess spatial coordinates before running Spacelink, regardless of the platform used. To confirm this from a data application perspective, we applied Spacelink to simulated data generated by scDesign3 from the Visium human DLPFC dataset under two scenarios: (i) using the original spatial coordinates, and (ii) using min-max scaled coordinates as in nnSVG, defined as: 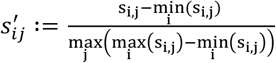 . We found that the estimates of 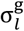 and τ^g^ were identical across all genes between the two scenarios, while the estimated length scales decreased proportionally in the scaled case — exactly as expected from the rescaling argument above — resulting in identical ESV values in both cases (**Supplementary Figure S8g**).

### Spacelink cell-type-specific model

Spatially variable genes at the whole tissue level are often enriched for cell-type markers due to the non-uniform spatial distribution of different cell types across the tissue. Therefore, it is often desirable to characterize spatial variability not only across the whole tissue, but also within individual cell types. In high-resolution spatial transcriptomics platforms such as CosMx, where each spot corresponds to a single cell with a cell-type annotation, we recommend applying the Spacelink global model to the subset of gene expression restricted to the cells of each cell type for assessing cell-type-specific spatial variability.

In contrast, in low-resolution spatial transcriptomics platforms such as 10x Visium, each spot aggregates transcripts from dozens of cells and may contain a mixture of several distinct cell types. In such cases, identifying and quantifying cell type SVGs becomes challenging, as a weakly representative cell type, like microglia in brain cortex, may heavily colocalize with another dominant cell type, like oligodendrocytes, in mixed spots, leading to confounded signals. For such cases, we propose a cell-type-specific extension of Spacelink that models spatial variability attributable to a focal cell type while accounting for potential confounding from spatial colocalization with other cell types. For a focal cell type r, Spacelink (ctSVG) models the full expression vector **y**_g_ as a linear mixed model.

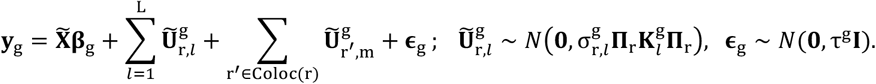

Here, 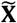 represents fixed effect covariates comprising the intercept, estimated cell-type proportions in spot as evaluated by the RCTD method^23^ as well as other spot-level metadata if available. **Π**_r._ represents an *N* × *N* diagonal matrix of the proportions of cell type r across spots. 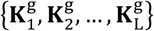 are the pre-selected kernels for gene g in the Spacelink global model (i.e., the kernels with the *L* log-spaced bandwidths within 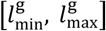 ), and 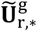 denote independent random effects capturing the projection of these kernels onto the cell type r. Coloc(r) denotes a set of cell types that colocalize with the focal cell type r, and our cell-type-specific Spacelink model conditions on the random effect terms 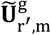 for these colocalizing cell types at the median spatial bandwidth m for gene g.

A key technical challenge in cell-type SVG modeling is in disentangling spatial random effects of the focal cell type from those of other colocalizing types. This colocalization may artificially induce correlated random effects across cell types and failing to correct for that may lead to false positives. On the other hand, naively including random effects for all cell types and length scales can lead to an overcrowding effect, where the model becomes overparameterized, diluting the power to detect true spatial signals in the focal cell type. To balance these issues, Spacelink (ctSVG) employs a data-driven gating procedure that defines the minimal set of colocalized cell types, Coloc(r), and corrects for their random effects at the median of the pre-selected bandwidths (i.e., the median of the *L* log-spaced bandwidths within 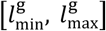) for gene g. Restricting correction to the median bandwidth, rather than all bandwidths, avoids excessive removal of true spatial signals from the focal cell type. The set Coloc(r) is determined as follows:

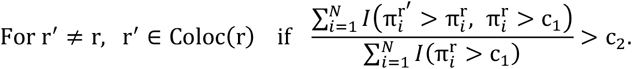

Here, 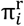 denotes the proportion of cell type r at the *i*-th spot. This rule identifies cell types that strongly colocalize with the focal cell type r, specifically, those that are more dominant than r (i.e., 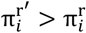 in more than a specific fraction (c_2_) of spots cell type r is moderately present (i.e., 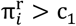). Intuitively, Coloc(r) attempts to identify “dominantly colocalizing” cell types – those that tend to outnumber the focal cell type in the spots where the focal type appears and are therefore most likely to confound its spatial signal if not accounted for. We note that colocalization under this definition is intentionally asymmetric: r^’^ ∈ Coloc(r) does not imply r ∈ Coloc(r^’^) . This asymmetry reflects a biologically meaningful distinction. For example, for brain cortex (DLPFC), oligodendrocyte cell type is included in Coloc(OPC) but OPC is not included in Coloc(Oligodendrocyte). OPCs are relatively rare in the brain cortex, and in spots where OPCs are present, mature oligodendrocytes tend to be co-present at substantially higher proportions (**Supplementary Figure S10a**). When modeling OPC-specific spatial variability, oligodendrocytes therefore need to be accounted for as a dominant colocalizing cell type, as their spatial signal would otherwise confound the OPC-specific inference. Conversely, in spots where oligodendrocytes are present, OPCs rarely outnumber them and including them as confounder may spuriously over-correct oligodendrocyte signal leading to reduced power. We use default cutoffs of c_1_ = 0.1 and c_2_ = 0.1. We evaluated alternative cutoff values in simulation studies and observed that performance was comparable to or worse than that obtained with the default settings (**Supplementary Figure S4g**), supporting the use of these defaults.

We assess whether a gene is spatially variable within cell type r by testing the null hypothesis 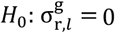 for all *l*. Under the null, **y** follows a normal distribution 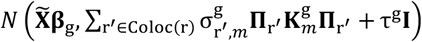 and we estimate the parameters **β**_g_, 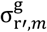 and τ^g^ via Restricted Maximum Likelihood (REML). Using these estimates, we compute the score test statistic for each kernel, obtain p-values via the Satterthwaite approximation, and then combine them with the Cauchy combination rule. The resulting combined p-value determines whether a gene is spatially variable within cell type r.

Analogous to *ESV*, we suggest cell-type-specific *ct-ESV* scores, by replacing 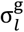 with 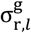, as follows:

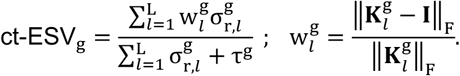

We estimate 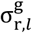 and τ^g^ by using Restricted Maximum Likelihood (REML).

We note that the current implementation of Spacelink, Spacelink-lite and Spacelink (ctSVG) supports multi-threaded execution on a single node and is naturally parallelizable across genes or gene subsets via HPC schedulers such as LSF or SLURM. Streaming-based processing is not currently supported, as the kernel matrix computation requires access to the full set of spatial coordinates.

### Simulation design for benchmarking global SVG methods

To evaluate global SVG detection methods, we designed three types of simulations: (i) scDesign3-based model^35^, (ii) covariance-based Gaussian process model, and (iii) model using custom-designed canonical spatial patterns. Following recent benchmarking work^34^, we first applied scDesign3 to generate biologically realistic data using 10 human 10x Visium samples as references. For each sample, we selected the top 200 SVGs based on Moran’s I^32^ and fitted their expression distributions using the `fit_marginal` and `fit_copula` functions in scDesign3. From these, we retained 50 genes with high deviance explained by the fitted model. For each gene, we obtained a mean expression parameter vector reflecting its spatial correlation (denoted ***μ***_s_) and generated a non-spatial counterpart (***μ***_*ns*_) by randomly shuffling ***μ***_*s*_. We then simulated data with varying spatial variability using weighted mixtures *α* · ***μ***_*s*_ + (1 − *α*) · ***μ***_*ns*_ as input to the `simu_new` function, with *α* ∈ {0.05, 0.1, …, 1}. Smaller *α* values correspond to weaker spatial variability. The *α* values were used to rank simulated genes, serving as the ground truth for evaluating SVG ranking methods. For each of the 50 genes, we generated 20 replicates across different *α* values, yielding 1,000 artificial SVGs. An equal number of non-SVGs were generated by shuffling the simulated SVG expressions. Across the 10 Visium datasets, this resulted in 2,000 simulated genes per dataset.

Next, we performed covariance-based simulations using a Gaussian process to model multi-scale spatial variance. We considered two versions: (i) a single-kernel covariance and (ii) a mixture of five kernels with different bandwidths. In the single-kernel version, the covariance was defined as

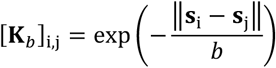

while in the five-kernel version, it was defined as

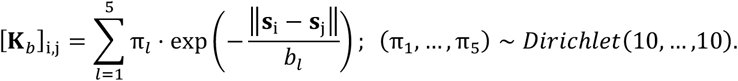

Gene expression count data (denoted **z**) were simulated using each type of covariance matrix as follows:

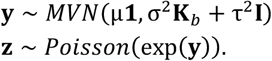

On the other hand, non-SVGs were simulated separately as described below.

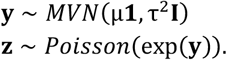

Spatial coordinates for all simulations were taken from the 10x Visium Human DLPFC dataset^25^ (N = 3,639). The parameter values used in the models were derived from nnSVG applied to the same dataset, which fits the model

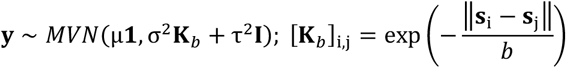

to normalized and log-transformed gene expression and provides estimates of µ, σ^2^, τ^2^ and *b* for each gene. For the simulation of SVGs, we first selected genes with adjusted p-values < 0.05 according to nnSVG. From these genes, we obtained the mean of µ estimates, the 10th, 30th, 50th, 70th, and 90th percentiles of 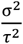 estimates, and the 10th, 30th, 50th, 70th, and 90th percentiles of τ^2^ estimates. These values were used as the µ, 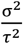, and τ^2^ parameters in the SVG simulation model. For the single-kernel bandwidth parameter *b*, among SVGs identified by nnSVG, we used the 25th, 50th, and 75th percentiles of the *b* estimates obtained by nnSVG, and generated separate datasets for each *b* value, which we refer to as ‘1 kernel I’, ‘1 kernel II’, and ‘1 kernel III’, respectively. Forty SVGs were generated for each parameter combination, yielding a total of 1,000 SVGs per dataset. For the five-kernel covariance, three sets of five bandwidth parameter values were used: 10th, 20th, 30th, 40th, and 50th percentiles; 10th, 30th, 50th, 70th, and 90th percentiles; and 50th, 60th, 70th, 80th, and 90th percentiles of *b* estimates obtained by nnSVG among identified SVGs. These corresponded to the weakest, intermediate, and strongest spatial autocorrelations, respectively, and the resulting datasets were named ‘5 kernels I’, ‘5 kernels II’, and ‘5 kernels III’. Forty SVGs were generated per parameter combination, yielding 1,000 SVGs per dataset. For non-SVGs, the parameter µ value was set to the mean of nnSVG estimates for all genes, and τ^2^ values were set to the 10th, 30th, 50th, 70th, and 90th percentiles of nnSVG estimates among all genes. For each parameter combination, 200 non-SVGs were generated, yielding 1,000 non-SVGs per dataset. Thus, each simulation dataset comprised 2,000 gene expressions.

We performed custom-designed simulations representing canonical spatial patterns, including gradients, streaks, and hotspots, based on Weber et al.^8^ and Shang et al.^13^. As in the covariance-based simulations, spatial coordinates and parameter settings were derived from the 10x Visium Human DLPFC dataset^25^. For non-SVG simulations, we applied the covariance-based design, using the mean of normalized and log-transformed expressions as the parameter µ value, and the 25th, 50th, and 75th percentiles of the variance as τ^2^. For each τ^2^ value, 360 non-SVGs were generated, yielding a total of 1,080 non-SVGs per dataset. For SVG simulations, we considered two tissue regions—the white matter and cortical layers, and generated gradient, streak, or hotspot patterns with different bandwidths in each region. For each canonical pattern, we obtained nine distinct spatial patterns having different bandwidths in two regions. After simulating non-SVGs as described above, we modulated expression values along the spatial patterns by downregulating them by 0.25x or 0.5x, or upregulating them by 2x or 4x, thereby generating gene expression data with spatial patterns (see **Supplementary Figure S2**). This procedure produced 1,080 SVGs for each pattern, yielding three datasets with a total of 2,160 simulated gene expression profiles. Additionally, to evaluate method performance under increased noise, we generated versions of each dataset in which 25%, 50%, or 75% of spots were randomly shuffled.

To assess the robustness of Spacelink to spot dropout, data sparsity, and reduced read depth, we performed (i) spot downsampling at varying probabilities as in Chen et al.^36^, (ii) data sparsification, in which gene expression values were set to zero at selected spatial locations with varying proportions, and (iii) binomial thinning of library size, applied with different probabilities of read-depth reduction following the strategy in Dey et al.^37^. For all three analyses, we used scDesign3-based simulation data with the 10x Visium Human DLPFC dataset as a reference. In the downsampling experiments, 10%, 20%, …, or 90% of spots were randomly removed across the entire dataset, generating datasets with fewer spatial locations. In the data sparsification experiments, for each gene, 10%, 20%, …, or 90% of spots were randomly selected and their expression values set to zero, producing datasets with varying levels of sparsity. In the binomial thinning experiments, each gene expression count was resampled from a binomial distribution with probabilities of 0.1, 0.2, …, or 0.9.

The simulation dataset used to compare length scale estimation between nnSVG^8^ and Spacelink was generated using the covariance-based simulation design with a single-kernel covariance. Because both nnSVG and Spacelink operate on normalized and transformed data, we did not use the count data **z**, but applied the models directly to **y**. The bandwidth parameter *b* was set to the 25th, 50th, and 75th percentiles of *b* estimates among SVGs identified by nnSVG. For each setting, 1,250 SVGs were generated, yielding a total of 3,750 SVGs for the analysis.

To evaluate ESV under varying numbers of spots, sparsity levels, and technical noise across tissues with different structural complexity, we generated scDesign3-based simulated datasets using 10x Visium human kidney, prostate cancer, and DLPFC data as references. For each tissue, we simulated 50 genes with high spatial variability and constructed datasets under three settings: (1) varying numbers of spots by randomly selecting 50% to 95% of spots in 5% increments (total: 50 × 10 = 500 genes); (2) varying sparsity by setting 5% to 95% of spot expression values to zero (total: 50 × 20 = 1000 genes); and (3) varying noise levels by taking weighted averages of the original expression and random noise (*α*=0.05 to 0.95; higher *α* indicates higher noise) (total: 50 × 20 = 1000 genes).

For simulations under the null hypothesis, we generated 10 datasets by permuting spatial spots from real data (10 human 10x Visium samples). We also included an additional 10 datasets consisting of simulated non-spatially variable genes (non-SVGs) generated by scDesign3, as in **Figure 2a**. For high-sparsity null simulations, we generated data following the procedure described in Zhu et al. (2021)^48^. Using the Visium human DLPFC dataset, we first generated 10,000 spatial spots within the tissue region via a Poisson point process. Gene expression was then simulated from a negative binomial distribution with a mean of 0.005 and a dispersion parameter of 2.5. In total, 10,000 genes were generated.

For the comparison of computational efficiency, global SVG detection methods were evaluated using the covariance-based simulation design with a single-kernel covariance. Instead of the spatial coordinates from the 10x Visium Human DLPFC dataset^25^, rectangular artificial tissues of various sizes were generated with dimensions 25×20, 25×40, 50×100, 100×100, 200×250 and 500×600. Due to computational burden, for simulation with 200×250 and 500×600 spots, we used 1,000 nearest neighbors to construct kernel covariance matrix and set the covariance of distant spots to 0. In each setting, 300 gene expressions were simulated.

To assess robustness to unbalanced data, we used scDesign3-based simulation datasets with the 10x Visium Human DLPFC data^25^ as a reference. In one scenario, 1,000 simulated SVGs were retained, and among non-SVGs, 5%, 10%, …, or 95% were randomly retained, producing datasets with more SVGs than non-SVGs. Conversely, 1,000 simulated non-SVGs were retained, and among SVGs, 5%, 10%, …, or 95% were randomly retained, yielding datasets with more non-SVGs than SVGs.

### Simulation design for benchmarking cell type SVG methods

To evaluate ct-SVG detection methods, we generated simulated datasets following the framework of Shang et al.^13^, using 10x Visium human DLPFC^25^ and STARmap mouse primary visual cortex^46^ datasets as references. We first obtained single-cell resolution spatial transcriptomics data and aggregated single-cell measurements into spot-level measurements for the analyses.

For the single-cell resolution simulations, we generated 20,000 cells on each tissue using a random-point-pattern Poisson process. Each cell was assigned to one of four spatial domains and one of four cell types, under four scenarios of cell type composition:

Scenario 1: Each domain contains only one cell type.

Scenario 2: Each domain contains 90% of one cell type and 5% of two others.

Scenario 3: Each domain contains 50% of one cell type and 25% of two others.

Scenario 4: Each domain contains three cell types at equal proportions.

Cell type distributions for each scenario are summarized in **Supplementary Table S2**. Based on the original datasets, we simulated gene expression for each cell. A negative binomial model was first fitted to real gene expression count data, and mean and dispersion estimates were extracted. We use the average of mean estimates and the median of the dispersion estimates as the parameter values for the simulation model. Using these parameters, we generated random negative binomial values to simulate non-SVGs (384 per scenario).

We next simulated cell type marker genes, defined as genes up- or downregulated only in a specific cell type but without spatial variation. We first generated non-SVGs, and multiplied the expression of cells belonging to the target type by 4, 2, 0.5, or 0.25. For each scenario, 96 marker genes were generated per cell type, totaling 384.

To simulate ct-SVGs, we adapted the pattern-based design used for global SVGs, based on a hotspot pattern. For each scenario, we randomly selected one cell of a target cell type as the hotspot center, then defined the hotspot radius using the 25th, 50th, or 75th percentile of distances to other cells of the same type. Within each hotspot, cells were divided into upper and lower halves. After generating either a non-SVG or a marker gene with one of four up- or down-regulation parameter values (4, 2, 0.5, or 0.25), we upscaled expression in the upper half by 2.5 or 2, and in the lower half by 2, or downscaled expression in the upper half by 0.5 or 0.4, and in the lower half by 0.5. This procedure produced ct-SVGs or marker ct-SVGs with hotspot spatial patterns. In each of the 3×4 and 3×4×4 scenarios, 16 ct-SVGs and 4 marker ct-SVGs were generated per cell type, yielding 768 of each in total. As a result, by combining the 384 non-SVGs with 384 non-ct-SVG cell type marker genes, we obtained 768 non-ct-SVGs in total, and for each cell type, 768 ct-SVGs and 768 marker ct-SVGs, thereby allowing for a balanced evaluation of sensitivity and specificity.

Finally, to simulate spot-level data, we overlaid a square grid on the single-cell resolution simulation. Each grid cell was treated as a spot, with gene counts summed across all cells inside. From 20,000 cells, we obtained 3,056 spots, each containing between 1 and 19 cells (mean 6–7). We recorded the true cell type proportion of each spot and additionally estimated proportions using RCTD, with the single-cell data as reference. Both the true and inferred proportions were used for ct-SVG methods in evaluation.

For the computational efficiency comparison of ct-SVG detection methods, scenario 3 of the pattern-based simulation design was used with the 10x Visium Human DLPFC dataset as a reference for parameters. Rectangular artificial tissues with dimensions 25×20, 25×40, 50×50, 50×100, and 65×100 were used instead of the original coordinates. In each setting, 300 gene expressions were simulated.

### Performance metrics to benchmark SVG detection methods

Throughout the paper, we evaluated SVG detection methods using various metrics, including false discovery rate (FDR), statistical power, area under the precision-recall curve (AUPRC), and Kendall’s tau correlation. FDR and power were computed from p-values with a significance cutoff of 0.05. AUPRC was used to assess ranking performance based on the binary ground truth label (SVG vs. non-SVG), implemented with the `pr.curve` function from the PRROC package. When a ground truth ranking of spatial variability was available, we used Kendall’s tau correlation for assessment of SVG ranking methods, which is defined as:

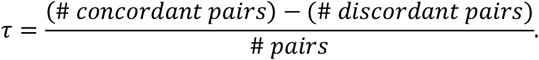

To assess accuracy in length scale estimation, for nnSVG, spNNGP and spBayes, we computed 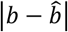 (true vs. estimated length scale), while for Spacelink we used the following absolute distance: 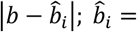 the bandwidth corresponding to the largest estimated sigma among five candidates.

For domain detection methods, we used the adjusted Rand index (ARI) and normalized mutual information (NMI), defined as:

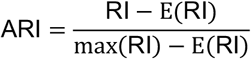

where RI is the Rand index measuring the fraction of agreement between two partitions, and

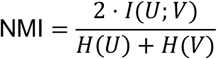

where *I*(*U*; *V*) is the mutual information between partitions *U* and *V*, and *H*(·) denotes entropy.

For the evaluation under the null simulation, we used the Kolmogorov–Smirnov (K–S) test. The K–S test assesses the maximum deviation between the empirical distribution function and the reference distribution, providing a general measure of goodness-of-fit. We implemented it using the `ks.test` function from the `stats` package.

### Processing spatial transcriptomic datasets and metadata

For the analysis of spatial transcriptomic datasets, we used sctransform as the default normalization method. To ensure fairness, all other SVG detection methods were also applied to counts or log-counts normalized with sctransform^73^. Because the focus of our study is not normalization, we did not conduct a systematic comparison of different normalization approaches. It is worth noting that the appropriate strategy for normalizing spatial transcriptomics data—indeed, whether normalization should be performed at all—remains controversial. Prior work has reported that sctransform can sometimes overcorrect for both library size and biological effects, leading to reduced performance in tasks such as spatial domain identification^125,126^. However, in our benchmarking against scater^127^, we found that certain technical biases were insufficiently removed by this alternative, resulting in false discoveries in SVG detection. For this reason, we adopted sctransform as the default. Our Spacelink implementation allows users to input data normalized by whichever method they deem most appropriate for their own datasets.

In the global SVG simulation analysis, we did not apply library size normalization, as technical biases across spots are absent in simulations. By contrast, in the ct-SVG simulations using spot-resolution data, we applied sctransform because rasterizing cell-resolution data into spots results in a varying number of cells per spot, which introduces variation in library size that requires adjustment.

For CosMx datasets and the mouse organogenesis Stereo-seq spatiotemporal dataset^27^, we applied rasterization for computational feasibility using the `SEraster::rasterizeGeneExpression` function with settings `fun = “sum”` and `square = FALSE`^54^. In the CosMx data, we used a resolution of 100 µm, and in the “Bin50” resolution mouse dataset, we used a resolution of 4 for the whole embryo and non-brain regions (rasterization was not applied to brain regions given the smaller number of spots). In the mouse dataset, rasterization was applied without distinguishing between tissues within the whole embryo or non-brain regions, so a single rasterized spot could contain cells from multiple tissues. For analyses comparing CosMx and Visium datasets, we adopted a slightly modified rasterization to better harmonize the two platforms: we used `SEraster::rasterizeGeneExpression` to obtain spots with neighbors’ distances fixed at 100 µm, then defined a 55 µm-diameter circle around each spot and summed expression values within this circle. This produced rasterized data minimizing discrepancies with Visium.

### Other SVG methods benchmarked against Spacelink

We employed six other methods for global spatially variable gene (SVG) identification and quantification to benchmark the performance against Spacelink. Moran’s I^32^ is a standard measure of spatial autocorrelation, where the statistic ranges from –1 to 1; values closer to 1 indicate stronger positive spatial autocorrelation. To compute Moran’s I, we used the `spatial_neighbors` function from the `squidpy`^128^ Python package with parameters `coord_type=“generic”, delaunay=True` to define spatial neighbors, followed by the `spatial_autocorr` function with `n_perms=100` to obtain both the Moran’s I statistic and the FDR-corrected one-sided p-value `(pval_z_sim_fdr_bh)`. SpatialDE^9^ is an early statistical framework for SVG detection that uses Gaussian process regression to model spatial patterns. Using the `SpatialDE` Python package, we ran `SpatialDE.run` to obtain the fraction of spatial variance (FSV), the log-likelihood ratio (`LLR`), and the corrected p-value (`qval`). SpatialDE2^10^ extends the model by combining multiple spatial kernels with a Poisson distribution, providing a framework for spatial count data. It introduces an omnibus test to jointly evaluate multiple kernels under a generalized linear model. We obtained the source code from https://github.com/PMBio/SpatialDE and applied the `SpatialDE.test` function with omnibus=True to calculate the adjusted p-value (`padj`). nnSVG^8^ leverages the nearest neighbor graph to evaluate each gene’s expression pattern across the spatial neighborhood structure, estimating spatial autocorrelation to identify SVGs. Using the `nnSVG` R package, we ran the `nnSVG` function with parameters `order=“AMMD”`, `n_neighbors=10` to obtain the proportion of spatial variance (`prop_sv`), the log-likelihood ratio (`LR_stat`), and the adjusted p-value (`padj`). SPARK^11^ models count data using a generalized linear spatial model (GLSM) based on the Poisson distribution, testing multiple kernels separately and combining the resulting p-values for SVG detection. SPARK-X^48^ adopts a nonparametric covariance test that evaluates multiple kernels with high computational efficiency. We applied the `SPARK` R package’s `spark.test` and `sparkx` functions to obtain the adjusted p-values (`adjusted_pvalue` and `adjustedPval`) of SPARK and SPARK-X models, respectively.

We additionally considered two Bayesian spatial models, spNNGP^44^ and spBayes^45^. Both assume a single spatial scale per gene; spNNGP is based on the nearest-neighbor Gaussian process (NNGP) approximation, whereas spBayes uses a Gaussian predictive process. We applied `spNNGP` function from the `spNNGP` R package and `spLM` function from the `spBayes` R package to fit the models. For both models, the spatial decay parameter `phi` was initialized as 3/(0.5 × maximum distance between spots) following the choice made in the primary simulation examples used in the spNNGP and spBayes papers^44,45^ and reference manuals^129,130^ (we did not find a clear default option in the corresponding package). In spNNGP, a uniform prior was specified over 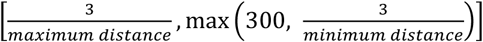, while in spBayes the range was 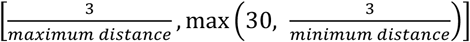, this again follows choices made in the primary simulation examples in the respective spNNGP and spBayes reference manuals^129,130^ (we did not find a clear default option in the corresponding packages). The scaling by distance between spots is aimed at aligning the scale of the initial value and prior of `phi` to match the unit domain size used in these simulation examples. All other parameters were set to the default values from these simulation examples. We drew 5,000 posterior samples for each model and used the final 500 samples to compute posterior means as parameter estimates. This choice was motivated by observed instability in posterior samples with 2,000–3,000 iterations, while larger sample sizes were avoided due to computational cost. Hypothesis testing to assess *H*_+_: σ^g^ = 0 against *H*_1_: σ^g^ ≠ 0 was performed using a log-likelihood ratio test based on the estimated parameters, following the same procedure as in nnSVG.

To benchmark performance against the Spacelink cell-type-specific model, we applied two methods— Celina^13^ and STANCE^12^—for identifying cell-type-specific spatially variable genes (ct-SVGs). Both rely on Gaussian process models but differ in design: Celina tests multiple kernels and combines the resulting p-values using a Cauchy combination method, while STANCE uses a single kernel. Celina also does not account for the spatial effects of other cell types, whereas STANCE incorporates the spatial effects of all other cell types into the model. For implementation, we followed the official tutorials for each method (Celina: https://lulushang.org/Celina_Tutorial/index.html; STANCE: https://haroldsu.github.io/STANCE/tutorial.html).In Celina, we used the `Testing_interaction_all` function from the `CELINA` R package to compute the combined p-values (`CombinedPvals`) and determined significance based on an empirical null distribution generated from 10 permutations of spot coordinates, with a 0.01 empirical FDR cutoff, as described in the original paper. In STANCE, we first applied the `runTest1` function from the `STANCE` R package to perform the overall score test. Genes with adjusted p-values (`p_value_adj`) < 0.05 were then subjected to `runTest2` to obtain p-values, and significance was assessed at the 0.05 level.

We additionally applied two methods to derive gene programs for benchmarking the disease informativeness of the Spacelink gene program. The sc-linker cell type program^51^ was generated from scRNA-seq data by performing differential expression analysis for each cell type versus all others using the Wilcoxon rank-sum test, with p-values transformed into scores between 0 and 1. The gsMap gene specificity score (GSS) program^7^ was computed by forming micro-domains of homogeneous neighboring spots, averaging gene expression ranks across each micro-domain, and normalizing by the global average rank to yield spot-level specificity scores. All scores were normalized to probabilistic scale ranging from 0 to 1 to facilitate fair comparisons.

### Processing disease gene-level association statistics

To analyze the association of complex traits and diseases with spatial variability, we considered three disease-relevant gene sets: (i) MAGMA-prioritized genes, (ii) PoPS-scored genes, and (iii) Mendelian disorder genes. MAGMA identifies genes with strong GWAS associations by aggregating SNP-level effects into gene-level scores^30^, whereas PoPS trains a linear model on the MAGMA scores using gene-level features derived from single-cell expression data, gene pathways and protein–protein interactions^29^. We used the published PoPS scores as generated by Weeks et al.^29^ for 78 UKBiobank traits, and additionally computed the PoPS scores for additional 35 non-UKBiobank traits to facilitate the SVG comparative analysis across tissues (**Supplementary Table S1**). Mendelian disorder genes for 22 Mendelian disorders, including monogenic autism were curated from OMIM^84^ by Freund et al.^85^. For MAGMA, we defined disease-relevant sets by selecting genes with Z-scores greater than 3 or by choosing the top N genes with the highest signed Z-scores, using varying values of N. For PoPS, we defined disease-relevant sets by selecting the top N genes with the highest PoPS scores.

### Evaluating complex disease informativeness of Spacelink and other SVG programs

To assess the disease informativeness of different gene programs, including Spacelink-based SVG programs, while controlling for potential confounders such as mean expression, we fit logistic regression models of the form:

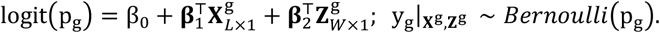

Here, y_g_ is a binary variable indicating whether gene g belongs to the top 500 PoPS-scored^29^ (or MAGMA-prioritized^30^) genes for each trait; β_0_ is the intercept; **X**^g^ denotes *L* predictive features of interest representing gene programs (including Spacelink spatial SVG programs and other programs of interest for benchmarking) with coefficients **β**_1_; and **Z**^g^ contains covariates representing confounders with coefficients **β**_2_ . As confounders, we included the mean and variance of normalized, log-transformed expression values—computed across the whole tissue when comparing different gene programs (**Figures 4a–d**), and within a specific cell type when examining disease-specific enrichment of Spacelink ct-ESV programs (**Figure 4e**)—as well as the proportion of non-zero expression values and a binary indicator for high variability (defined by `scran::modelGeneVar`^131^ with FDR < 0.05).

We performed two types of analyses.

1. *Single-program models* Here, **X**^g^ contained a single gene program (*L* = 1) as a predictive feature. Programs included Spacelink ESV or ct-ESV scores, sc-linker programs per cell type^51^, gsMap gene specificity scores (GSS)^7^ per cell, nnSVG PropSV or ct-PropSV scores, SpatialDE FSV or ct-FSV scores, or binary indicators for global or cell-type-specific SVGs defined by SVG detection methods. All these gene programs are non-negative by design and to ensure fair comparison, each program was rescaled to the [0,1] interval as follows.

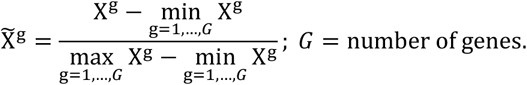

For cell-type-specific programs, SVG detection or scoring methods (including Spacelink) were applied at the cell-type level by subsetting expression in the CosMx data to the corresponding cells. Performance was evaluated using probabilistic recall and adjusted odds ratio, defined as follows:

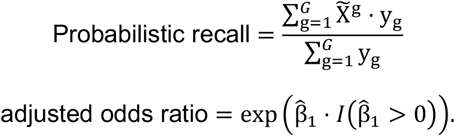
2. *Multi-program models* Here, **X**^g^ contains multiple programs (*L* > 1) as predictive features in the same model. For example, when evaluating Spacelink scoring method, we constructed an **X** matrix with the global ESV and all ct-ESV scores (average r=0.57 across datasets); when evaluating sc-linker, we included programs across all cell types (average r=0.18 across datasets). For GSS, due to the large number of programs (equal to the number of spots) and strong correlations among them, we retained only programs with pairwise correlations < 0.5. Genes were partitioned into training and test sets by randomly splitting the 22 autosomes into two groups of 11; genes on one group of chromosomes were assigned to training and the other to testing. We opted for this random split against leave-one-chromosome-out approach as in Weeks et al.^29^ and Fabiha et al.^53^, as the number of positives per chromosome is very small. Logistic regression models in this setting were fitted on the training set with a ridge penalty to account for collinearity across features, and performance was assessed on the test set. The evaluation metric was relative AUPRC, defined as the ratio between the model’s AUPRC on the test set and the baseline AUPRC (equal to the proportion of the top 500 PoPS-scored (or MAGMA-prioritized) genes in the test set).

### Temporal regression of Spacelink ESV in mouse organogenesis spatiotemporal data

To investigate the temporal dynamics of gene-level spatial variability while accounting for systematic biases, we employed a two-step regression framework with stage-specific (or sample-specific) random effects. In the first step, we estimated the conditional mean of stage random effect by fitting the following regression model:

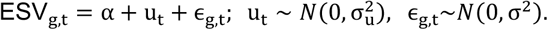

Here, ESV_g,t_ indicates the raw ESV value for gene g at the t-th developmental stage (t = 1, …,8), α is the intercept, u_t_ represents the random effect at stage t, and ϵ_g,t_ is the noise component. We estimated the conditional mean of random effect, û_t_, from this model and subtracted it from each gene’s ESV at each stage:

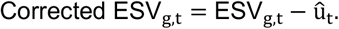

In the second step, we fitted a gene-wise linear regression model using corrected ESV:

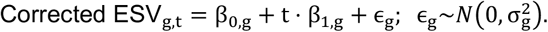

The same correction and regression procedure was applied to the mean and variance of normalized, log-transformed expression values. Genes were defined as showing an association between spatial variability and developmental progression—but without expression changes—if the slope in the corrected ESV regression was significant (P < 0.05) while the slopes in regressions on corrected mean and variance were not (P > 0.2).

To identify genes with divergent patterns between brain and non-brain regions, we considered cases where (i) one region exhibited an increasing trend in corrected ESV 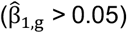 while the other showed any of the following: a decreasing trend in corrected ESV 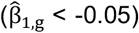, a monotonically high profile 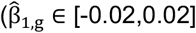 with corrected ESV > 0.4 in at least 4 stages), or a monotonically low profile 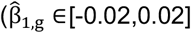 with corrected ESV < 0.2 in at least 4 stages); or (ii) one region exhibited a decreasing trend while the other showed any of: an increasing trend, a monotonically high profile, or a monotonically low profile, as defined above.

### Assessment of model assumptions

In the second step of the two-step model (the gene-specific regression), over 99% of cases satisfied the normality assumption and 100% met homoscedasticity, with moderate coefficients of determination on average (**Supplementary Figure S13e**). Here, normality and homoscedasticity were assessed using Shapiro–Wilk and Breusch–Pagan tests, respectively (BH-adjusted p-value threshold: 0.05). However, the first step — a linear mixed model for the stage random effect — failed both assumption tests and showed low R^2^ (Kolmogorov–Smirnov p < 2e-16; Breusch– Pagan p < 2e-16; R^2^ = 0.02–0.03 across whole-tissue, brain, and non-brain models). We considered two alternatives. Applying a logit transformation to ESV improved fit for some models but still violated most assumptions for the brain and non-brain models (Kolmogorov–Smirnov p < 2e-16 for all models; Breusch–Pagan p < 2e-16 except for the whole-tissue model where p = 0.8; R^2^ = 0.02–0.08). Binarizing ESV at multiple cutoffs and fitting logistic regression models — using Firth’s bias-reduced logistic regression to address small-sample bias, with stage-level random effects included as an offset — captured trends in some individual genes but oversimplified the continuous nature of ESV and failed to retain apparent temporal trends in all cases. For example, gene *Camp* in **Supplementary Figure S13f** shows an increasing trend in spatial variability, which is captured by our original two-step model; however, the logistic regression model does not detect this trend as significant under any binarization cutoff. In the absence of a clearly superior alternative, we verified robustness by applying logit-transformed ESV in all downstream analyses. The main conclusions remained the same (**Supplementary Note**). Importantly, our downstream analyses use only the sign of the ESV trend — positive, negative, or absent — rather than coefficient magnitudes or p-values. We therefore consider our two-step approach appropriate in practice: despite misspecification in the first step, it reliably identifies the direction of ESV trends, which is the only quantity used in downstream analyses. **Infeasibility of single linear mixed model**. One might consider a single linear mixed model accounting for stage-specific random effects:

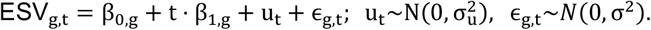

However, this approach is infeasible, as it requires simultaneous estimation across all genes, resulting in a prohibitively large number of parameters. Fitting a linear mixed model per gene is also inadequate, as each gene has only one ESV value per stage, making stage-specific random effects ill-defined.

### Differential ESV analysis in mouse organogenesis spatiotemporal data

We conducted differential gene expression (DGE) analysis with 9,481 and 16,566 genes that were identified as SVGs in brain and non-brain regions, respectively, in at least 6 of the 8 developmental stages. For each non-brain tissue, at each developmental stage, we fitted a zero-inflated negative binomial (ZINB) model to test whether genes were expressed at different levels inside the tissue compared to the other non-brain regions. Genes were defined as differentially expressed if they showed a fold change greater than 1 and an adjusted p-value below 0.05 in at least three stages for that tissue.

### Associating Spacelink against Alzheimer’s disease pathology markers

To assess changes in average spatial variability across pathological progression while accounting for donor effects, we fitted the following mixed model:

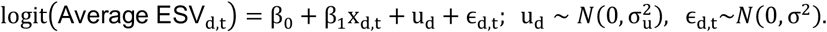

Here, Average ESV_d,t_ denotes the mean ESV value across all genes in the t-th sample from donor d,

*β*_0_ and *β*_1_ are fixed effects, x_d,t_ is the pathological measurement for the t-th sample of donor d, u_d_ is the random intercept for donor d, and ϵ_d,t_ is the residual error term. Using the conditional mean of donor-level random effect û_d_, we obtained the donor-corrected average ESV as logit^−1^(logit(Average ESV_d,t_) − û_d_) . The above model met the normality assumption but slightly violated homoscedasticity (Shapiro–Wilk p = 0.49-0.76; Breusch–Pagan p = 0.02-0.03; R^2^ = 0.68-0.69).

To systematically identify genes whose spatial variability is associated with Alzheimer’s disease (AD) pathology, we performed regression analyses linking gene-level ESVs across samples to pathological measures, while adjusting for cortical layer composition and mean expression. Specifically, we fit the following model for each gene:

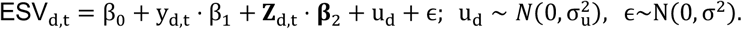

where ESV_d,t_ refers to the ESV of the gene in the t-th sample of donor d; y_d,t_ denotes the square root– transformed amyloid burden, tangles, or number of plaques for the sample; **Z**_d,t_ indicates covariates representing confounders including the mean of normalized and log-transformed expression of the gene and the proportions of spots corresponding to each cortical layer within the sample; u_d_ is the random intercept for donor d. Genes whose regression coefficient for y_d,t_ had a p-value < 0.05 were considered to have spatial variability significantly associated with AD pathology. In the above gene-specific model, over 98% of genes satisfied the normality assumption, more than 90% met homoscedasticity, and coefficients of determination were high on average (**Supplementary Figure S17i**).

To identify genes whose mean expression is associated with AD pathology, we applied the same model but replaced ESV_g_ with the mean normalized log-expression of gene g as the predictor, while including cortical layer proportions as covariates.

### Excess of overlap (EOO) analysis

To quantify the concordance between two gene sets, we used an excess of overlap (EOO) statistic^132^. The EOO is defined as the ratio of the observed overlap to the expected overlap under the null hypothesis of random overlap:

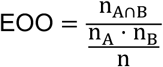

where n_A∩B_ is the number of genes shared between the two sets, n_A_ and n_B_ are the sizes of the respective gene sets, and n is the total number of genes considered. We assessed the statistical significance of the excess overlap using Fisher’s exact test.

### Colocalization analysis

We leveraged 17 cis-xQTL contexts spanning 3 molecular modalities (gene expression, splicing, protein abundance) from the aging brain cortex of ROSMAP donors (average N = 595)^105^. All participants enrolled without known dementia and agreed to detailed clinical evaluation and brain donation at death. All studies were approved by an Institutional Review Board of Rush University Medical Center. Each participant signed informed and repository consents and all ROSMAP participants signed an Anatomic Gift Act. The eQTL data encompassed bulk RNA-seq from three cortical regions (dorsolateral prefrontal cortex, posterior cingulate cortex, and head of caudate nucleus), bulk monocytes isolated from peripheral blood, and single-nucleus RNA-seq pseudo-bulk data from six major brain cell types (excitatory neurons, inhibitory neurons, microglia, oligodendrocyte precursor cells, oligodendrocytes, and astrocytes). Bulk-sQTL were generated from three cortical regions using LeafCutter2^133^ and bulk-pQTL was performed on the dorsolateral prefrontal cortex. Standard quality control procedures and normalization were applied to all molecular phenotypes, with technical factors including batch effects, RNA integrity, and post-mortem interval adjusted alongside biological covariates such as sex, age at death, and principal components, as described in our previous study^105^.

To understand molecular mechanisms underlying spatially variable genes associated with AD and AD pathology, we conducted two targeted multi-trait colocalization analyses using ColocBoost^105^: pathology-xQTL and AD-xQTL colocalizations. For the pathology-xQTL colocalization analysis, we generated pathology phenotypes based on the square-root transformed amyloid burden and tau accumulation metadata corresponding to 595 ROSMAP samples, and then performed the same adjustment by covariates as in cis-xQTL data, except for expression principal components, as the latter can be causally mediated by the underlying pathology. We colocalized these phenotypes against 17 cis-xQTLs in the ColocBoost multi-phenotype colocalization model. For the AD-xQTL colocalization analysis focusing on *CLU*, we utilized summary statistics from a recent AD GWAS meta-analysis^134^ (N_case=111,326, N_control=677,663) after performing quality control against a custom LD reference panel derived from European ancestry samples in the Alzheimer’s Disease Sequencing Project (ADSP)^105^.

## Data Availability

The 10x Visium datasets for DLPFC, prostate, lymph node, heart, kidney, lung cancer, cervical cancer, breast cancer, prostate cancer, and skin melanoma used as input for scDesign3 are available at the following links: https://research.libd.org/spatialLIBD/ (sample ID: 151673), https://www.10xgenomics.com/datasets/normal-human-prostate-ffpe-1-standard-1-3-0, https://www.10xgenomics.com/datasets/human-lymph-node-1-standard-1-0-0, https://www.10xgenomics.com/datasets/human-heart-1-standard-1-0-0, https://www.10xgenomics.com/datasets/human-kidney-11-mm-capture-area-ffpe-2-standard, https://www.10xgenomics.com/datasets/human-lung-cancer-11-mm-capture-area-ffpe-2-standard, https://www.10xgenomics.com/datasets/human-cervical-cancer-1-standard, https://www.10xgenomics.com/datasets/human-breast-cancer-ductal-carcinoma-in-situ-invasive-carcinoma-ffpe-1-standard-1-3-0, https://www.10xgenomics.com/datasets/human-prostate-cancer-adjacent-normal-section-with-if-staining-ffpe-1-standard, https://www.10xgenomics.com/datasets/human-melanoma-if-stained-ffpe-2-standard, respectively.

The STARmap data on mouse visual cortex is available at https://xzhoulab.github.io/SRTsim/.

The CosMx datasets for brain cortex, lymph node and liver are available at the following links: https://nanostring.com/products/cosmx-spatial-molecular-imager/ffpe-dataset/human-frontal-cortex-ffpe-dataset/, https://nanostring.com/products/cosmx-spatial-molecular-imager/ffpe-dataset/cosmx-human-lymph-node-ffpe-dataset/, https://nanostring.com/products/cosmx-spatial-molecular-imager/ffpe-dataset/human-liver-rna-ffpe-dataset/

The 10x Visium dataset for lymph node used for cross-platform evaluation is available at https://www.10xgenomics.com/datasets/human-lymph-node-1-standard-1-1-0.

The Stereo-seq mouse organogenesis spatiotemporal transcriptomics data is available at https://db.cngb.org/stomics/mosta/.

The mouse in-vivo Perturb-seq data is available through the Gene Expression Omnibus (accession GSE157977).

The 32 ROSMAP 10x Visium samples from brain dorsolateral prefrontal cortex (DLPFC) at different stages of Alzheimer’s disease (AD) are available via the AD Knowledge Portal (https://adknowledgeportal.org). Pathologic and phenotypic data from ROSMAP are available at https://www.radc.rush.edu.

The 10x Visium data from 5xFAD transgenic mouse model is available through the Gene Expression Omnibus (accession GSE233208).

The supplementary tables contain the Spacelink hypothesis test results, ESV and ct-ESV scores across different datasets, as well as the data needed to reproduce the main and supplementary figures.

## Code Availability

Spacelink is available as an R package, with the source code freely accessible at https://github.com/hanbyul-lee/spacelink. All the codes to reproduce the Figures in the manuscript are provided separately in the GitHub repository: https://github.com/hanbyul-lee/spacelink_paper.

## Acknowledgements

K.K.D. was supported by R00HG012203 and R01HG014008, NIH/NCI Cancer Center Support Grant P30CA008748, AWS Imagine Fund and Josie Robertson Investigator Program. HUK and MT were supported by an Alzheimer’s Association Grant through the AD Strategic Fund (ADSF-21-816675). ROSMAP is supported by P30AG10161, P30AG72975, R01AG17917, R01AG015819, U01AG072572, and U01AG046152.

## Supplementary Figures

**Supplementary Figure S1. Schematics for selecting data-adaptive length scales and computing distances in Spacelink-lite, and illustration of ESV, PropSV and FSV across different signal-to-noise ratios and length scales**

(a) We first initialize 2*L* (default *L* = 5) log-spaced bandwidths between the minimum and maximum spatial distances across *N* locations, which are shared across all genes. For each gene g, we fit the Spacelink global model using Non-Negative Least Squares (NNLS) and identify the smallest and largest bandwidths with non-zero weights (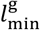 and 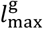 ). We then define *L* log-spaced bandwidths within 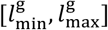] to obtain the final gene-specific set of spatial bandwidths and corresponding kernels.(b)In Spacelink-lite, distances are computed differently depending on the relative locations of spots. The tissue space is first divided into grids. If two spots lie within neighboring grids (each grid has up to eight surrounding neighboring grids), their distance is calculated exactly. Otherwise, the distance is approximated using the distance between the corresponding grid centers. (c). Contour heatmaps of three spatial variability metrics — ESV, PropSV, and FSV — across a parameter space of signal-to-noise ratio and length scale in a single-kernel model, showing how each metric varies across this space.

**Supplementary Figure S2. Schematic of scDesign3-based and pattern-based simulations.**

(a). **scDesign3-based simulation**. We first selected the top 50 SVGs using Moran’s I and model deviance explained (via scDesign3 functions). For each gene, we derived a mean expression parameter vector capturing spatial correlation (*µ*_*s*_) and generated a non-spatial counterpart (*µ*_*ns*_) by randomly shuffling *µ*_*s*_. Data were simulated with varying spatial variability using weighted mixtures *α* ⋅*µ*_*s*_ + (1 − *α*) ⋅ *µ*_*ns*_, with *α* ∈ {0.05, 0.1, …, 1}. Smaller *α* values indicate weaker spatial variability, while larger values indicate stronger spatial variability. (b). **Pattern-based canonical simulation**. We used two tissue regions from 10x Visium Human DLPFC data—the white matter and cortical layers—and generated hotspot, streak, or gradient with different bandwidths. Each canonical pattern (hotspot, streak, gradient) produced nine distinct spatial configurations (three bandwidths × two regions; top panel). By up- or downregulating non-SVG expression along these spatial patterns, we generated gene expression data with spatial structure (bottom panel, left). To assess robustness, we additionally generated versions of each dataset in which 25%, 50%, or 75% of spots were randomly shuffled (bottom panel, right).

**Supplementary Figure S3. Additional results from global SVG simulation analyses.**

(a). Statistical power and false discovery rate (FDR) of Spacelink versus six other global SVG identification methods across 6 covariance-based and 12 pattern-based simulation settings. (b). Area under the precision–recall curve (AUPRC) of Spacelink p-values compared with p-values from the six global SVG hypothesis tests across 6 covariance-based and 12 pattern-based simulations. Asterisks indicate significance from 1-sided pairwise t-tests comparing Spacelink against each method (*** P < 0.001). (c). Statistical power, FDR, AUPRC, and Kendall’s tau correlation of Spacelink versus global SVG methods under varying levels of spot downsampling, data sparsification, and library size thinning (**Methods**). (d). An alternative visual illustration of **Figure 2d**. Left: Violin plots of distances between true and estimated length scales from Spacelink and nnSVG in a set of 3,750 simulated SVGs aligned with nnSVG’s framework. Right: An example simulation where Spacelink closely recovers the true length scale whereas nnSVG does not. (e). Extended violin plot of distances between true and estimated length scales from nnSVG, spNNGP, spBayes, and Spacelink. Asterisks indicate significance from 1-sided pairwise t-tests (*** P < 0.001). (f). FDR and statistical power comparisons across methods on 10 scDesign3-based simulation datasets. (g). Kendall’s tau correlation comparing SVG ranking consistency across methods on 10 scDesign3-based simulation datasets. Asterisks indicate significance from 1-sided pairwise t-tests comparing Spacelink against each method (*** P < 0.001, * P < 0.05). For pairs showing obvious differences by visual inspection, t-test results are omitted. (h). Boxplots of estimated signal-to-noise ratio (SNR) as a function of the spatial variability parameter *α* for Spacelink, nnSVG, spNNGP, and spBayes. (i). Runtime comparison for Spacelink, nnSVG, spNNGP, and spBayes. Numerical results are reported in **Supplementary Table S2**.

**Supplementary Figure S4. Additional results from cell-type-specific SVG simulation analyses.**

(a). Distributions of four cell types in 4 different simulation scenarios constructed based on the mouse visual cortex dataset to evaluate ct-SVG methods (**Methods**). (b). Statistical power, false discovery rate (FDR), and area under the precision–recall curve (AUPRC) of Spacelink (ct-SVG) compared with Celina and STANCE for each of the four cell types across four simulation scenarios based on the human dorsolateral prefrontal cortex (DLPFC) dataset (left) and the mouse visual cortex dataset (right). Cell type proportions estimated by RCTD were used to apply each ct-SVG method. (c). Statistical power, FDR, and AUPRC of Spacelink (ct-SVG) compared with Celina and STANCE for each of the four cell types across four simulation scenarios based on the human DLPFC dataset (left) and the mouse visual cortex dataset (right), using oracle cell-type proportions derived from the true cell-type distributions. (d). Spacelink ct-ESV distributions for the focal cell type versus other cell types across four scenarios from the human DLPFC and mouse visual cortex datasets. (e). AUPRC of two variants of the Spacelink method versus Celina and STANCE for each of the four cell types across four simulation scenarios from the human DLPFC dataset (top) and the mouse visual cortex dataset (bottom). Spacelink variant 1 and variant 2 use the same data-driven multiple kernels as the original Spacelink (ct-SVG) but differ in how random effects from non-focal cell types are modeled: variant 1 excludes these effects completely (analogous to Celina), whereas variant 2 includes all of them (analogous to STANCE). Asterisks indicate significance from 1-sided pairwise t-tests comparing Spacelink (variant or original) with Celina or STANCE (*** P < 0.001, ** P < 0.01, * P < 0.05). (f). AUPRC of the Spacelink variants described in (e) for each of the four cell types across four simulation scenarios from the human DLPFC dataset (top) and the mouse visual cortex dataset (bottom). (g). Average F1 score of Spacelink across four cell types and four simulation scenarios in the human DLPFC and mouse visual cortex datasets, using varying cutoffs c_1_ and c_2_ to define the set of colocalized cell types, Coloc(r) (see **Methods**). Numerical results are reported in **Supplementary Table S2**.

**Supplementary Figure S5. Evaluation of p-value calibration for different SVG methods.**

(a). Q-Q plots (expected vs. observed p-values on –log10 scale) for different SVG methods across 10 scDesign3-based null simulations. (b). Q-Q plots for different SVG methods across 10 shuffling-based null simulations. (c). Kolmogorov-Smirnov (K-S) distance between observed p-value distributions and the uniform distribution for different methods and null simulation datasets under both scDesign3-based and shuffling-based frameworks. (d). Q-Q plots for Spacelink and SPARK-X on a high-sparsity null simulation. (e). Power and AUPRC curves comparing SPARK-X and Spacelink as a function of the level of sparsity. The gray area represents a range within 10% above and below the power of SPARK-X. (f). FDR, Power, AUPRC, and Kendall’s tau comparisons of SPARK-X and Spacelink across varying level of sparsity in the binned and un-binned versions of simulation datasets. Numerical results are reported in **Supplementary Table S2**.

**Supplementary Figure S6. Results of computational scalability and benchmarking Spacelink-lite against other SVG methods.**

(a). Runtime (seconds) and peak memory usage (GB) across different numbers of spots for SVG methods including Spacelink and Spacelink-lite. (b). FDR and statistical power comparisons between Spacelink and Spacelink-lite on both scDesign3-based and covariance-based simulations. (c). Scatter plots comparing ESV values estimated by Spacelink (Original) vs. Spacelink-lite (Approx.) across multiple covariance-based and scDesign3-based simulations. (d). FDR and power comparisons of nnSVG, SPARK-X, Moran’s I, and Spacelink-lite on covariance-based simulations using large-scale CosMx Cortex and Liver tissues. (e). Runtime (seconds) and peak memory usage (GB) across different numbers of spots for Celina, STANCE, and Spacelink (ct-SVG). Numerical results are reported in **Supplementary Table S2**.

**Supplementary Figure S7. Results of secondary analyses under different numbers of spots, levels of sparsity, and degrees of technical noise, benchmarking Spacelink ESV against other SVG metrics.**

(a). Cell type proportions in the kidney, prostate cancer, and brain cortex datasets, estimated by RCTD, along with an example of original data and its variations with 50% spots, 50% sparsity, and 50% technical noise. (b). Left: Line plots showing how ESV, propSV, FSV, and Moran’s I coefficient change across different percentage of spots in different datasets. Middle: Bar charts of proportion of genes showing less than 10%, 25%, and 50% change in each metric score at different reduction in the number of spots. Right: Boxplots of maximum change in each metric across different numbers of spots, for kidney, prostate cancer, and DLPFC datasets. (c). Line plots showing how ESV, PropSV, FSV, and Moran’s I coefficient change across different levels of sparsity in different datasets. (d). Left: Line plots showing how ESV, PropSV, FSV, and Moran’s I coefficient change across different degrees of technical noise in different datasets. Right: Boxplots of Kendall’s tau correlation between each metric score and noise level for kidney, prostate cancer, and DLPFC datasets. In (b) and (d), asterisks indicate significance from 1-sided pairwise t-tests comparing Spacelink against each method (*** P < 0.001, * P < 0.05). Numerical results are reported in **Supplementary Table S2**.

**Supplementary Figure S8. Secondary analyses benchmarking Spacelink against other SVG methods.**

(a). Statistical power and false discovery rate (FDR) of the original Spacelink and 5 variants of the Spacelink method adopting an expanded or alternative kernel set, such as Matérn kernels (p = 1 or 2), Gaussian or RBF kernels (equivalent to Matérn with p = ∞), or cosine periodic kernels, across 10 scDesign3-based simulations (b). Statistical power, FDR, and AUPRC of Spacelink versus global SVG methods in unbalanced datasets (**Methods**). (c). Adjusted Rand Index (top) and Normalized Mutual Information (bottom) of Spacelink ESV and nine other gene scoring metrics used to select top-ranked genes for domain detection of cortical layers in the human DLPFC Visium datasets^25^. Different boxplot colors indicate the number of top-ranked genes used. Asterisks indicate significance from 1-sided pairwise t-tests comparing Spacelink ESV with each other method (*** P < 0.001, ** P < 0.01, * P < 0.05). (d). Kendall’s tau comparisons between original Spacelink ESV, Spacelink ESV using 𝓁_1_-norm in weight parameters, and Spacelink ESV using Spectral norm in weight parameters. (e). Percentage of non-SVGs with ESV < 0.05 across 10 scDesign3-based simulations. (f). Spatial expression plots of two example simulated genes under the covariance-based model with σ = 1 and *τ* = 0.5 (left) and with *σ* = 1 and *τ* = 2 (right). (g). Scatter plots comparing estimated parameter values and Spacelink ESV between two scenarios: i) using original spatial coordinates and ii) using min-max scaled coordinates in the scDesign3-based simulation using DLPFC data. Numerical results are reported in **Supplementary Table S2**.

**Supplementary Figure S9. Comparison between Spacelink and Spacelink-lite on CosMx datasets, gene-specific spatial bandwidths estimated by Spacelink in three human tissues, cortical layer structure in the brain cortex, and examples of gene expression in CosMx and Visium brain cortex.**

(a). Confusion matrices comparing SVG classification results between Spacelink (after rasterization) and Spacelink-lite (with original dataset) for Cortex, Liver, and Lymph node CosMx datasets. (b). Heatmap of the weights (i.e., estimated variance components 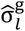) corresponding to gene-specific length scales inferred by the Spacelink model for each gene in brain cortex (left), lymph node (middle), and liver (right) tissues from CosMx (top) and Visium (bottom) datasets. Across spatial platforms, we observe greater heterogeneity in spatial length scales in lymph node and liver, compared to cortex. (c). Cortical layer structure of the brain cortex in the Visium dataset. (d). Spatial expression plots of three genes (*CNP, SCGB2A2, EIF5B*) illustrating distinct ESV patterns between CosMx and Visium in brain cortex (also highlighted in the scatter plot in **Figure 3d**). *CNP* and *EIF5B* exhibit consistently high and low ESV values across both platforms, whereas *SCGB2A2* shows high spatial variability in Visium but not in CosMx. Numerical results are provided in **Supplementary Table S3**.

**Supplementary Figure S10. Estimated cell type proportions in Visium human samples and cross-platform evaluation of ct-SVG methods in liver.**

(a). Cell type proportions in the brain cortex (top), lymph node (middle), and liver (bottom) Visium datasets, estimated by RCTD using the corresponding CosMx datasets as references. (b). Cross-platform consistency of cell-type-specific SVG (ct-SVG) methods (Spacelink, Celina, STANCE) between CosMx and Visium in the liver, evaluated by Spearman correlations of p-values for each cell type. Scatter plots show correlations across liver cell types, comparing Spacelink with Celina and STANCE. Numerical results are provided in **Supplementary Table S3**.

**Supplementary Figure S11. Additional results from cross-platform evaluation between matched tissue CosMx and Visium panel data.**

(a). Left: Spearman correlations of Spacelink ESV and five alternative metrics of spatial variability between CosMx and Visium data, restricted to tissue-relevant disease/trait genes in liver. Right: Average Spearman correlations of p-values from Spacelink, STANCE, and Celina across cell types between CosMx and Visium, also restricted to tissue-relevant disease/trait genes in liver (we did not benchmark ct-ESV as Celina and STANCE do not offer a comparable metric). Red circles denote values computed using all genes; other circles correspond to individual tissue-relevant complex diseases and traits. Asterisks indicate significance from 1-sided pairwise t-tests (*** P < 0.001, ** P < 0.01, * P < 0.05). (b). Spearman correlations of p-values from Spacelink and six other SVG methods between CosMx and Visium data, restricted to tissue-relevant disease/trait genes in brain cortex (top left), lymph node (top right), and liver (bottom). Red circles denote values computed using all genes; other circles correspond to individual tissue-relevant complex diseases and traits. Asterisks indicate significance from 1-sided pairwise t-tests (*** P < 0.001, ** P < 0.01, * P < 0.05). (c). Fractions of the union of the top 500 PoPS-prioritized genes for tissue-matched traits implicated as top N ranked global SVGs based on Spacelink ESV and 7 other spatial gene prioritization metrics applied to CosMx and Visium data from different tissues. We consider 4 values of N (100, 200, 500 and 1,000). Numerical results are provided in **Supplementary Table S3**.

**Supplementary Figure S12. Additional results from disease-informativeness analyses of gene programs in 3 CosMx human tissues – brain cortex, lymph node and liver.**

(a). Logistic regression analyses as in **Figure 4a**, but for Spacelink ct-ESV programs, conditioned on additional cell-type-level confounders (mean and variance of log-normalized expression within each cell type). (b). Logistic regression analyses as in **Figure 4a**, but using MAGMA-prioritized disease gene set (z-score > 3) for each tissue-matched trait, instead of the top 500 PoPS-prioritized disease gene set. (c). Logistic regression analyses as in **Figure 4a**, but using gene programs obtained from Spacelink ESV, nnSVG PropSV, and SpatialDE FSV. (d). Logistic regression analyses as in **Figure 4a**, but using gene programs obtained from Spacelink ESV and p-values from Spacelink and six other SVG identification methods. (e). Method-specific joint regression analyses of all nominally significant (p < 0.05) global and cell-type-level gene programs in each tissue, as in **Figure 4b**, but using gene programs obtained from Spacelink ESV and p-values from Spacelink and six other SVG identification methods. Relative AUPRC (against baseline AUPRC; see **Methods**) was used to evaluate each joint regression model with respect to each tissue-matched trait. Asterisks indicate significance from 1-sided pairwise t-tests (*** P < 0.001, ** P < 0.01, * P < 0.05). (f). Additional joint regression analyses comparing gene programs obtained from Spacelink-lite ESV, Spacelink ESV and p-value from SPARK-X. Asterisks indicate significance from 1-sided pairwise t-tests (*** P < 0.001) (g). Heatmap of disease- or trait-specific enrichment as in **Figure 4e**, but using MAGMA-prioritized disease gene set (z-score > 3) for each trait instead of the top 500 PoPS-prioritized disease gene set. Numerical results are provided in **Supplementary Table S4**.

**Supplementary Figure S13. Additional results from the application of Spacelink on the mouse embryonic Stereo-seq SRT dataset.**

(a). Number of of SVGs identified by Spacelink at each stage within the non-brain region (top), and average ESV at each stage within the whole embryo (second), brain region (third), and non-brain region (bottom), with bars shaded from light to dark green across developmental time. Error bars denote 95% confidence intervals. (b). Volcano plots of gene-level regressions associating stage-corrected mean of sc-transform normalized gene expressions (adjusted for embryo size; **Methods**) against developmental time in the whole embryo (left) and brain (right). (c). Volcano plots of gene-level regressions associating stage-corrected overall variance (spatial + non-spatial) of sc-transform normalized gene expressions (adjusted for embryo size) against developmental time in the whole embryo (left) and brain (right). In (b) and (c), each point represents the regression slope (x-axis) and –log10 p-value (y-axis). Genes with nominally significant (P < 0.05) negative and positive associations are shown in light and dark green, respectively. (d). Pathway enrichment analyses (based on ConsensusPathdb^74^) of genes with significantly negative (light green) and positive (dark green) associations between mean (top panel) or variance (bottom panel) of sc-transform normalized gene expressions and developmental progression in the whole embryo (left column) and brain region (right column). (e). Bar charts summarizing the proportion of gene-specific regression models satisfying normality and homoscedasticity assumptions, and the mean coefficient of determination (R^2^), for whole, brain, and non-brain region models. (f). Gene expressions of the example gene *Camp* across developmental stages (E9.5–E16.5) with ESV values annotated at each stage. (g). Average stage-corrected logit(ESV) within brain regions at each stage among genes with nominally significantly (P < 0.05) negative (light green) and positive (dark green) associations between spatial variability and developmental progression. (h). Disease informativeness of genes with negative (circles) or positive (triangles) temporal logit(ESV) associations in brain (top) and non-brain (bottom), focusing on four brain-related Mendelian disorders. Informativeness is measured by the fraction of disorder genes with significant ESV trends (X axis) and the odds ratio between gene sets (Y axis). Numerical results are reported in **Supplementary Table S5**.

**Supplementary Figure S14. Odds ratios based on overlap of perturbation programs in each cell type for each pair of 35 autism knockout genes in in-vivo Perturb-seq assay.**

Odds ratios of excess overlap between downstream-altered genes (perturbation programs) in each of 5 cell types, and across cell types, for each pair of 35 de-novo autism knockout (KO) genes in Jin et al.^86^. The magnitude of overlap is represented by odds ratio (dot size), and significance by −log10(p-value) (dot color). Panels show results across all cell types (top left), astroglia (top middle), excitatory neurons (top right), inhibitory neurons (bottom left), microglia (bottom middle), and oligodendrocytes (ODCs; bottom right). Numerical results are reported in **Supplementary Table S6**.

**Supplementary Figure S15. Additional results for Spacelink application on downstream perturbation programs in mouse in vivo perturbations of autism risk genes.**

(a). Effective Spatial Variability (ESV) of the 35 autism risk genes that were knocked out (KO) in the vivo Perturb-seq assay^86^ across mouse organogenesis stages. (b). Average ESV of the KO genes (green) and downstream perturbation program genes in astroglia (dark blue), excitatory neurons (light blue), inhibitory neurons (red), microglia (yellow), and oligodendrocytes (orange) across organogenesis stages. Error bars denote 95% confidence intervals. (c). Average stage-corrected ESV of perturbation program genes in specific pairs of top three cell types with the strongest odds ratios of overlap in **Figure 6b**. Bars are shaded from light to dark green across developmental time. The left panel shows genes altered in astroglia and excitatory neurons but not microglia; the middle panel, in excitatory neurons and microglia but not astroglia; and the right panel, in astroglia and microglia but not excitatory neurons. Error bars denote 95% confidence intervals. (d) For each of the eight organogenesis stages, correlation between brain-specific and non–brain-specific mean (top) or variance (bottom) of sc-transform normalized gene expression across genes from three broad classes: (i) union of perturbation program genes in excitatory neurons and astroglia, (ii) union of perturbation program genes in inhibitory neurons, microglia, and oligodendrocytes, and (iii) all genes. (e). ESV of perturbation program genes in astroglia (top left), excitatory neurons (top right), inhibitory neurons (bottom left), microglia (bottom middle), and oligodendrocytes (bottom right) across eight organogenesis stages, stratified by the number of perturbed genes affecting them. Thick lines denote average ESV; shaded areas indicate confidence bands. The red dashed line shows the overall trend estimated by linear regression. Asterisks denote significance of Pearson correlation (*** P < 0.001). Numerical results are reported in **Supplementary Table S6**.

**Supplementary Figure S16. Additional results integrating Spacelink with perturbation programs in mouse in vivo perturbations of autism risk genes.**

(a) Scatter plots of the average ESV of (i) all genes in genetically autism disease-associated gene set versus (ii) genes in the gene set that are in a perturbation program for some autism risk gene knockout (KO) in the in-vivo Perturb-seq assay. The disease-associated gene sets include (i) the top 100 MAGMA-prioritized polygenic genes (left), (ii) the top 100 PoPS-prioritized polygenic genes (middle), and (iii) 108 monogenic autism risk genes^84,85^ (right). (b). Top: Scatter plots of the average mean of sc-transform normalized gene expression of (i) perturbation programs genes per perturbation in each stage versus (ii) subset of genes from (i) that overlap either of the three autism-associated gene sets. Bottom: Scatter plots of the overall mean of sc-transform normalized expression of (i) all genes in genetically autism disease-associated gene set versus (ii) genes in the gene set that are in a perturbation program for some autism risk gene knockout (KO) in the in-vivo Perturb-seq assay. (c). Top: Scatter plots of the average overall variance (spatial + non-spatial) of sc-transform normalized expression of (i) perturbation programs genes per perturbation in each stage versus (ii) subset of genes from (i) that overlap either of the three autism-associated gene sets. Bottom: Scatter plots of the average overall variance of sc-transform normalized expression of i) all genes in genetically autism disease-associated gene set versus (ii) genes in the gene set that are in a perturbation program for some autism risk gene knockout (KO) in the in-vivo Perturb-seq assay. (d). Density plots of perturbation program genes intersecting each autism-associated gene set, stratified by the number of perturbations affecting them. Shown are results across all cell types (top), as well as within astroglia (second), excitatory neurons (third), inhibitory neurons (fourth), microglia (fifth), and oligodendrocytes (bottom). Genes intersecting the top 100 MAGMA-prioritized polygenic genes are shown in red, those intersecting the top 100 PoPS-prioritized polygenic genes in blue, and those intersecting the 108 monogenic autism risk genes in yellow. In (a-c), red lines indicate the fitted regression line from a linear model with zero intercept; slopes are reported. Numerical results are reported in **Supplementary Table S6**.

**Supplementary Figure S17. Additional results from the application of Spacelink to 10x Visium human DLPFC dataset from 32 samples at different stages of Alzheimer’s disease.**

(a). Boxplots of maximum normalized distances of tissue spots from each of 32 human samples in the 10x DLPFC dataset and 8 mouse samples in the Stereo-seq organogenesis dataset. For each sample, the maximum normalized distance was calculated by dividing all pairwise spot distances by the minimum distance and then taking the maximum of these normalized values. (b). Boxplots of the number of spatially variable genes (SVGs) (top panel) and the corrected or uncorrected average ESV (bottom panel) across genes at control and three stages of AD pathology, defined by amyloid burden (first column), tangle accumulation (second column), and neuritic plaque counts (third and fourth columns). For amyloid burden and tangle accumulation, CTRL includes 4 control samples from 2 healthy donors; AD-Lower includes 10 samples with the lowest values of the pathology indicator; AD-Mid includes 9 samples with intermediate values of the pathology indicator; and AD-Higher includes 9 samples with the highest values of the pathology indicator. For neuritic plaque counts, CTRL includes 4 control samples from 2 healthy donors; AD-Lower includes 8 samples with the lowest counts; AD-Mid includes 10 samples with intermediate counts; and AD-Higher includes 10 samples with the highest counts. The difference in sample numbers between the two classifications (4, 10, 9, 9 vs. 4, 8, 10, 10) arises because some samples had identical neuritic plaque counts, requiring their assignment to the same group. (c). Volcano plots of regression analyses associating ESV with neuritic plaque count at the gene level, adjusting for cortical layer composition, mean expression levels, and donor-level random effects. Each point represents the regression slope coefficient and its –log p-value for a gene. Genes nominally significant (P < 0.05) are highlighted in green. (d). Pathway enrichment analysis of 115 genes with nominally significant (P < 0.05) negative associations between spatial variability (ESV) and neuritic plaque count. Enrichment was performed using ConsensusPathDB^74^ with the Visium panel genes as background. (e). Spatial expression plots of *YWHAG* (left column), with layer annotations (middle column) and plaque annotations (right column) in one control sample (top panel) and one high–tangle-accumulation sample (bottom panel). (f). Spatial expression plots of *SNCB* (left column), with layer annotations (middle column) and plaque annotations (right column) in one control sample (top panel) and one high–neuritic-plaque-count sample (bottom panel). (g). Volcano plots of regression analyses associating ESV with amyloid burden (left), tangle accumulation (middle), and neuritic plaque count (right) at the gene level, adjusting for cortical layer composition, mean expression levels, variance of expression levels, and donor-level random effects. Each point represents the regression slope coefficient and its –log p-value for a gene. Genes nominally significant (P < 0.05) are highlighted: amyloid (blue), tau burden (orange), and neuritic plaque counts (green). (h). Pathway enrichment analyses of 260, 166, and 103 genes with nominally significant (P < 0.05) negative associations between spatial variability (ESV) and amyloid burden (left), tangle accumulation (middle), and neuritic plaque count (right). Enrichment was performed using ConsensusPathDB with the Visium panel genes as background. (i). Bar charts showing the proportion of gene-specific regression models satisfying normality and homoscedasticity assumptions, and mean R^2^ values, for models with and without variance of expression as a covariate, across all three AD pathology measures. Numerical results are reported in **Supplementary Table S7**.

**Supplementary Figure S18. Additional results from the application of Spacelink to 10x Visium human DLPFC dataset from 32 samples at different stages of Alzheimer’s disease.**

(a). Volcano plots of regression analyses associating the overall mean of sc-transform normalized expression values with amyloid burden (left), tangle accumulation (middle), and neuritic plaque count (right) at the gene level, adjusting for cortical layer composition and donor-level random effects. Each point represents the regression slope coefficient and its –log p-value for a gene. Nominally significant genes (P < 0.05) are highlighted in blue (amyloid), orange (tau), and green (plaques). (b). Pathway enrichment analyses of 1,051 and 745 genes negatively associated with tangle accumulation (left) and neuritic plaque count (right) (P < 0.05), using ConsensusPathDB with Visium panel genes as background. (c). Boxplots of the number of SVGs (top) and average ESV (bottom) within each cortical layer in control and three AD groups defined by amyloid burden, tangle accumulation, or neuritic plaque counts. Group definitions (CTRL, AD-Lower, AD-Mid, AD-Higher) follow **Supplementary Figure S17b**. (d). Spatial plots of *PGK1* (left) and *GPI* (right) in 10x Visium human DLPFC data (top) and mouse SRT data (bottom). For human data, expression, layer, and plaque annotations are shown for one control and one high–amyloid-burden sample. For mouse data, expression and layer annotations are shown for 4- and 12-month-old samples. (e). Spatial plots of *MEF2C* in human DLPFC (top) and mouse SRT (bottom). For human data, expression, layer, and plaque annotations are shown for one control and one high–amyloid-burden sample. For mouse data, expression and layer annotations are shown for 4- and 12-month-old samples. Numerical results are reported in **Supplementary Table S7**.

