## Supplementary Figures for "Mapping disease critical spatially variable gene programs by integrating spatial transcriptomics with human genetics"

### Supplementary Figure S1

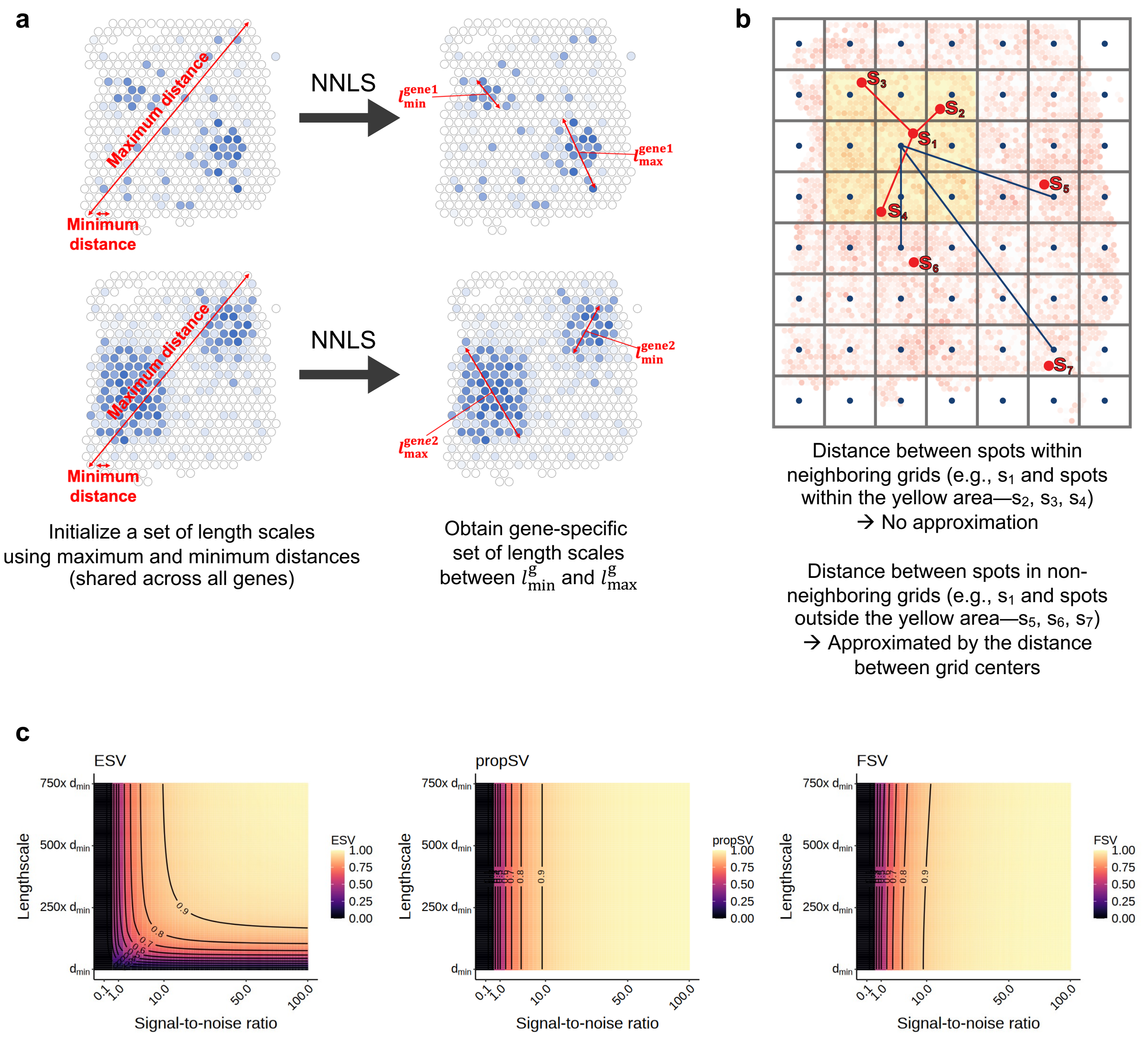

#### Supplementary Figure S2

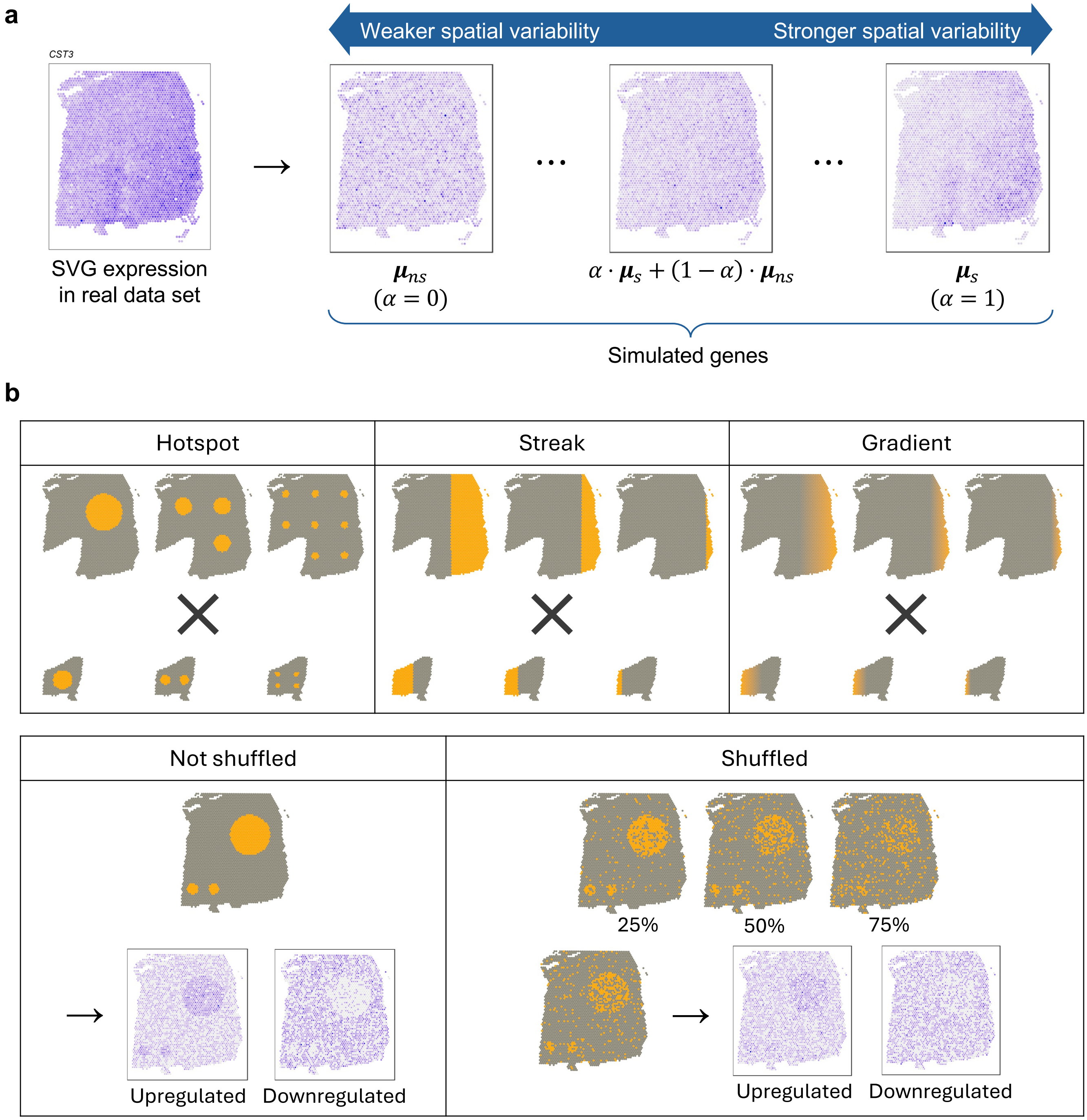

### Supplementary Figure S3

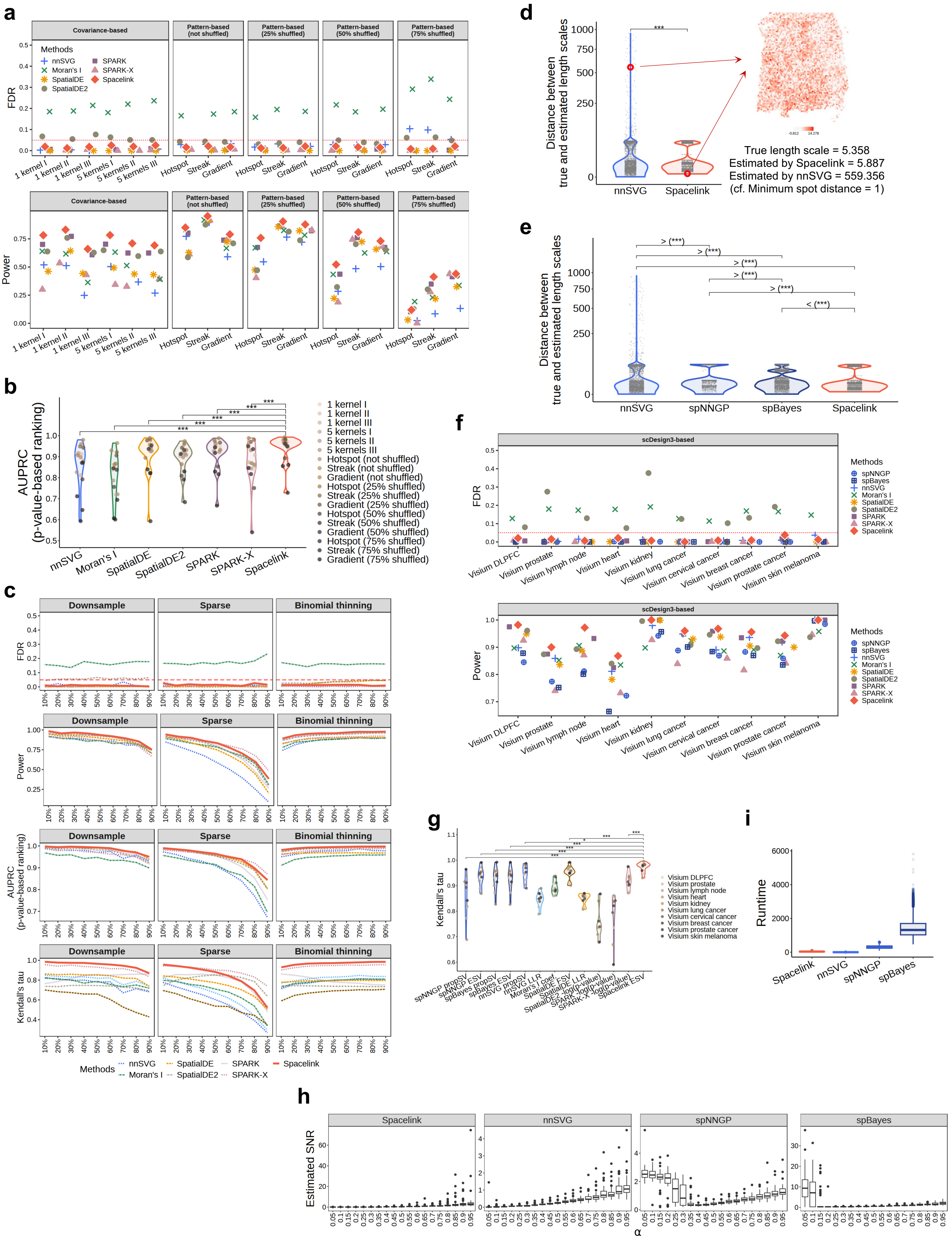

### Supplementary Figure S4

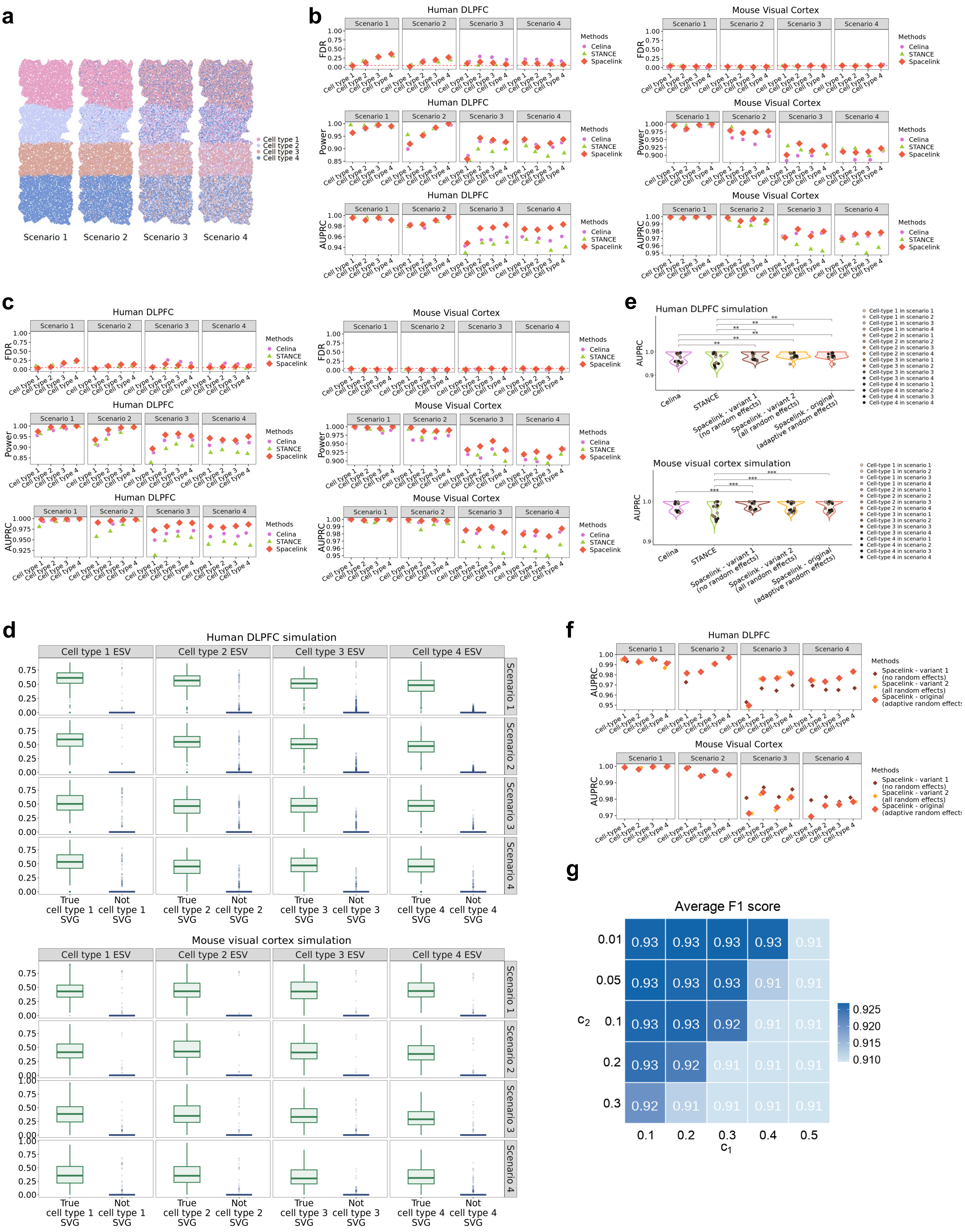

### Supplementary Figure S5

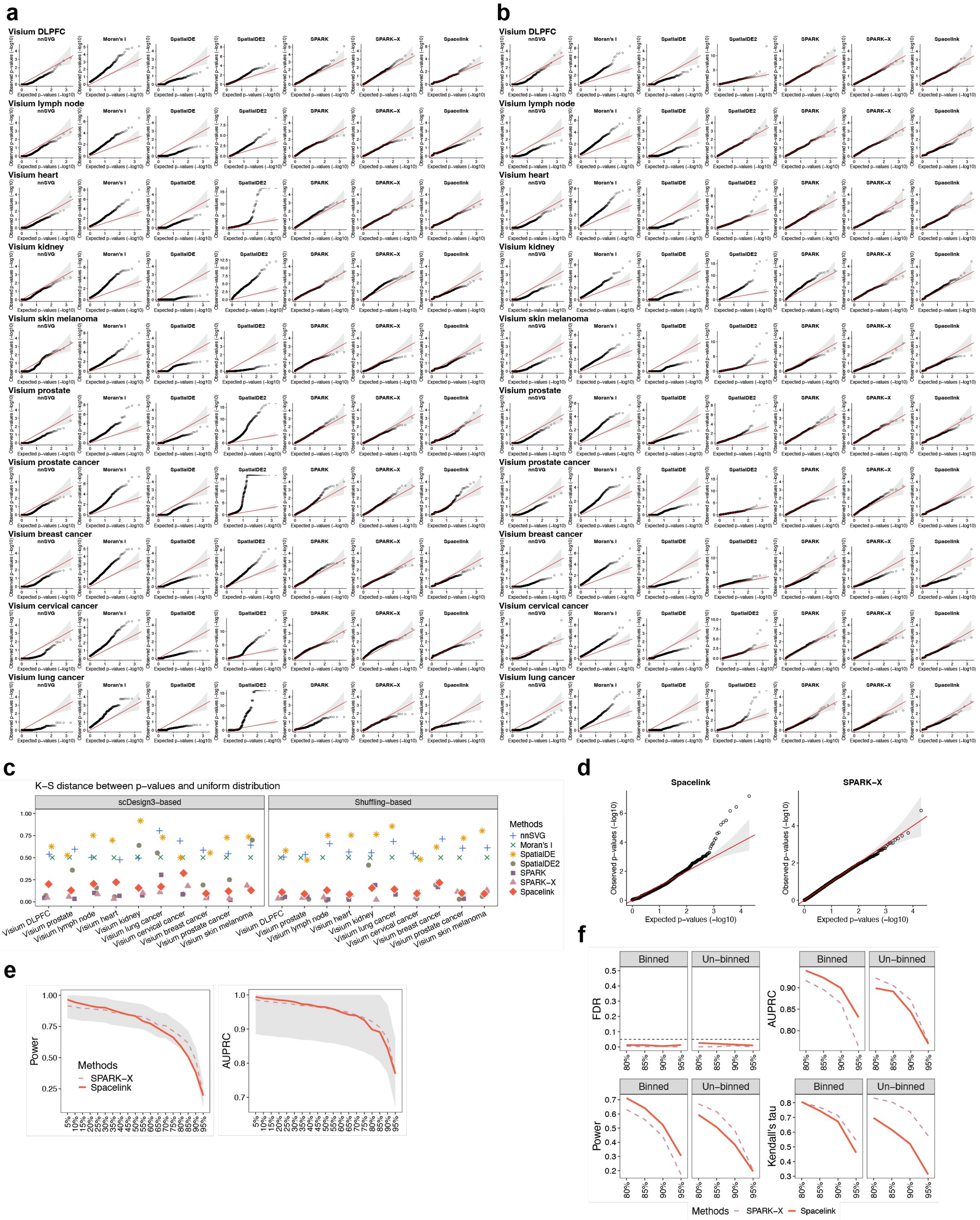

#### Supplementary Figure S6

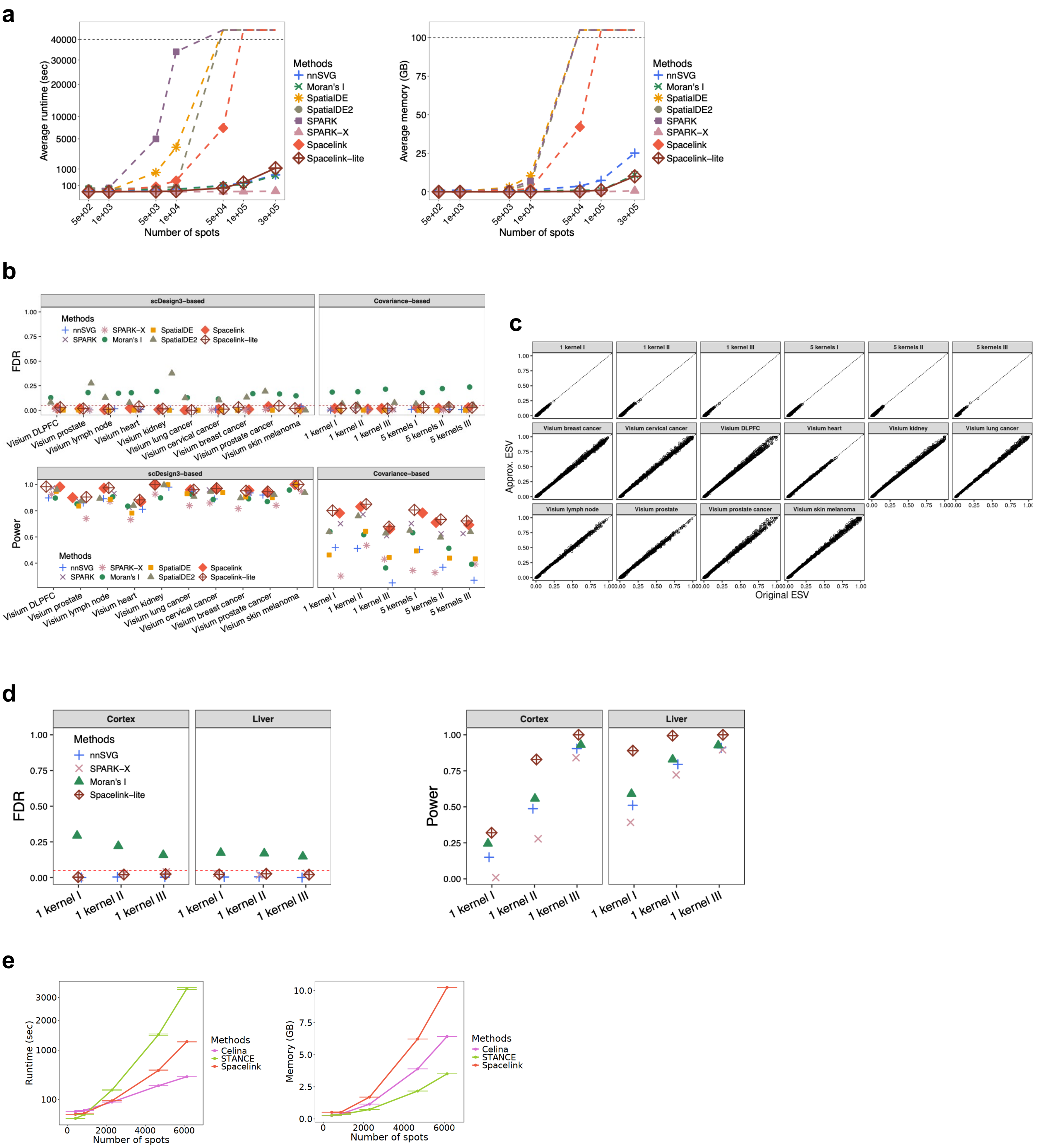

### Supplementary Figure S7

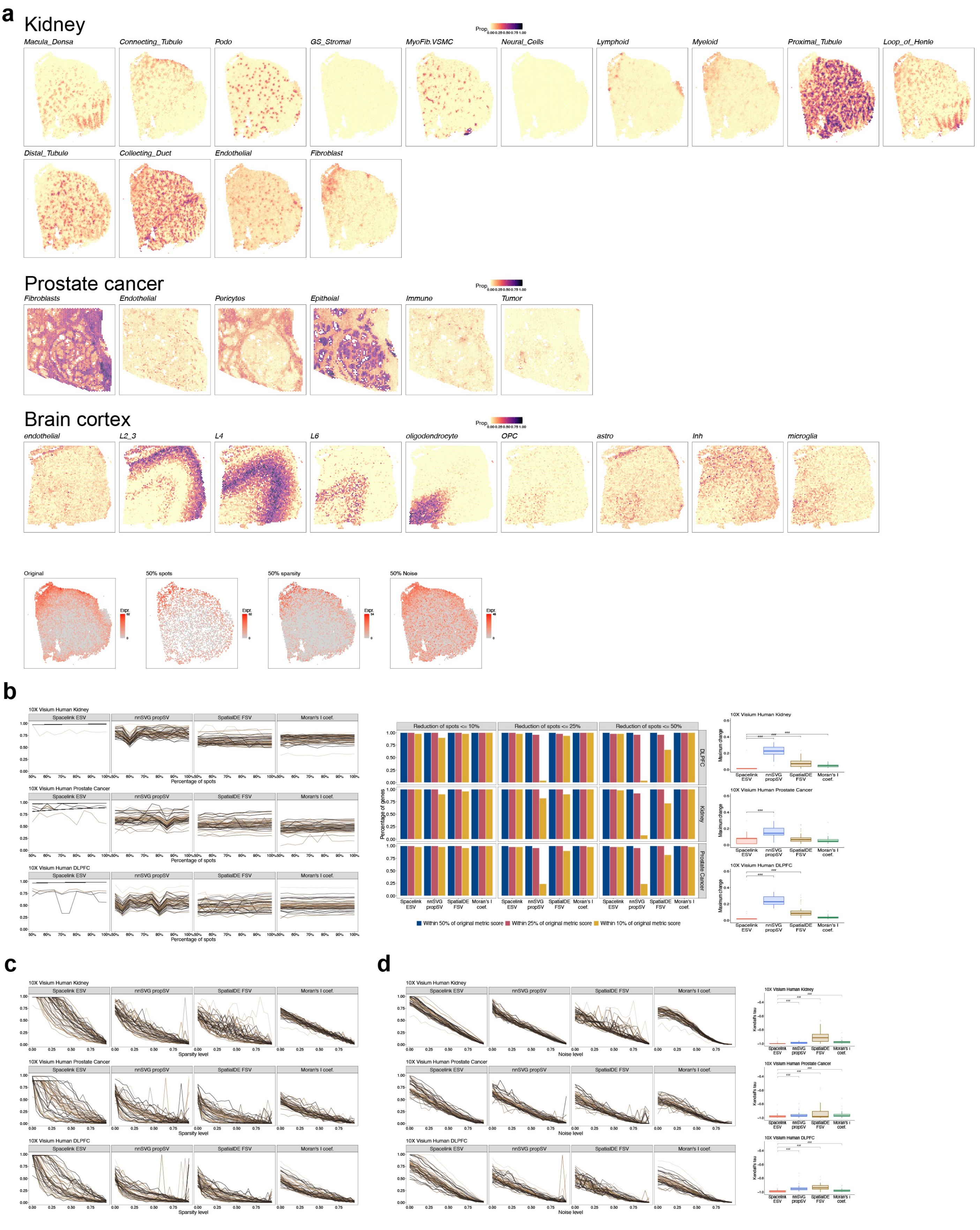

### Supplementary Figure S8

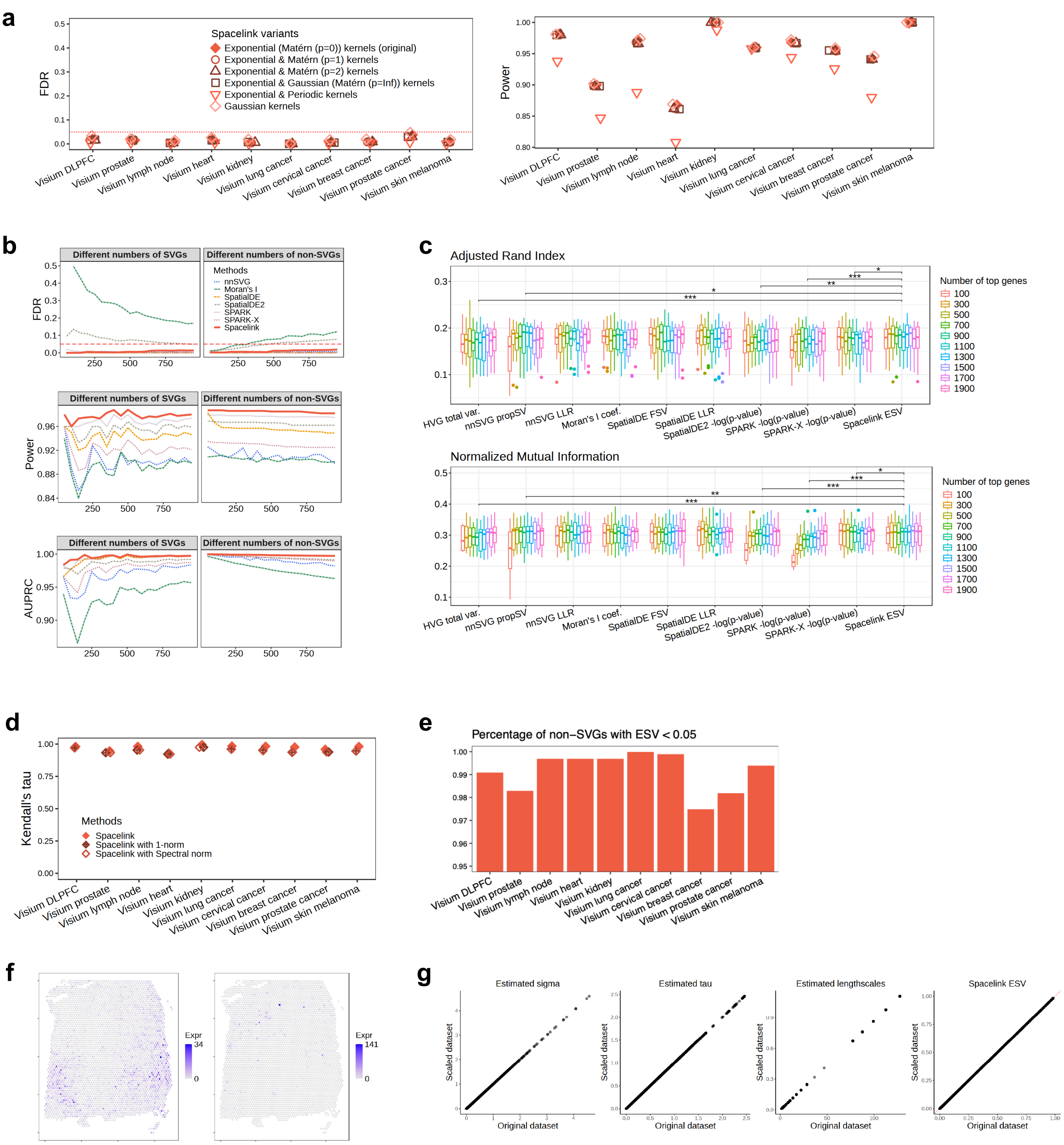

### Supplementary Figure S9

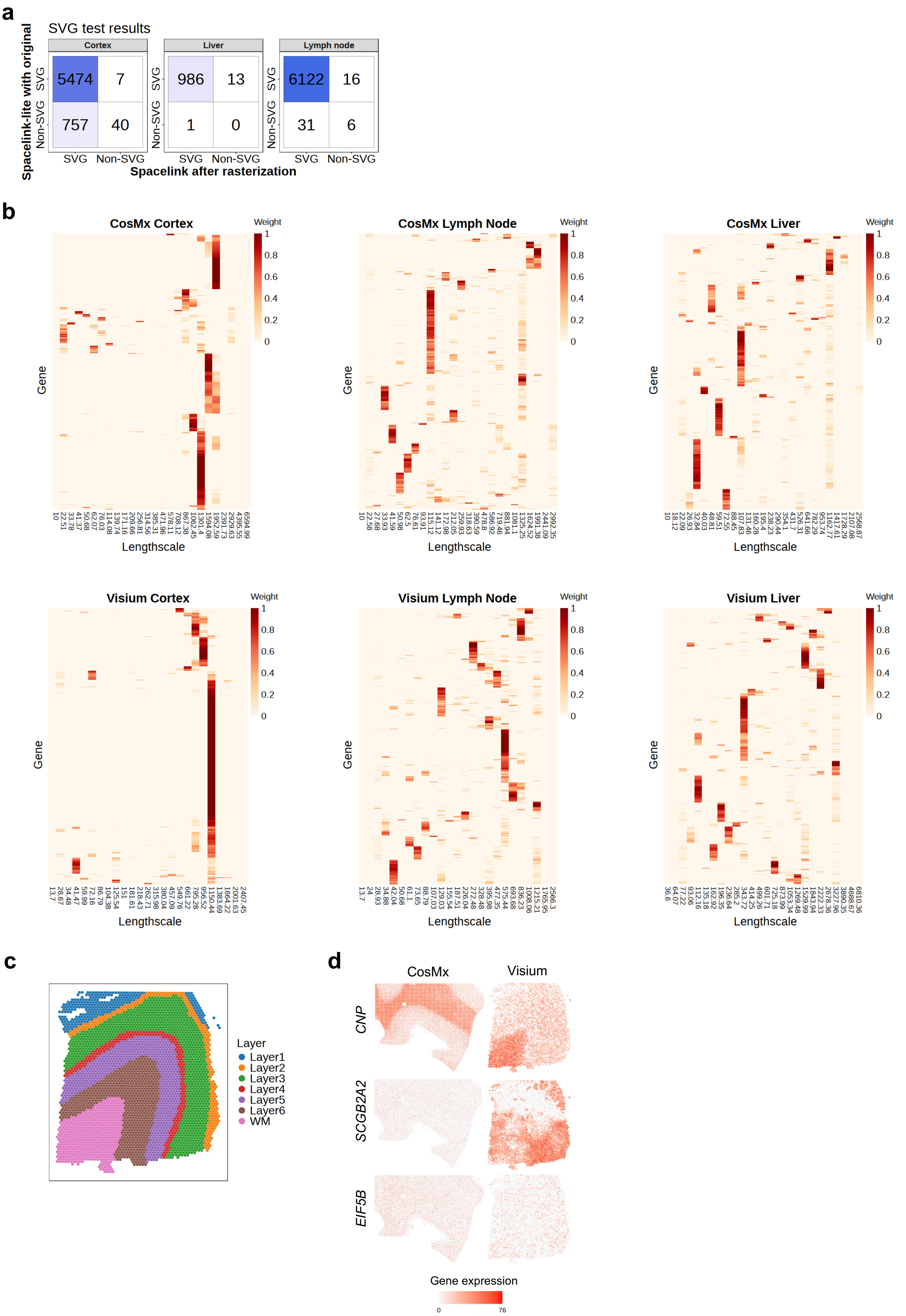

#### Supplementary Figure S10

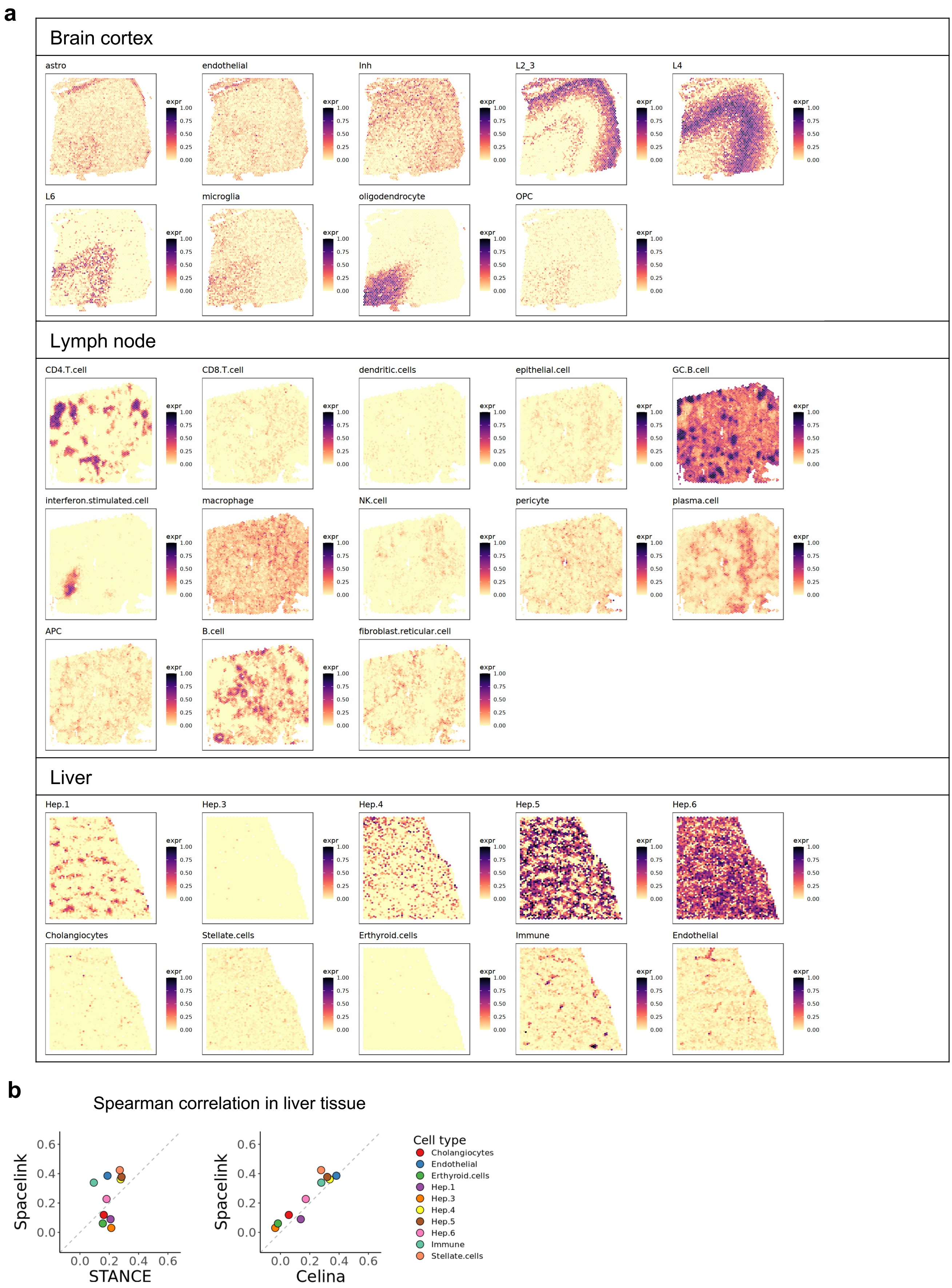

### Supplementary Figure S11

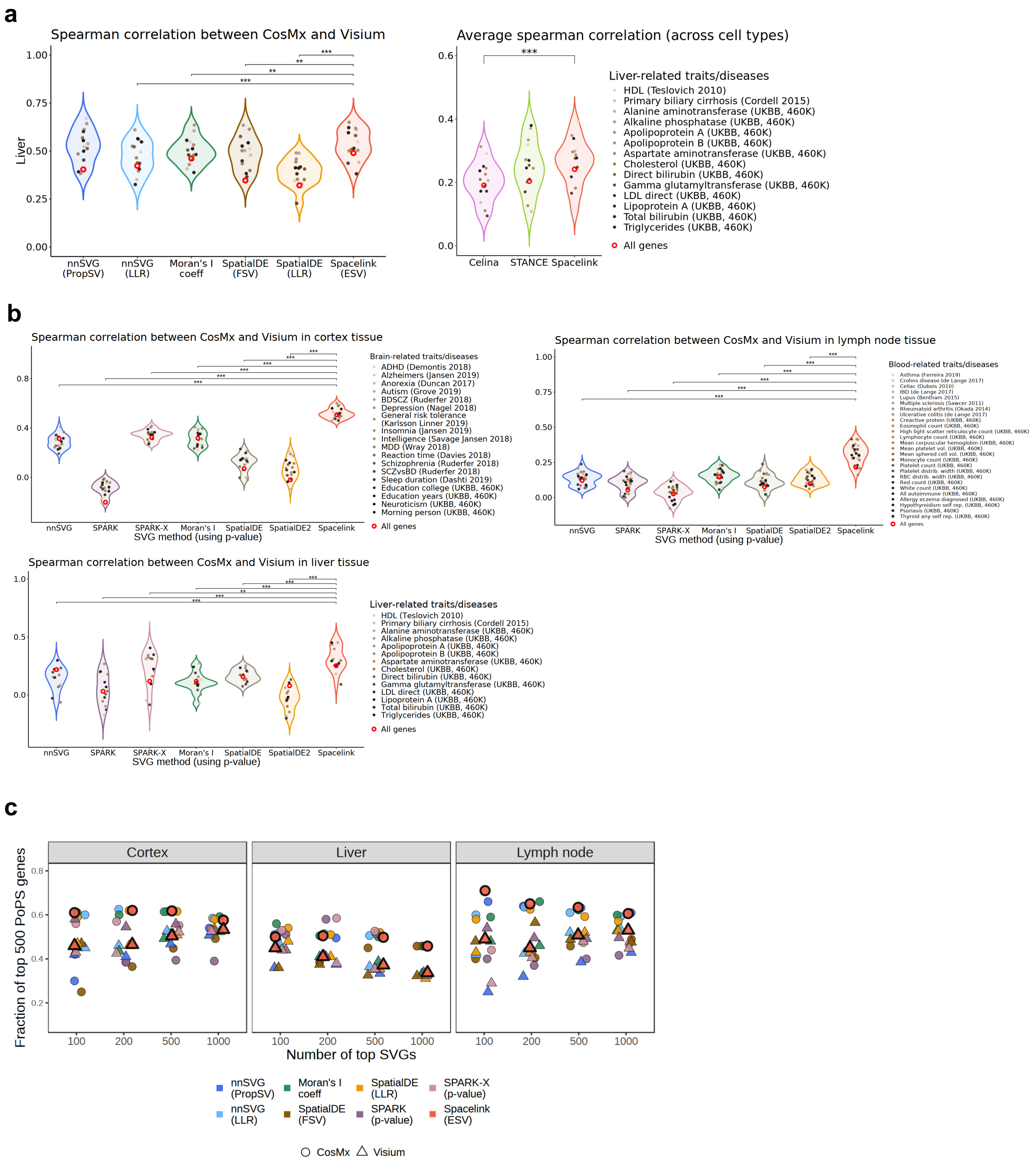

### Supplementary Figure S12

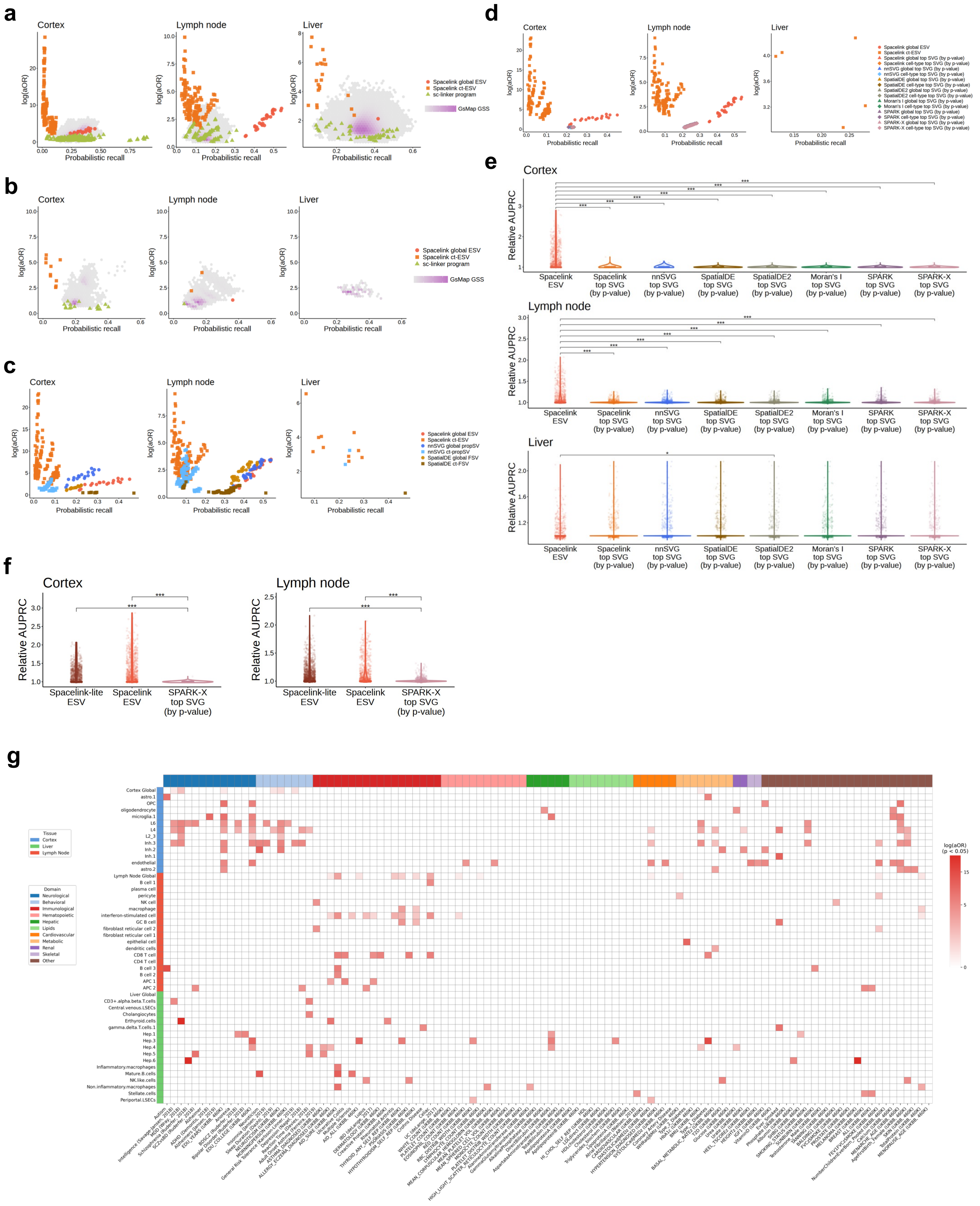

### Supplementary Figure S13

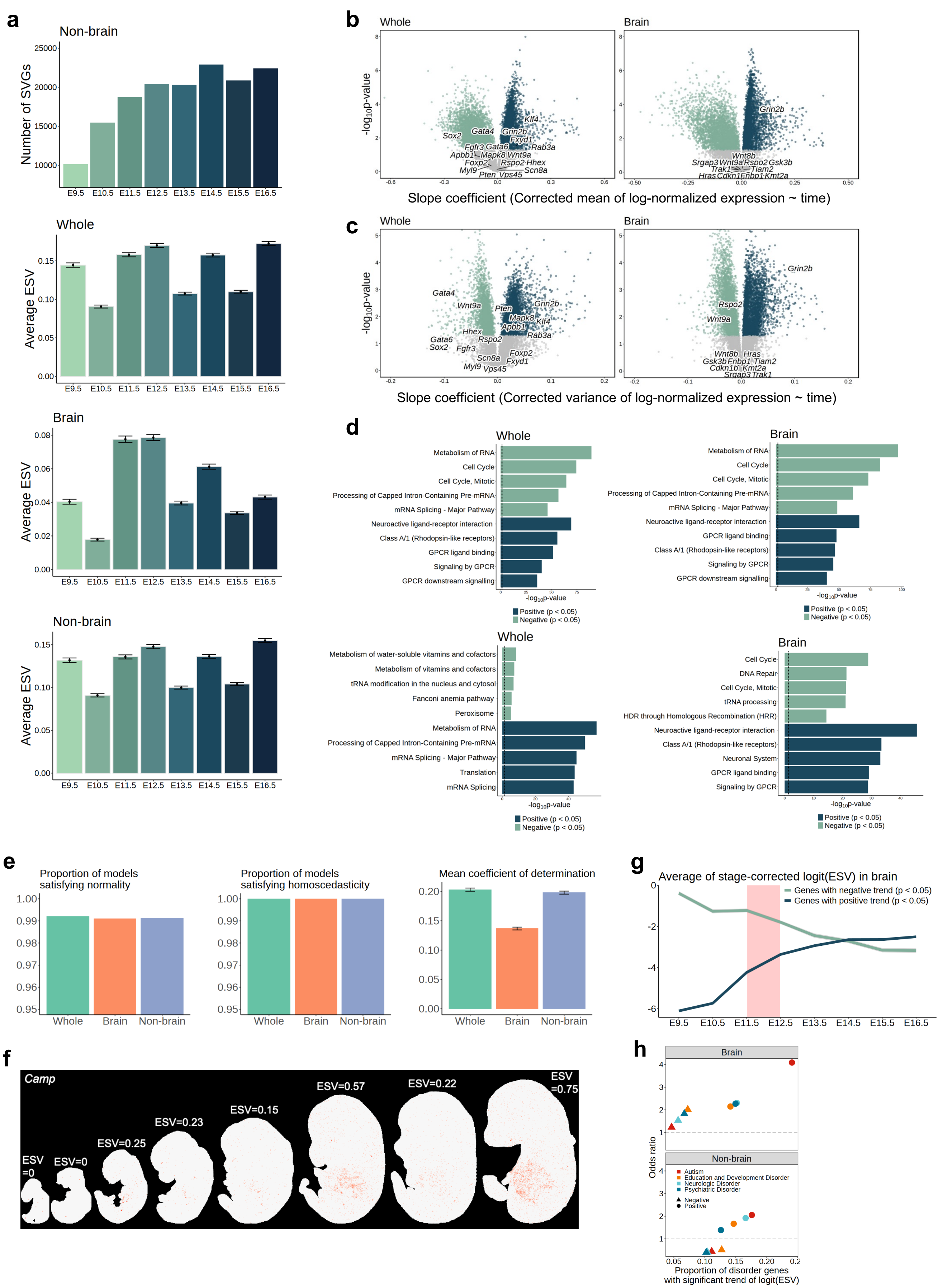

### Supplementary Figure S14

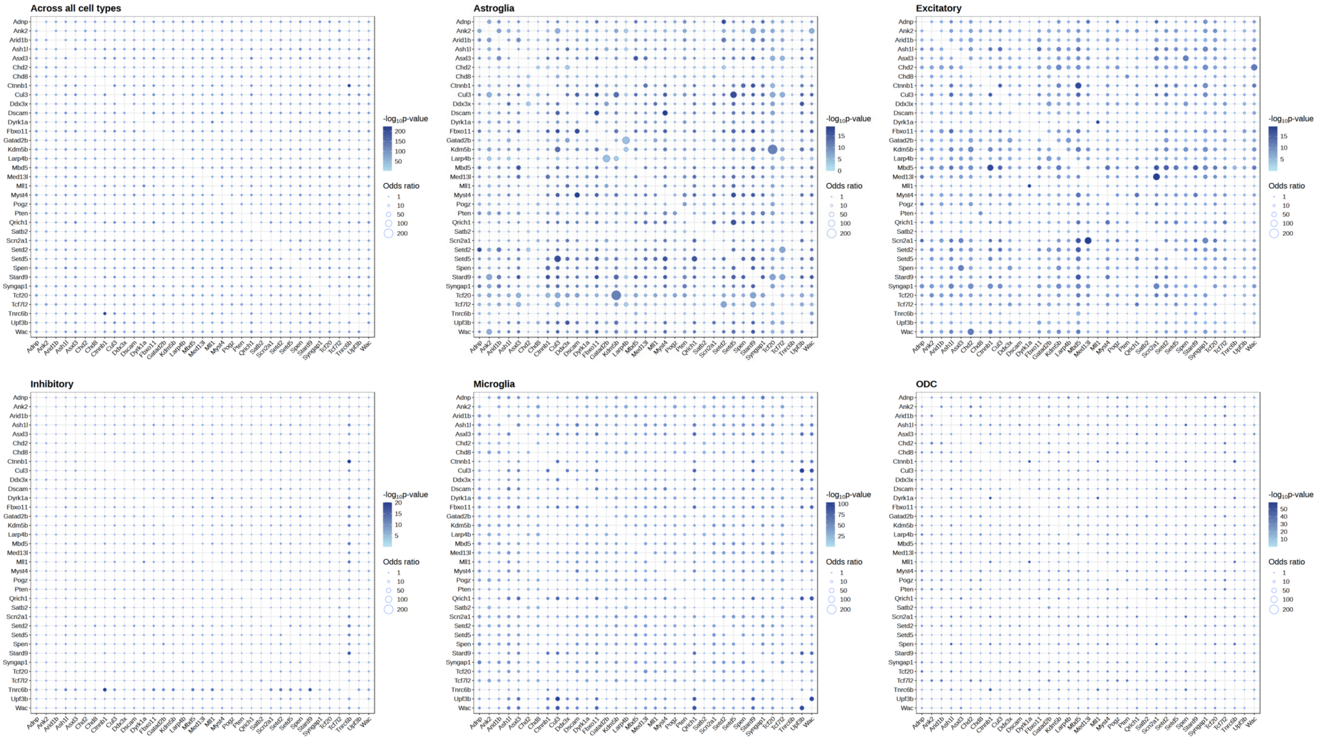

### Supplementary Figure S15

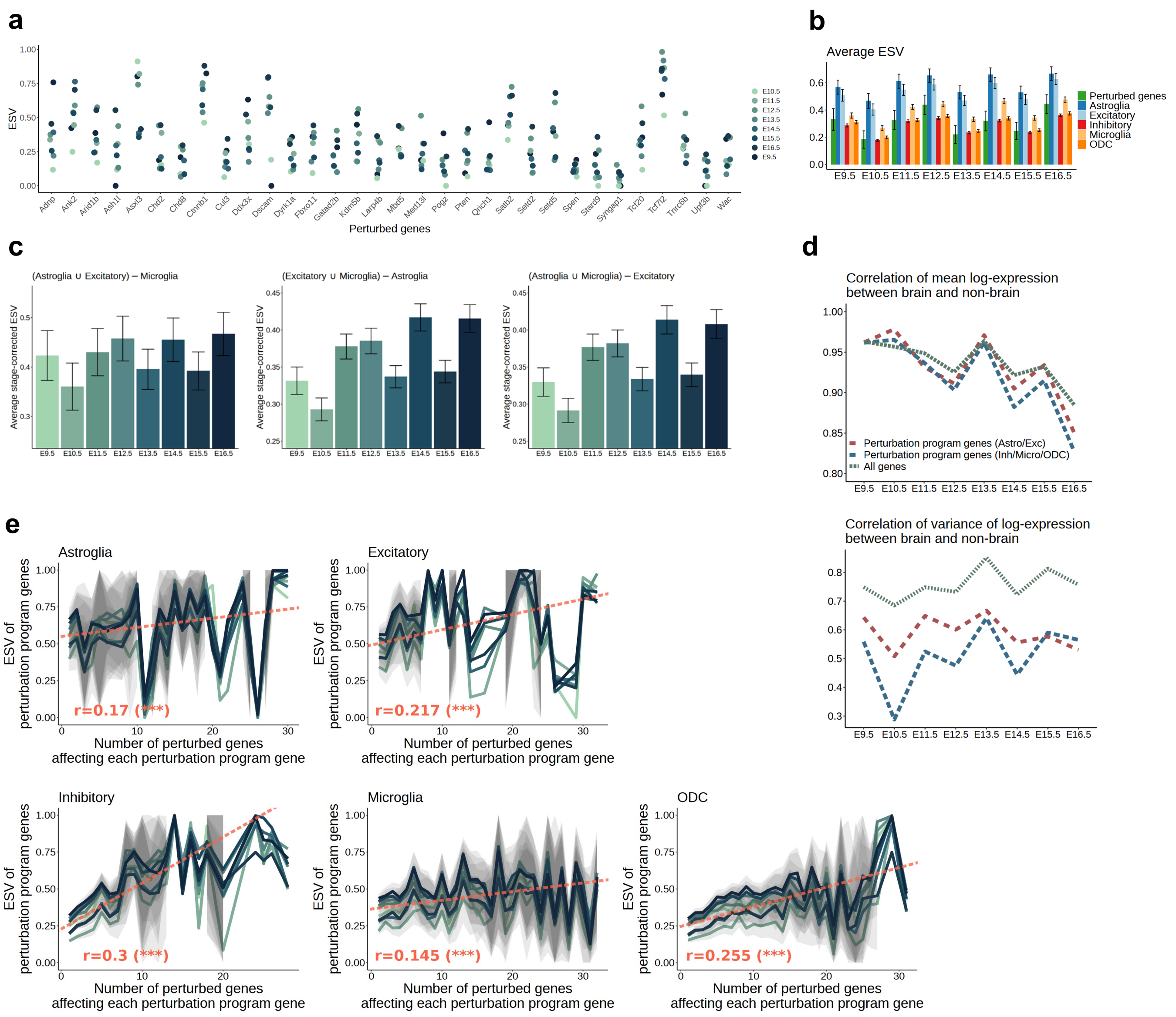

### Supplementary Figure S16

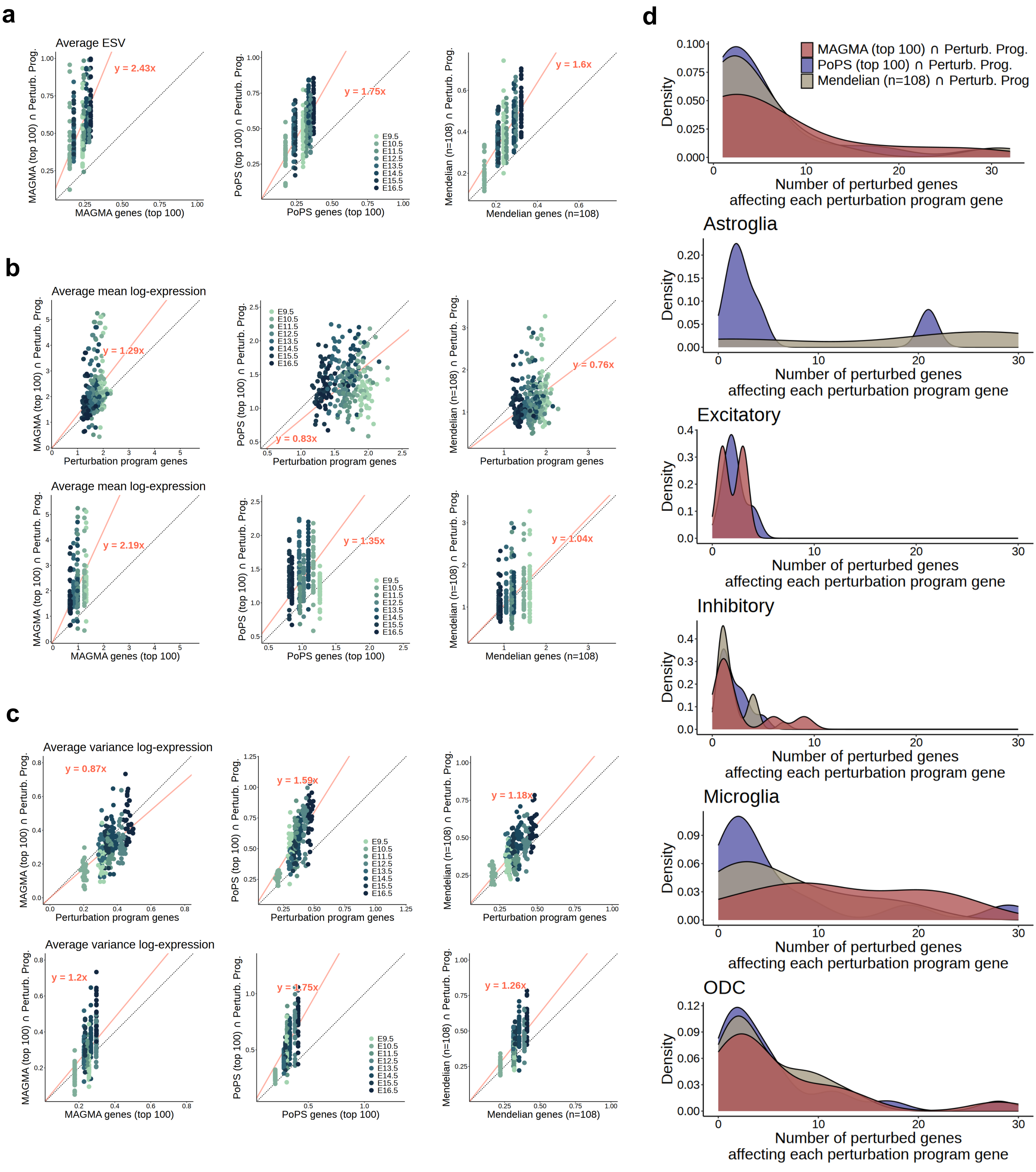

### Supplementary Figure S17

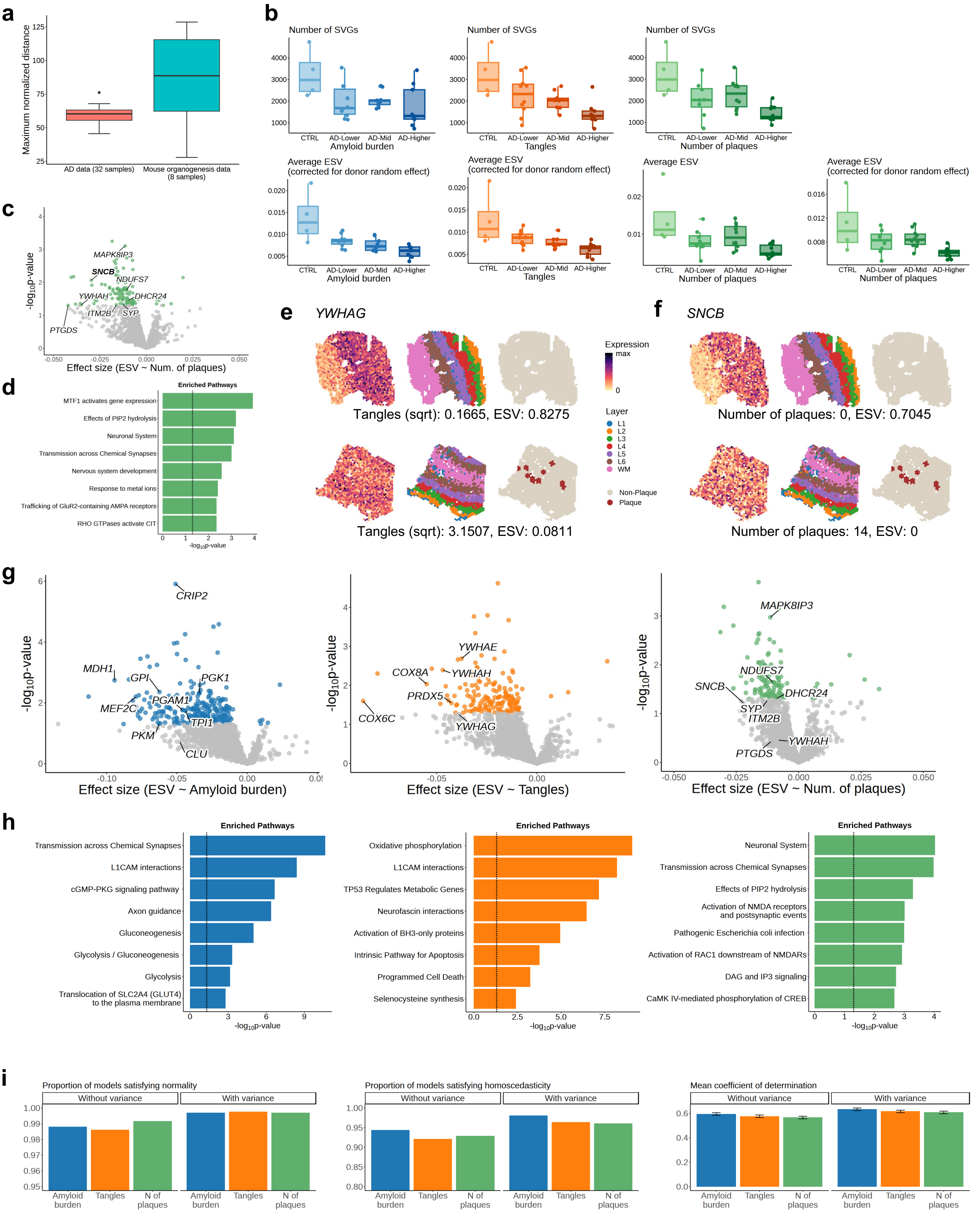

### Supplementary Figure S18

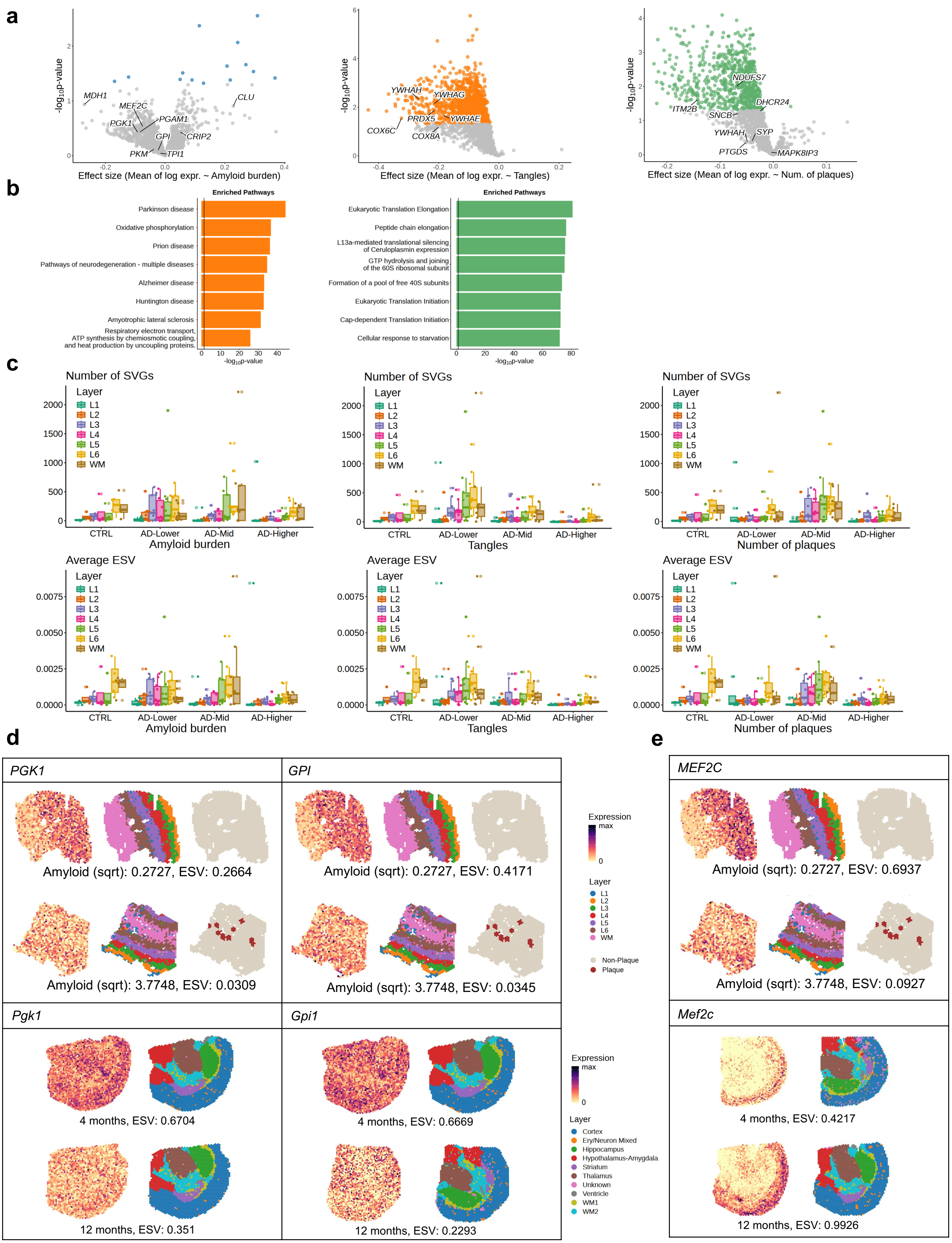
