## Supplementary Note for "Mapping disease critical spatially variable gene programs by integrating spatial transcriptomics with human genetics"

#### 1 Interpretation of Kernel Function

Our kernel function can be interpreted as an approximation of any completely monotone isotropic kernel function by using Bernstein's theorem. Consider an isotropic kernel function $K^*(d)$, defined as a function of distance $d$. If it is completely monotone, by Bernstein's theorem, it can be written as a Laplace transform of a Borel measure $\mu$, that is,

$$K^*(d) = \int_0^\infty \exp(-td) d\mu(t).$$

In practical applications, the Lebesgue integral representation can be approximated using a finite sum of simple functions. Specifically, we partition the domain $[0, \infty)$ into intervals $[t_{l-1}, t_l)$ for $l = 1, \dots, L$, where $t_0 = 0$ and $t_L = \infty$, and approximate $\exp(-td)$ on each interval as a constant $\exp(-t_l d)$. This gives the approximation:

$$K^*(d) \approx \sum_{l=1}^{L} \exp(-t_l d) \mu([t_{l-1}, t_l)).$$

Using the notations $b_l = \frac{1}{t_l}$ and $\sigma_l = \mu([t_{l-1}, t_l))$, the RHS becomes:

$$K^*(d) \approx \sum_{l=1}^{L} \sigma_l \exp(-d/b_l).$$

#### 2 Hypothesis Testing

##### 2.1 Hypothesis Testing for Global SVGs

We denote $\boldsymbol{y}$ as the vector of normalized, log-transformed expression values of gene $g$ across $N$ spatial locations, $\boldsymbol{s} = (\boldsymbol{s}_1, \dots, \boldsymbol{s}_N)$ (for simplicity, we omit the index $g = 1, \dots, G$). At the whole-tissue level, we model $\boldsymbol{y}$ with a Gaussian process as follows:

$$\boldsymbol{y} \sim \mathcal{N}\left(\boldsymbol{X\beta}, \sum_{l=1}^{L} \sigma_l \boldsymbol{K}_l + \tau \boldsymbol{I}\right),$$

where $[\boldsymbol{K}_l]_{i,j} = \exp\big(-\frac{\|\boldsymbol{s}_i - \boldsymbol{s}_j\|}{b_l}\big)$.

To assess whether each gene exhibits spatial variability, we test the following hypotheses:

$$H_0 : \sigma_l = 0 \text{ for all } l = 1, \dots, L$$
$$H_a : \text{At least one } \sigma_l \text{ is not zero.}$$

Here, we apply the score test to each variance component and combine the resulting p-values using the Cauchy combination method. Specifically, the score test statistic for variance component $\sigma_l$ is given by

$$S_l(\boldsymbol{\beta}, \tau) = \frac{1}{\tau} (\boldsymbol{y} - \boldsymbol{X\beta})^\top \boldsymbol{K}_l (\boldsymbol{y} - \boldsymbol{X\beta}).$$

Under the null hypothesis, the distribution of $S_l(\boldsymbol{\beta}, \tau)$ is a mixture of chi-square distributions [ZL03]. To simplify computation, we use the Satterthwaite approximation to represent it by a scaled chi-square distribution, $\kappa_l \chi^2(\nu_l)$, where

$$\kappa_l = \frac{tr(\boldsymbol{K}_l^2)}{tr(\boldsymbol{K}_l)} \quad \text{and} \quad \nu_l = \frac{\big(tr(\boldsymbol{K}_l)\big)^2}{tr(\boldsymbol{K}_l^2)}.$$

Since the parameters $\boldsymbol{\beta}$ and $\tau$ are unknown, we estimate them by their MLEs under the null hypothesis:

$$\hat{\boldsymbol{\beta}} = (\boldsymbol{X}^\top \boldsymbol{X})^{-1} \boldsymbol{X}^\top \boldsymbol{y}, \quad \hat{\tau} = \frac{(\boldsymbol{y} - \boldsymbol{X}\hat{\boldsymbol{\beta}})^\top (\boldsymbol{y} - \boldsymbol{X}\hat{\boldsymbol{\beta}})}{N}.$$

Using the estimated score test statistic $S_l(\hat{\boldsymbol{\beta}}, \hat{\tau})$, we compute the approximate p-value for each variance component as

$$\mathbb{P}\bigg( \frac{S_l(\hat{\boldsymbol{\beta}}, \hat{\tau})}{\kappa_l} > \chi^2_{\nu_l} \bigg).$$

After obtaining these p-values across all $l = 1, \ldots, L$, we combine them using the Cauchy combination method.

#### 2.2 Hypothesis Testing for Cell-Type Specific SVGs

To assess the cell-type-specific spatial variability of a gene expression $\boldsymbol{y}$ for a focal cell type $r$, we model it as follows:

$$\boldsymbol{y} \sim \mathcal{N}\Bigg( \tilde{\boldsymbol{X}}\boldsymbol{\beta}, \sum_{l=1}^{L} \sigma_{r,l} \boldsymbol{\Pi}_r \boldsymbol{K}_l \boldsymbol{\Pi}_r + \sum_{r' \in Coloc(r)} \sigma_{r',m} \boldsymbol{\Pi}_{r'} \boldsymbol{K}_m \boldsymbol{\Pi}_{r'} + \tau \boldsymbol{I} \Bigg).$$

To identify whether each gene is spatially variable within the cell type $r$, we test the following hypotheses:

$$H_0 : \sigma_{r,l} = 0 \text{ for all } l = 1, \ldots, L$$
$$H_a : \text{At least one } \sigma_{r,l} \text{ is not zero.}$$

In this case, we define the score test statistic for variance component $\sigma_{r,l}$ as follows:

$$S_{r,l}(\boldsymbol{\beta}, \{\sigma_{r',m}; r' \in Coloc(r)\}, \tau)(= S_{r,l}) = (\boldsymbol{y} - \tilde{\boldsymbol{X}}\boldsymbol{\beta})^\top \boldsymbol{\Omega}^{-1} \boldsymbol{\Pi}_r \boldsymbol{K}_l \boldsymbol{\Pi}_r \boldsymbol{\Omega}^{-1} (\boldsymbol{y} - \tilde{\boldsymbol{X}}\boldsymbol{\beta})$$

where $\boldsymbol{\Omega} = \sum_{r' \in Coloc(r)} \sigma_{r',m} \boldsymbol{\Pi}_{r'} \boldsymbol{K}_m \boldsymbol{\Pi}_{r'} + \tau \boldsymbol{I}$. Under the null hypothesis, the distribution of $S_{r,l}$ is a mixture of chi-square distributions [ZL03]. As in the global SVG test, we use the Satterthwaite approximation to represent it by a scaled chi-square distribution, $\kappa_{r,l}\chi^2(\nu_{r,l})$, where

$$\kappa_{r,l} = \kappa_{r,l}(\{\sigma_{r',m}; r' \in Coloc(r)\}, \tau) = \frac{tr(\{\boldsymbol{\Pi}_r \boldsymbol{K}_l \boldsymbol{\Pi}_r \boldsymbol{\Omega}^{-1}\}^2)}{tr(\boldsymbol{\Pi}_r \boldsymbol{K}_l \boldsymbol{\Pi}_r \boldsymbol{\Omega}^{-1})}$$
$$\text{and} \quad \nu_{r,l} = \nu_{r,l}(\{\sigma_{r',m}; r' \in Coloc(r)\}, \tau) = \frac{\big(tr(\boldsymbol{\Pi}_r \boldsymbol{K}_l \boldsymbol{\Pi}_r \boldsymbol{\Omega}^{-1})\big)^2}{tr(\{\boldsymbol{\Pi}_r \boldsymbol{K}_l \boldsymbol{\Pi}_r \boldsymbol{\Omega}^{-1}\}^2)}.$$

Since the parameters $\boldsymbol{\beta}$, $\tau$, and $\sigma_{r',m}$, are unknown, we estimate them under the null hypothesis using the REML estimator implemented in the `lmm.aireml` function of the R package `gaston`. Using the estimates $\hat{S}_{r,l}$, $\hat{\kappa}_{r,l}$, and $\hat{\nu}_{r,l}$ based on the REML estimates $\hat{\boldsymbol{\beta}}$, $\hat{\tau}$, and $\hat{\sigma}_{r',m}$, we compute the approximate p-value for each variance component as

$$\mathbb{P}\bigg( \frac{\hat{S}_{r,l}}{\hat{\kappa}_{r,l}} > \chi^2_{\hat{\nu}_{r,l}} \bigg).$$

After obtaining these p-values across all $l = 1, \ldots, L$, we combine them using the Cauchy combination method.

### 3 Why Spatial Coordinate Transformation Is Not Required

To explain why no spatial coordinate transformation is required for our analyses, it suffices to show that the kernel matrices used in the model remain unchanged under spatial coordinate transformation, since spatial coordinates are involved in the model only through the kernel matrices. Suppose the spatial coordinates are scaled as follows:

$$s'_{i,j} = \frac{s_{i,j} - \min_{a=1,\ldots,N} s_{a,j}}{\max_{j=1,2}\Big\{\max_{a=1,\ldots,N} s_{a,j} - \min_{a=1,\ldots,N} s_{a,j}\Big\}}$$

where $i = 1, \ldots, N$ and $j = 1, 2$. Denote $M := \max_{j=1,2}\big\{\max_{a=1,\ldots,N} s_{a,j} - \min_{a=1,\ldots,N} s_{a,j}\big\}$. The distance between the scaled $\boldsymbol{s}'_i$ and $\boldsymbol{s}'_k$ is then reduced by a factor of $M$ relative to the original distance between $\boldsymbol{s}_i$ and $\boldsymbol{s}_k$ as follows:

$$\|\boldsymbol{s}'_i - \boldsymbol{s}'_k\| = \sqrt{(s'_{i,1} - s'_{k,1})^2 + (s'_{i,2} - s'_{k,2})^2} = \sqrt{\left(\frac{s_{i,1} - s_{k,1}}{M}\right)^2 + \left(\frac{s_{i,2} - s_{k,2}}{M}\right)^2} = \frac{\|\boldsymbol{s}_i - \boldsymbol{s}_k\|}{M}.$$

We determine the lengthscale parameters of the kernel matrices as follows: We first pre-select $2L$ logarithmically spaced bandwidths ranging from the minimum to the maximum spatial distances across the $N$ locations. Since the minimum and maximum distances between the scaled coordinates are reduced by a factor of $M$ relative to the originals, the pre-selected bandwidths ($b_{l'}$) are likewise reduced by a factor of $M$ from the original bandwidths ($b_l$).

In the next step, we fit the global model with these candidate bandwidths using Non-Negative Least Squares (NNLS) and then identify the smallest and largest bandwidths with non-zero weights (denoted $l_{\min}$ and $l_{\max}$). The kernel used in NNLS is identical to the original kernel, as shown below:

$$[\boldsymbol{K}_{l'}]_{a,b} = \exp\left(-\frac{\|\boldsymbol{s}'_a - \boldsymbol{s}'_b\|}{b_{l'}}\right) = \exp\left(-\frac{M^{-1}\cdot\|\boldsymbol{s}_a - \boldsymbol{s}_b\|}{M^{-1}\cdot b_l}\right) = \exp\left(-\frac{\|\boldsymbol{s}_a - \boldsymbol{s}_b\|}{b_l}\right) = [\boldsymbol{K}_l]_{a,b}.$$

Therefore, the resulting parameter estimates remain the same. Consequently, the smallest and largest bandwidths with non-zero weights correspond to the original $l_{\min}$ and $l_{\max}$, reduced by a factor of $M$.

Finally, we define $L$ log-spaced bandwidths within $[l_{\min}, l_{\max}]$, which specify the final set of spatial bandwidths and the corresponding set of kernels. The final set of spatial bandwidths is also reduced by a factor of $M$ relative to the originals, and the kernels derived from them are identical to the original kernels, as shown above. Therefore, SpaceLink learns the same model regardless of spatial coordinate transformation.

### 4 Assessment of Result Robustness Under Model Assumption Violations in Mouse Organogenesis Spatiotemporal Data Analysis

The linear mixed model for the stage random effect in mouse organogenesis spatiotemporal data analysis failed normality assumption and homoscedasticity tests and showed low $R^2$. However, a clearly superior alternative is absent. Therefore, we verified robustness of results by applying one alternative, using logit-transformed ESV, in all downstream analyses. The main conclusions remained the same as follows:

- In whole tissue, genes with negative trend of logit(ESV) were significantly enriched in the muscle contraction pathway ($p = 0.001$), while genes with positive trend of logit(ESV) were significantly enriched in the Rab regulation of trafficking pathway ($p = 0.015$).
- In brain region, negatively ESV-associated genes were significantly enriched in the Wnt signaling pathway ($p = 0.002$), and positively ESV-associated genes were significantly enriched in the Rap1 signaling pathway ($p = 0.021$).

- Genes discussed in the main text (Myl9, Rab3a, Rspo2, Grin2b) retained significance in the same direction (Myl9: coefficient -0.44, p = 4e-4; Rab3a: coefficient 0.77, p = 0.045; Rspo2: coefficient -0.89, p = 3e-4; Grin2b: coefficient 0.85, p = 0.006).
- In brain region, genes with decreasing or increasing spatial variability across development exhibited logit(ESV) trends similar to the original ESV (**Supplementary Figure S13g**).
- Example genes in the main text showing different ESV trends between brain and non-brain regions also exhibited the same pattern with logit(ESV): Dlx6 and Dlx5 showed increasing logit(ESV) in brain (r = 0.88 and 0.77) but decreasing in non-brain (r = 0.88 and 0.85), Dmrt3 decreased in brain (r = 0.93) but increased in non-brain (r = 0.66), and Hoxd3 decreased in non-brain (r = 0.95) but showed no strong trend in brain (r = 0.42).
- Monogenic autism still exhibited a markedly stronger excess overlap with genes showing increasing spatial variability in brain, whereas genes associated with developmental Mendelian, monogenic psychiatric, or neurological disorders showed relatively weak overlap with genes displaying temporal trends in spatial variability in both brain and non-brain regions (**Supplementary Figure S13h**).

### 5 Alternative Approach: Hypothesis Testing Using Copula Distribution

We note that the hypothesis testing framework based on score test statistics in the previous sections does not account for dependencies among statistics from different kernels. One possible way to address these dependencies is to model the joint distribution of the score test statistics using a copula.

We illustrate this testing method by considering the model and hypotheses for the global SVG test in Section 2.1, and define the score vector as follows:

$$\boldsymbol{S} = \begin{pmatrix} \frac{1}{\tau}(\boldsymbol{y}-\boldsymbol{X\beta})^\top \boldsymbol{K}_1(\boldsymbol{y}-\boldsymbol{X\beta}) \\ \frac{1}{\tau}(\boldsymbol{y}-\boldsymbol{X\beta})^\top \boldsymbol{K}_2(\boldsymbol{y}-\boldsymbol{X\beta}) \\ \vdots \\ \frac{1}{\tau}(\boldsymbol{y}-\boldsymbol{X\beta})^\top \boldsymbol{K}_L(\boldsymbol{y}-\boldsymbol{X\beta}) \\ \frac{1}{\tau}(\boldsymbol{y}-\boldsymbol{X\beta})^\top(\boldsymbol{y}-\boldsymbol{X\beta}) \end{pmatrix} \in \mathbb{R}_+^{L+1}.$$

Under the null hypothesis, the marginal distribution of each $S_l$ $(l = 1, \ldots, L+1)$ can be derived as a mixture of chi-square distributions [ZL03], and the Satterthwaite approximation method is used to approximate each marginal distribution by a scaled chi-square distribution, $\kappa_l \chi^2(\nu_l)$, where:

$$\kappa_l = \frac{tr(\boldsymbol{K}_l^2)}{tr(\boldsymbol{K}_l)} \quad \text{and} \quad \nu_l = \frac{\left(tr(\boldsymbol{K}_l)\right)^2}{tr(\boldsymbol{K}_l^2)}.$$

Previous approach for hypothesis testing just relies on these marginal distributions and employed Cauchy combinations to combine multiple p-values. However, to account for dependencies among the statistics, we can instead leverage the joint distribution of the score vector $\boldsymbol{S}$, which directly captures these dependencies. According to equation (382) in [PP+08], the covariance of the score vector under $H_0$ is given by:

$$Cov(S_k, S_j) = tr(\boldsymbol{K}_k \boldsymbol{K}_j)$$

for $k, j = 1, \ldots, L+1$, where $\boldsymbol{K}_{L+1} = \boldsymbol{I}_N$. Hence, we can approximate the joint distribution by a $L+1$-dimensional multivariate distribution of scaled chi-square variables with covariance matrix $\boldsymbol{V}$, where $V_{k,j} = tr(\boldsymbol{K}_k \boldsymbol{K}_j)$.

However, deriving p-values directly from this multivariate distribution is computationally challenging. To overcome this, we can employ a copula method based on Sklar's theorem, which utilizes the marginal distributions and the correlation matrix of the target distribution. This method provides an efficient approach to derive the empirical distribution of the score vector. Detailed steps for the copula-based procedure are outlined in Algorithm 1.

---

**Algorithm 1:** Copula method to derive an empirical distribution

---

**Data:** $\{F_l\}_{l=1,\dots,L+1}$: marginal distributions of the target distribution, $\tilde{\boldsymbol{V}}$: correlation matrix of the target distribution, $n$: number of samples
**Result:** $\{\tilde{\boldsymbol{S}}_i\}_{i=1,\dots,n}$: samples to construct empirical distribution of the target distribution
**for** $i$ = 1,...,$n$ **do**
1. Sample $\boldsymbol{Z}_i$ from $\mathcal{N}(\boldsymbol{0}, \tilde{\boldsymbol{V}})$;
2. Calculate $U_{i,l} := F(Z_{i,l})$ for $l = 1, \dots, L+1$ where $F$ is the cdf of standard normal distribution.
   Let $\boldsymbol{U}_i = (U_{i,1}, \dots, U_{i,L+1})^\top$;
3. Obtain $\tilde{S}_{i,l} := F_l^{-1}(U_{i,l})$ for $l = 1, \dots, L+1$.
   Let $\tilde{\boldsymbol{S}}_i = (\tilde{S}_{i,1}, \dots, \tilde{S}_{i,L+1})^\top$;

**end**

---

Now, we consider the following $L$-dimensional vector of scaled score statistics:

$$\boldsymbol{S}^*(\boldsymbol{\beta}) = \begin{pmatrix} S_1/S_{L+1} \\ S_2/S_{L+1} \\ \vdots \\ S_L/S_{L+1} \end{pmatrix} = \begin{pmatrix} \frac{(\boldsymbol{y}-\boldsymbol{X\beta})^\top \boldsymbol{K}_1 (\boldsymbol{y}-\boldsymbol{X\beta})}{(\boldsymbol{y}-\boldsymbol{X\beta})^\top (\boldsymbol{y}-\boldsymbol{X\beta})} \\ \frac{(\boldsymbol{y}-\boldsymbol{X\beta})^\top \boldsymbol{K}_2 (\boldsymbol{y}-\boldsymbol{X\beta})}{(\boldsymbol{y}-\boldsymbol{X\beta})^\top (\boldsymbol{y}-\boldsymbol{X\beta})} \\ \vdots \\ \frac{(\boldsymbol{y}-\boldsymbol{X\beta})^\top \boldsymbol{K}_L (\boldsymbol{y}-\boldsymbol{X\beta})}{(\boldsymbol{y}-\boldsymbol{X\beta})^\top (\boldsymbol{y}-\boldsymbol{X\beta})} \end{pmatrix}.$$

The vector $\boldsymbol{S}^*(\boldsymbol{\beta})$ scales the components of the score vector $\boldsymbol{S}$ relative to $S_{L+1}$, thereby eliminating the influence of $\tau$.

We denote the marginal distributions of $\boldsymbol{S}^*(\boldsymbol{\beta})$ by $\{F_l^*\}_{l=1,\dots,L}$. As the test statistic, we consider the minimum of the p-values of the components of $\boldsymbol{S}^*(\boldsymbol{\beta})$, defined as:

$$p = \min_{l=1,\dots,L} \left\{1 - F_l^*\big(S_l^*(\boldsymbol{\beta})\big)\right\}.$$

Since $\boldsymbol{\beta}$ and $\{F_l^*\}_{l=1,\dots,L}$ are unknown, we estimate $\boldsymbol{\beta}$ by the MLE, $\hat{\boldsymbol{\beta}} = (\boldsymbol{X}^\top\boldsymbol{X})^{-1}\boldsymbol{X}^\top\boldsymbol{y}$, and each $F_l^*$ by its empirical distribution $\hat{F}_l^*$, obtained from the samples $\{\tilde{S}_{i,l}/\tilde{S}_{i,L+1}\}_{i=1,\dots,n}$ generated using Algorithm 1.

It is important to note that the resulting p-value does not follow a uniform distribution but rather corresponds to the minimum of correlated uniform variables. To account for this, we adjust the p-value based on the empirical distribution of the minimum of these variables. The correlation among the uniform variables is approximated by the correlation of the score vector, $\tilde{\boldsymbol{V}}$. Using Algorithm 1, we generate samples $\{\tilde{\boldsymbol{U}}_i\}_{i=1,\dots,n}$ of correlated uniform variables and derive the empirical distribution of their minimum, denoted as $F'$. The adjusted minimum p-value is then calculated as:

$$\textit{adjusted } p = F'\Big(\min_{l=1,\dots,L} \left\{1 - \hat{F}_l^*\big(S_l^*(\hat{\boldsymbol{\beta}})\big)\right\}\Big).$$

Finally, to control the false discovery rate (FDR), we apply the Benjamini-Hochberg (BH) procedure to the individual p-values $\left\{1 - \hat{F}_l^*\big(S_l^*(\hat{\boldsymbol{\beta}})\big)\right\}$ across genes, before computing the adjusted minimum p-value. If the adjusted minimum p-value is below the significance level $\alpha$, we reject the null hypothesis and conclude that the gene is spatially variable.

Even though the above approach directly accounts for dependencies among score test statistics, it did not perform better than the Cauchy combination approach in simulations. Supplementary Figure S16 presents the power and FDR of the two approaches across eight scDesign3-based simulations. The two approaches yield comparable results, although the Copula method tends to have higher power and FDR, and it fails to control FDR in one case. Because the Cauchy combination approach is more computationally efficient, we adopt it as our final method.

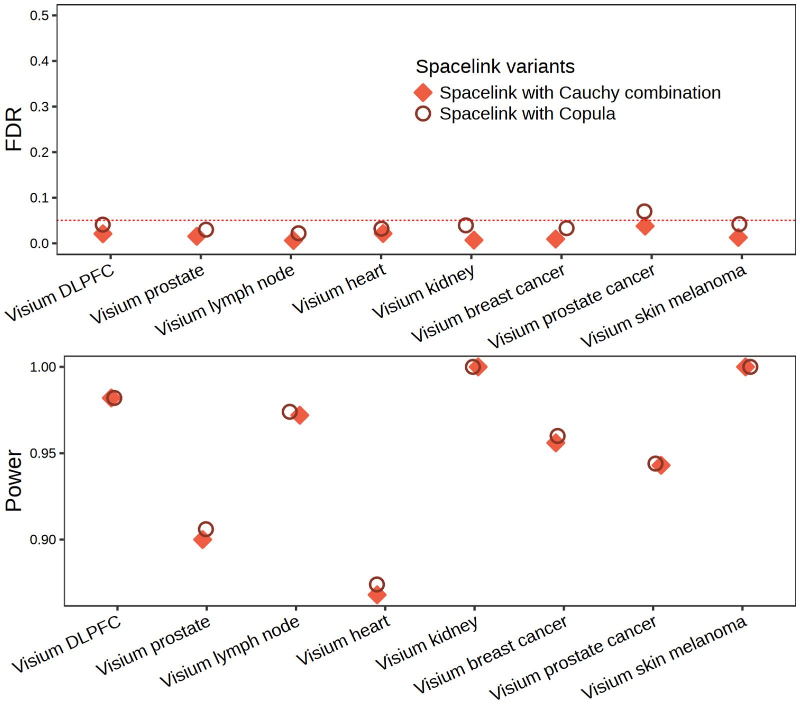


Supplementary Figure S16. False discovery rate and statistical power of Spacelink variants with Cauchy combination and Copula approaches across 8 scDesign3-based simulation datasets.
